# Sex- and age-specific development of ssRNA virus receptor expression in the human brain

**DOI:** 10.1101/2023.10.11.561925

**Authors:** Leanne Monteiro, Negeen Halabian, Jiawei Li, Maheshwar Panday, Nadine Barone, Peiyi Wang, Kathryn M. Murphy

## Abstract

Neurotropic single-stranded RNA (ssRNA) viruses can disrupt brain function, yet little is known about how host virus receptor expression develops across the human lifespan or whether these trajectories differ between females and males. We focused on postnatal development using postmortem transcriptomic data from 33 human donors (4 months–82 years; both sexes), comprising 52 cerebral hemispheres and 15 brain areas, to characterize developmental patterns of 67 host receptor genes that mediate entry of major ssRNA viral families. Using sex-specific LOESS trajectory modelling, hierarchical clustering, and sliding-window differential expression analyses, we identified multiple non-linear developmental programs governing virus receptor expression. About half of female–male gene–area trajectory pairs exhibited divergent developmental patterns. Sex differences were most pronounced in the cortex, where high-dimensional receptor expression profiles were sufficient to predict the sex of individual cases. Sex differential expression was most frequent during early childhood, coinciding with sensitive periods of cortical circuit refinement. A targeted prenatal analysis revealed that virus receptor expression was predominantly male-biased before birth, contrasting with the predominantly female-biased expression observed during the early postnatal period. Although receptors from most viral families were broadly distributed across developmental programs, a subset of sex-by-age expression clusters showed some enrichment for specific viral lineages and glial-associated cell-type signatures. Together, these findings reveal that virus receptor expression in the human brain is organized into structured, sex-biased trajectories that dynamically reorganize over development, providing the first developmental atlas of viral entry receptors in the human brain.

**Significance Statement:** Many viruses that affect the brain use host receptors to enter cells, yet it is unclear how these receptors are expressed across the human lifespan or whether their expression differs between females and males. Using human postmortem brain transcriptomic data spanning infancy to older adulthood, we show that virus receptor genes follow dynamic, sex-biased developmental trajectories, particularly in the cortex. These patterns are strongest during early childhood and are sufficient to distinguish female and male expression profiles at the individual level. Rather than acting as static entry points, virus receptors are embedded within developmental and neuroimmune programs. These findings provide a developmental framework for understanding why vulnerability to neuroinfectious disease may vary by age, brain area, and sex.

## Introduction

The COVID-19 pandemic, widespread cases of West Nile Virus (WNV), the Zika virus outbreak, and rising Dengue fever cases highlight the growing public health threat posed by neuroinfectious diseases caused by single-stranded RNA (ssRNA) viruses [1–4]. These viruses can impact the brain through direct or indirect routes and trigger neurological symptoms ranging from headache, anosmia, and confusion to encephalitis, stroke, and death [5–8]. Beyond acute symptoms, viral infection and the associated immune response can disrupt neuronal and glial function, alter neuroimmune signalling, and induce structural and transcriptomic changes in the cortex that are associated with neurodevelopmental, neuropsychiatric, and neurodegenerative disorders [9–16].

The intensity, persistence, and fatality of these outcomes vary widely among individuals and are strongly influenced by host factors such as sex and age [17–21]. For example, studies have found that men and older adults are more likely to experience greater clinical severity with COVID-19, SARS, and MERS [20,22,23], and have a greater risk of neuroinvasive disease with WNV [24,25]. In contrast, women show higher morbidity and mortality from influenza and greater persistence of post-acute sequelae such as long COVID [19,26–28] with cognitive impairment that often persists for more than a year [29]. In childhood, several sex differences have been reported, including males having higher odds of severe disease with SARS-CoV-2 infection [30], a higher incidence of viral meningitis [31–33] and a higher risk for neuroinvasive WNV [25]. These sex- and age-related differences highlight the importance of host biology in shaping the severity and outcomes of these viral infections.

Despite this, little is known about how virus receptor expression in the human brain varies by sex and age. Viral pathogens can invade the brain via various routes, and their receptors mediate entry into neurons, glia and other brain cell-types [8]. This gap is significant given evidence that severe infections can leave behind structural and transcriptomic signatures resembling brain aging [34,35]. Furthermore, understanding the cellular context of virus receptor expression is important because it dictates how a virus interacts with the brain’s architecture and which pathological outcomes may follow. Comparing the development of virus receptors between the sexes, across brain areas, within cell-type associations, and across the lifespan may provide insights into sex biases associated with neurological infections.

Here, we use transcriptomic data to map lifespan trajectories of 67 receptors for ssRNA viruses in the human brain in females and males. Our primary analyses focus on the postnatal lifespan, characterizing how virus receptor expression is organized from infancy through aging as the brain undergoes experience-dependent plasticity, prolonged maturation, and age-related remodelling. We also performed a targeted prenatal analysis to determine whether sex differences in virus receptor expression are already established during corticogenesis. We implemented a data-driven computational workflow that integrates sex as a biological variable, enabling the identification of robust sex- and age-related differences in receptor expression. The approach captures both linear and non-linear developmental patterns and uses high-dimensional analytical tools to discover the range of patterns and sex differences in virus receptor expression in the human brain across the lifespan. We use a machine-learning approach to determine whether population-level developmental structures encode sufficient information to accurately predict the sex of individual cases. Finally, we explore the association of the developmental patterns with virus families and cell-types.

## Methods

### Human transcriptomic dataset

The Hamilton Integrated Research Ethics Board reviewed and approved the use of the dataset in this study. We analyzed postnatal human brain transcriptomic data from Kang et al. (2011)(GSE25219), as it is the most comprehensive publicly available dataset covering the human lifespan. This dataset includes exon-level gene expression data from multiple brain areas in both sexes and across diverse ethnic backgrounds. The original study conducted rigorous quality assessments (PMI, pH, RIN, DS) and reported no significant correlations between these measures and expression levels. Details on tissue acquisition, processing, and validation are available in Kang et al. (2011) and its Supplementary Material.

The study focused on postnatal development using data from 15 brain areas with lifespan sampling for both sexes: 11 cortical areas (OFC, DFC, VFC, MFC, M1C, S1C, IPC, A1C, STC, ITC, V1C) and 4 subcortical areas (HIP, AMY, MD, CBC)(total tissue samples: n = 700; 306 females, 394 males)(Fig. 1A). The postnatal dataset comprised 33 postnatal donors (aged 4 months to 82 years; 14 female). Because 18 donors contributed both cerebral hemispheres, the final dataset contained 52 hemispheres (Fig. 1A; Supplementary Table 1). Each hemisphere was treated as a case, with multiple brain areas sampled per hemisphere. Although areas from the same hemisphere are biologically related, each tissue sample was processed and quantified independently. Developmental trajectories were modelled separately for each gene and brain area, and sex differences were assessed at the trajectory level, thereby minimizing repeated measurement of the same tissue sample (Fig. 1B).

**Figure 1.**
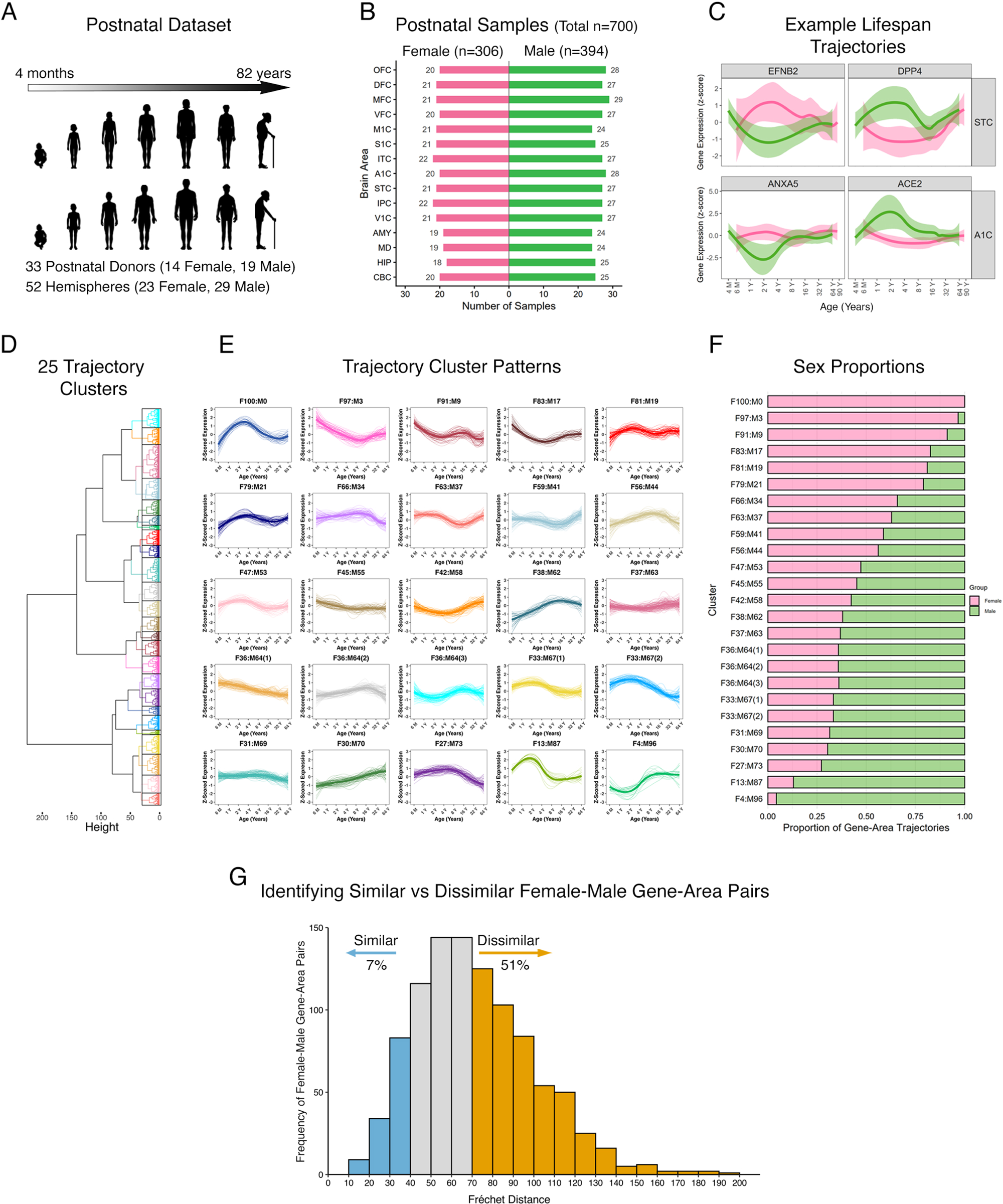
Postnatal developmental trajectories of ssRNA virus receptor genes in the human brain reveal sex-specific patterns. (A) Summary of the postnatal dataset, including age range, number of donors, and hemispheres (cases) by sex. (B) Postnatal sample composition across the 15 brain areas, showing the number of samples by sex (total samples: n = 700; 306 females, 394 males). (C) Representative examples of postnatal lifespan trajectories generated by locally estimated scatterplot smoothing (LOESS) for individual gene–area combinations, fitted separately for females (magenta) and males (green). (D) Unsupervised hierarchical clustering of 2010 gene–area–sex trajectories (67 genes × 15 brain areas × 2 sexes) identified 25 distinct developmental patterns. (E) Lifespan trajectories for each cluster, showing individual trajectories and the cluster-average trajectory, with clusters labelled based on the proportion of female and male trajectories (e.g., F100:M0, F4:M96). (F) Stacked bar plots showing the proportion of female (magenta) and male (green) trajectories within each cluster. (G) Distribution of Fréchet distances for female–male gene–area trajectory pairs, with thresholds derived from within-cluster variation identifying 7% of pairs as strongly similar (teal) and 51% as strongly dissimilar (orange), and intermediate values (grey) representing trajectories within the range of expected variation. Human development images in panel A were adapted from Servier Medical Art (https://smart.servier.com), licensed under CC BY 4.0 (https://creativecommons.org/licenses/by/4.0/).

For the targeted prenatal analyses, we used cortical transcriptomic data from the prenatal portion of the Kang et al. (2011) dataset (13–37 post-conception weeks). The prenatal dataset comprised 17 donors contributing 304 cortical tissue samples (162 female, 142 male)(Supplementary Table 1). Because the prenatal dataset does not include complete sampling of all cortical regions represented in the postnatal dataset, prenatal analyses were performed using cortical tissue collapsed across areas and were analyzed separately from the postnatal lifespan trajectories.

We retrieved the processed and normalized exon-array expression data from Gene Expression Omnibus (GSE25219) and reformatted it into a working file for this study (see Data availability statement). Probe identifiers were annotated using the Affymetrix Human Exon 1.0 ST array annotation file, and probes lacking gene annotation (n = 37) were removed. The full exon-array dataset initially contained genes represented by multiple probes (285 genes with two probes and 3 genes with three probes). Probe-level expression values were retained because different probes may target distinct exons or transcript variants. However, after filtering to the 67 virus receptor genes analyzed in the present study, each gene was represented by a single probe. Therefore, no probe selection or collapsing was required for the analyses presented here. Information on case characteristics, brain area distribution, and causes of death is provided in Supplementary Table 1. In this study, “sex” refers to the tissue’s biological characteristics, as donor gender was not recorded. None of the individuals included in our study died from a viral infection.

### Analysis of virus receptor lifespan trajectories

We studied lifespan changes in the expression of 67 host receptor genes that mediate entry of ssRNA viruses into human cells. Receptor genes were identified using a curated virus receptor database (http://computationalbiology.cn:5000/viralReceptor/type?type=4) [36]. These 67 receptors represent 10 viral families: *Arenaviridae*, *Coronaviridae, Filoviridae, Flaviviridae, Orthomyxoviridae, Paramyxoviridae, Picornaviridae, Pneumoviridae, Rhabdoviridae, and Togaviridae* (see Supplementary Table 2).

For each virus receptor gene, we examined developmental trajectories by z-scoring the expression values within each brain area and fitting LOESS curves (locally estimated scatterplot smoothing) to capture both linear and non-linear trends without assuming a predefined model. Lifespan trajectories were fitted separately for females and males using the loess() function from the *stats* package in R [37] with a quadratic local polynomial (degree = 2), Gaussian family, and span = 0.75 (Fig. 1C). A quadratic local polynomial (degree = 2) was used because developmental trajectories frequently exhibit changes in curvature that cannot be captured by a locally linear fit, while avoiding the greater instability associated with higher-order polynomials. After fitting the LOESS curves, 128 evenly spaced points spanning the overlapping age range of the female and male trajectories were extracted to provide a standardized representation of each developmental trajectory for downstream distance-based and clustering analyses. This resolution preserved the overall trajectory shape while maintaining computational efficiency. Age was plotted on a log₂ scale to facilitate visualization of rapid developmental changes in early life versus slower changes in adulthood. Confidence intervals (95%) were derived from the standard errors of the LOESS fits. In total, 2010 trajectories were generated (67 receptor genes × 15 brain areas × 2 sexes) (Supplementary Fig. 1). Some donors contributed tissue from both cerebral hemispheres (Supplementary Table 1), resulting in limited biological nesting within the dataset. Because LOESS was used to generate descriptive developmental trajectories rather than for coefficient-based statistical inference, donor identity was not incorporated into the smoothing procedure. The stability of the fitted trajectories was evaluated using repeated leave-one-case-out resampling (see below).

The trajectories were hierarchically clustered using Ward.D2 linkage with Euclidean distance computed on the 128 LOESS-fitted points. The number of clusters was determined using the elbow method, based on the total within-cluster sum of squares plotted against k, with the optimal k corresponding to the inflection point of the curve (Supplementary Fig. 3).

Clusters were labelled based on the ratio of female to male trajectories. For example, a cluster labelled F100:M0 contained only female trajectories. The uniqueness of the cluster trajectories was validated using the Fréchet distance metric to compare trajectory shapes while preserving the orderly progression of age [38]. The Fréchet distance is 0 for identical trajectories and increases as the difference between trajectories increases. Because Fréchet distances were used as descriptive measures of trajectory similarity rather than individual statistical tests, inference was performed only on the resulting distance distributions. Statistical inference was based on estimation statistics rather than null-hypothesis significance testing. Differences between Fréchet distance distributions were quantified using Hodges–Lehmann effect size estimates with 95% confidence intervals [39,40]. Between-cluster differences were summarized using effect size estimates of the median difference with bootstrapped confidence intervals (Supplementary Fig. 4). Within-cluster distances (Supplementary Fig. 5A), as well as direct contrasts between within- and between-cluster distance distributions, were quantified using the Hodges–Lehmann estimator, a robust, non-parametric estimator of the median difference with analytic confidence intervals. In all cases, within-cluster distances were smaller than between-cluster distances, validating the unique patterns of virus receptor development identified by the trajectory clusters (Supplementary Fig. 6).

To evaluate the robustness of the LOESS trajectories and the stability of sex differences, we performed repeated leave-one-case-out resampling on a random subset of 100 gene–area combinations. Combinations were selected by uniform random sampling without replacement from the full set. For each gene-area combination, we refit the corresponding female and male LOESS trajectories (gene–area–sex) across 10,000 iterations, each time randomly omitting one female and one male case. In each iteration, we recalculated the Fréchet distance between the refitted female and male trajectories, generating a distribution of simulated distances from which we computed the mean for each combination. Estimation statistics were used to compare the original Fréchet distance with the mean of the simulated distribution by calculating the unpaired median difference and its 95% confidence interval. This approach allowed us to test whether the observed sex differences remained stable when individual cases were removed.

### Analyzing the high-dimensional pattern of virus receptor development among brain areas

To identify brain areas for females and males with similar developmental profiles, we applied hierarchical clustering to brain area–sex LOESS trajectories (15 brain areas × 2 sexes). To visualize the structure in this high-dimensional data, we then applied den-SNE, a density-preserving modification of t-distributed stochastic neighbour embedding (t-SNE), implemented in the densvis package (perplexity = 30, dens_frac = 0.9, dens_lambda = 0.1)[41]. This analysis projects high-dimensional data into a two-dimensional (2D) space while preserving both local and global structure. Each point in the embedding represented an LOESS-fitted point for a given brain area and sex, encoded by its 67-gene expression vector. den-SNE was run using multiple random seeds to confirm stability. Cluster labels derived from hierarchical clustering were overlaid onto the den-SNE embedding, and points were additionally colour-coded by age or sex. Subsequent analyses were performed on the cortical trajectories identified in the preceding hierarchical clustering analysis.

### Random Forest model

Random forest classification was used to assess whether population-level developmental structure in virus receptor expression could predict the sex of individual cases. To train the model, we implemented a leave-one-case-out approach using sex-specific developmental information derived from LOESS-fitted trajectories for the 67 virus receptor genes in females and males. For each iteration, one case was removed from the dataset, population-level LOESS curves were refitted without that case, and the female and male trajectories were each sampled at the same 128 evenly spaced ages spanning their overlapping age range. Each sampled age constituted one training observation, and the corresponding LOESS-derived expression values for the 737 gene–area combinations (67 genes x 11 cortical areas) formed the feature vector, yielding a training dataset of 256 observations (128 female and 128 male) × 737 features. The random forest classifier was coded in R using the randomForest() function from the *randomForest* package [42], and the model was trained 10 times using different random seeds to account for variability in model initialization and training, and mean classification accuracy across runs was used as the summary performance metric (see code for implementation).

The trained model generated a predicted probability of female classification. Sex was treated as a binary outcome variable, with females coded as the positive class. A probability threshold of 0.5 was used to assign class labels (>0.5 classified as female, <0.5 classified as male). Model outputs were visualized by plotting the predicted probability of female classification as a function of age, allowing assessment of how population-level developmental structure relates to individual-level sex prediction across the lifespan.

### Differential expression-sliding window analysis (DE-SWAN)

To identify virus receptor genes showing sex differences across the lifespan, we applied Differential Expression–Sliding Window Analysis (DE-SWAN) [43] to the LOESS-fitted expression trajectories. Sliding windows were defined over adjacent LOESS-fitted points in a data-driven manner [43]. Robustness of the windows was evaluated by running 3 sizes (±4, ±10, and ±20 LOESS points) and multiple significance thresholds (p = 0.001, 0.01, 0.05). P values were adjusted using the Benjamini–Hochberg procedure to generate q values (Supplementary Fig. 8). At each window center, the number of significantly differentially expressed genes (q < 0.05) was counted and plotted. Inflection points in this curve (peaks and troughs) were used to define the reference ages for downstream analysis, which were then used to generate the DE-SWAN heatmap. Twelve windows were identified, centered at the following reference ages: 1, 1.5, 2, 3, 5, 8, 12, 16, 24, 32, 45, and 60 years (Supplementary Fig. 8). Sex differential expression was visualized as heatmaps, with rows representing genes, columns representing reference ages, and colours indicating the direction of expression between the sexes (magenta: female > male; green: male > female). The rows were organized using hierarchical clustering to generate DE-SWAN clusters that identified groups of genes with similar sex-differential expression patterns.

### Virus family enrichment across trajectory clusters

To assess whether virus families were enriched in the developmental patterns, we compared the virus family composition of each DE-SWAN cluster with the global composition using Jensen–Shannon (JS) divergence. JS divergence was used because it provides a symmetric, bounded measure of dissimilarity between multi-category compositions and does not rely on asymptotic assumptions about count distributions. DE-SWAN clusters comprised gene–area trajectories, and virus family annotations were assigned at the gene level.

Statistical significance was evaluated by permuting virus family labels across all gene–area items (10,000 permutations), thereby preserving cluster sizes and the global family frequency structure, to generate an empirical null distribution of JS divergence for each cluster. Observed JS divergence values were standardized relative to the permutation-derived null to obtain scores in SD units. Values of approximately 1, 2, and ≥3 SD were interpreted as modest, strong, and very strong enrichment, respectively. Clusters with strong to very strong divergence (SD ≥ 2) from the global distribution were identified with an asterisk.

To identify which virus families contributed to the observed compositional differences within JS-enriched clusters, family-level enrichment was assessed using permutation testing restricted to clusters with SD ≥ 2. For each cluster–family pair, observed family counts were compared to a permutation-derived null distribution generated by randomly reassigning family labels across genes while preserving cluster size and global family frequencies. One-sided permutation p-values were computed to test for enrichment, and were corrected for multiple comparisons using the Benjamini–Hochberg FDR.

### Cell-type fidelity analysis

To relate trajectory clusters to putative cell-type programs, we quantified the representation of major brain cell-type signatures within each trajectory. Genome-wide estimates of expression fidelity for astrocytes, neurons, microglia, and oligodendrocytes were obtained from human frontal, occipital, parietal, and temporal cortex reference datasets (Kelley et al. 2018, http://oldhamlab.ctec.ucsf.edu/). Fidelity scores were extracted for the cortical area corresponding to each trajectory: frontal (DFC, OFC, MFC, VFC), occipital (V1C), parietal (IPC, M1C, S1C), and temporal (ITC, A1C, STC). All 67 genes associated with the trajectories were present in the reference dataset.

Each trajectory (gene × cortical area) contributed Fidelity scores to all cell-type categories, allowing non-exclusive assignment to multiple cell-types. For each trajectory cluster, the reference distribution was defined as the Fidelity scores of all trajectories within that cluster. For each cell-type, we then computed the unpaired median difference between its Fidelity score distribution and this reference distribution (Supplementary Fig. 9). A 95% confidence interval excluding zero indicated a significant deviation from the reference, with positive median differences indicating over-representation and negative differences indicating under-representation.

## Results

### Sex-biased lifespan trajectories of virus receptor expression

We characterized the range of lifespan trajectory patterns for virus receptor expression in the human brain and assessed how these patterns are differentially represented in females and males. To do this, we used a data-driven approach that avoids age-based binning of samples. Instead, we fit LOESS curves separately for females and males for each virus receptor gene in each brain area, generating continuous trajectories that capture gradual, non-linear changes in expression across development. This approach yielded 2010 trajectories (67 receptor genes × 15 brain areas × 2 sexes), with each trajectory representing the developmental profile of a single virus receptor gene in a specific brain area (gene-area trajectory) for one sex (Fig. 1C, Supplementary Fig. 1).

Unsupervised hierarchical clustering of all 2010 trajectories identified 25 trajectory patterns (Fig. 1D,E), and the trajectories within a cluster were more similar than between clusters (Supplementary Fig. 6). These analyses validated the presence of multiple developmental programs governing virus receptor expression in the human brain. While some trajectories showed predominantly monotonic changes across the lifespan, including monotonic increases (e.g., F30:M70) or decreases (e.g., F36:M64(1)), the majority were characterized by non-linear, often undulating trajectories, with peaks and troughs occurring at different developmental stages (e.g., F100:M0, F13:M87; Fig. 1E).

Importantly, these developmental patterns exhibited a range of sex biases. Clusters differed widely in their female-to-male composition (Fig. 1F), with several clusters exclusively or strongly biased toward females (e.g., F100:M0, F97:M3) or toward males (e.g., F13:M87, F4:M96), and others containing trajectories from females and males in roughly equal proportions (e.g., F47:M53). Inspection of clusters containing both sexes indicated that most trajectories differed across gene–brain area combinations in females and males (Supplementary Table 3), suggesting that females and males preferentially allocate different gene–area combinations to shared trajectory shapes.

To directly assess the degree of similarity versus dissimilarity between female and male trajectories for the same virus receptor gene and brain area, we defined an empirical threshold for developmental similarity and then compared that with the Fréchet distance for each female-male gene pair. Developmental similarity was quantified as the Fréchet distance among trajectories within each of the 25 clusters. Within-cluster distances represent trajectories with similar lifespan patterns and thus define the expected range of developmental similarity (Supplementary Fig. 5). Fréchet distances equal to or less than the within cluster median were defined as developmentally similar (Fréchet distances =< 35.27, bootstrap 95% CI: 35.16–35.40), and values larger than the upper 97.5th percentile were defined as dissimilar (Fréchet distance => 67.5). Applying this criterion, we found that only 7% of gene-area trajectory pairs showed similar developmental patterns, whereas 51% exhibited dissimilar developmental patterns between females and males (Fig. 1G).

To validate the comparison of female and male trajectory patterns, we ran a leave-one-case-out simulation to test whether individual cases influenced the sex differences. The leave-one-case-out simulation was run 10,000 times, and the Fréchet distances were averaged across simulations for each gene–area combination. The resulting distribution did not differ from the original distribution of Fréchet distances (Supplementary Fig. 2). This supports the robustness of the trajectory approach for quantifying sex differences in the lifespan patterns of virus receptor expression in the human brain.

Together, these findings indicate that sex differences are not limited to shifts in the magnitude or timing of expression, but instead reflect sex-biased trajectory patterns for virus receptors in the human brain.

### Sex differences in virus receptor development are most pronounced in the cortex

To determine whether specific brain areas underlie the sex biases identified by trajectory clustering, we conducted a brain–area–by–sex trajectory analysis. For this analysis, trajectories were reorganized into a high-dimensional matrix with 30 rows, corresponding to each brain area and sex (15 brain areas × 2 sexes), and 8,576 columns representing the 128 LOESS-fitted values for each of the 67 virus receptor genes (128 values × 67 genes). Unsupervised clustering of this matrix identified five groups of brain areas (Fig. 2A).

**Figure 2.**
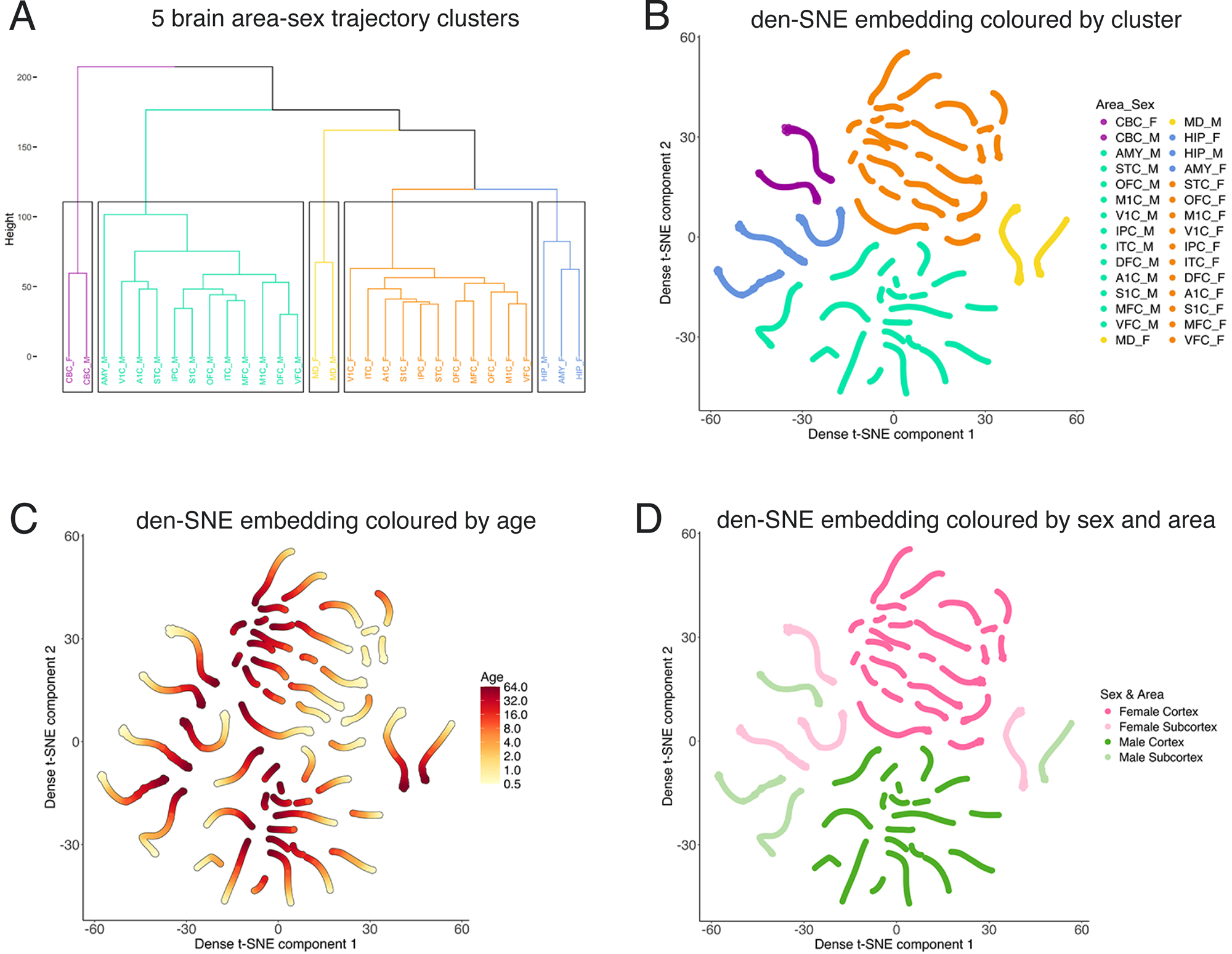
Cortical brain areas show pronounced sex-specific organization of virus receptor development. (A) Unsupervised hierarchical clustering of brain-area-by-sex trajectories (15 brain areas × 2 sexes) identified 5 distinct clusters. (B–D) Density-preserving t-distributed stochastic neighbour embedding (den-SNE) of the same high-dimensional trajectories illustrates relationships among brain areas across the lifespan. Plots are coloured by (B) cluster assignment from hierarchical clustering (k = 5), (C) age (lighter colours representing younger ages and progressively darker shades indicating older ages), and (D) sex (female, magenta; male, green) and brain region, with saturated colours indicating cortical areas and unsaturated colours indicating subcortical areas. Cortical trajectories segregate strongly by sex, whereas subcortical trajectories show greater overlap between females and males. Brain area abbreviations: A1C, primary auditory cortex; AMY, amygdala; CBC, cerebellar cortex; DFC, dorsolateral prefrontal cortex; HIP, hippocampus; IPC, posterior inferior parietal cortex; ITC, inferior temporal cortex; M1C, primary motor cortex; MFC, medial prefrontal cortex; OFC, orbital prefrontal cortex; S1C, primary somatosensory cortex; STC, superior temporal cortex; V1C, primary visual cortex; VFC, ventrolateral prefrontal cortex; MD, mediodorsal thalamus.

To visualize relationships among brain areas in this high-dimensional space, we applied density-preserving t-distributed stochastic neighbour embedding (den-SNE)[41] (Fig. 2B). When colour-coded by age (from early life, light yellow, to older adulthood, dark red), all clusters contained trajectories spanning the whole lifespan (Fig. 2C), indicating that age alone did not account for the observed clustering structure.

In contrast, colour-coding by sex (female, magenta; male, green) and by cortical (saturated) versus subcortical (unsaturated) brain areas revealed a clear separation among the clusters. The 11 cortical areas formed separate female- and male-specific clusters, whereas subcortical areas contained both sexes (Fig. 2D, Supplementary Fig. 7). This pattern indicates that sex-specific developmental programs of virus receptor expression are most pronounced in the cortex, while subcortical regions show more similar developmental profiles between females and males. Because sex-specific developmental organization was observed only in the cortical regions, all subsequent analyses (DE-SWAN, Random Forest classification, virus-family enrichment, and cell-type fidelity analysis) were restricted to the cortex.

### Cortical virus receptor expression predicts sex at the individual level

The preceding analyses identified sex-specific, population-level cortical development of virus receptor expression. We next asked whether these developmental programs were sufficiently robust to generalize to individual brains. Specifically, we tested whether a Random Forest (RF) classifier trained on population-level, LOESS-derived developmental trajectories could correctly classify the sex of individual cases based on their observed cortical virus receptor expression profiles. Training on LOESS-derived trajectories allowed the model to learn the underlying sex-specific developmental programs while minimizing the influence of individual-level variability. Model performance was then evaluated by testing its ability to classify the sex of individual cases using their observed expression profiles.

The RF model achieved an overall classification accuracy of 82% (SD = 3.85%, CI = 80.8%–83.2%) for sex prediction. All female cases were correctly classified, whereas only 5 male cases were misclassified, including 4 of 14 in the adolescent-to-adult age range (Fig. 3). LOESS curves fitted to the predicted probabilities showed clear separation between females and males across the lifespan, with female predictions above and male predictions below the decision boundary. These results indicate that sex differences in cortical virus receptor expression are strong and systematic across the lifespan, reflecting reproducible developmental organization rather than random variation.

**Figure 3.**
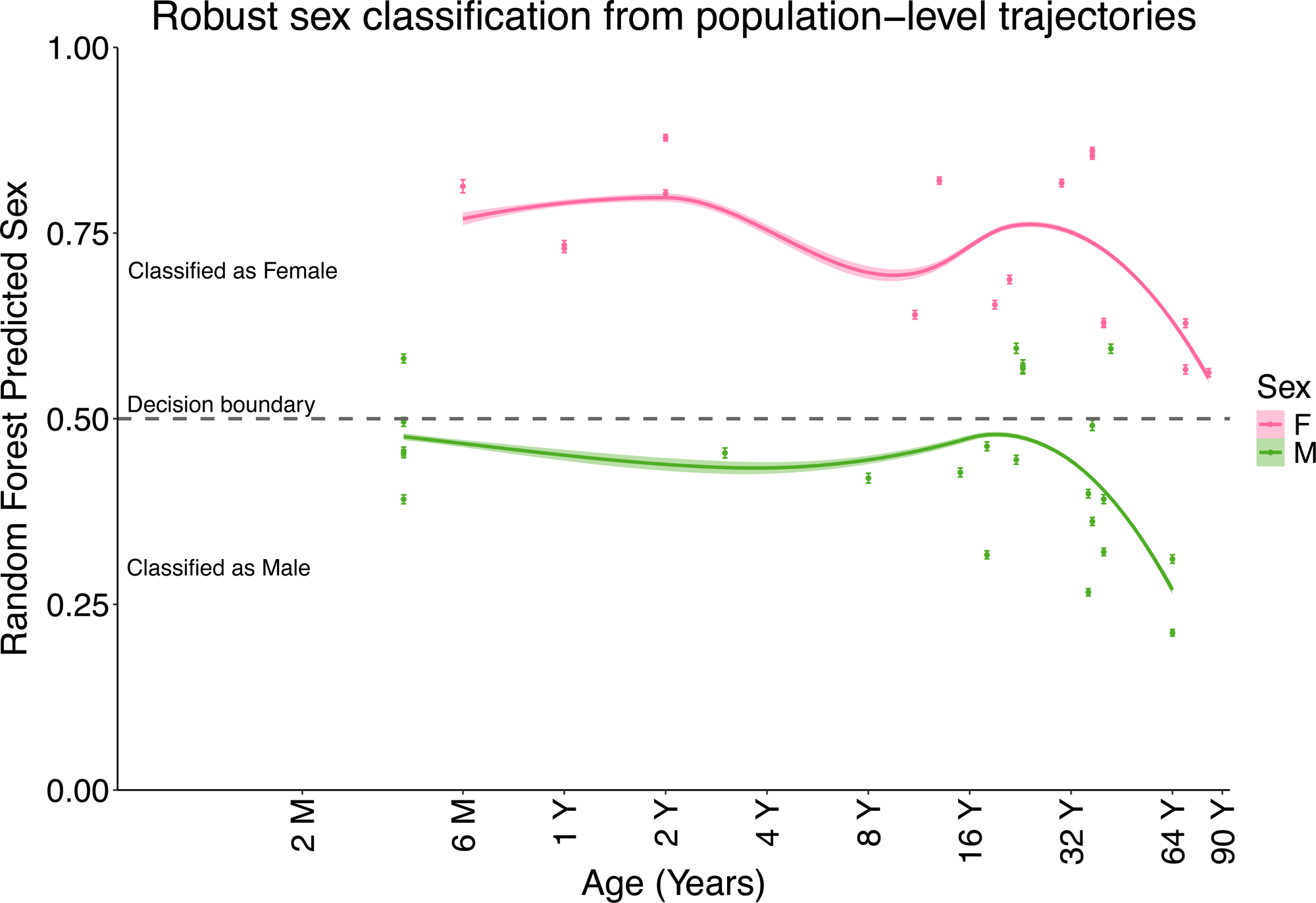
Robust classification of sex from population-level developmental trajectories. Random Forest classification was performed using LOESS-fitted developmental trajectories of 67 virus receptor genes across 11 cortical areas to predict the sex of individual cases. A leave-one-case-out approach was implemented in which, for each held-out case, LOESS trajectories were refit excluding that case, and the model was retrained ten times. Each point represents a single case and shows the mean predicted probability of being female (magenta, female cases; green, male cases), with error bars indicating 95% confidence intervals. A probability threshold of 0.5 was used for classification (>0.5: female; ≤0.5: male). LOESS-smoothed curves illustrate age-related trends in predicted probability for each sex. The female curve remained above and the male curve below the decision boundary across the lifespan. No female cases were misclassified, whereas 5 male cases were misclassified, including 4 of 14 cases in the adolescent-to-adult range. Overall classification accuracy was 82%.

### Dynamic sex-by-age patterns of virus receptor expression in the cortex

To determine when sex differences in cortical virus receptor expression emerge across development and whether the various virus families are associated with distinct sex-specific patterns, we applied Differential Expression–Sliding Window Analysis (DE-SWAN)[43] to the cortical data (11 cortical areas × 67 virus receptor genes). This analysis compares female and male expression for the same gene within overlapping age windows across the lifespan, allowing identification of age-specific periods of differential expression.

DE-SWAN revealed sex differences in virus receptor expression throughout the lifespan, with approximately equal instances of higher expression in females (magenta) and males (green) (Fig. 4A). However, sex differences were most frequent during early childhood, when approximately half of the cortical trajectories showed significant differential expression between females and males.

**Figure 4.**
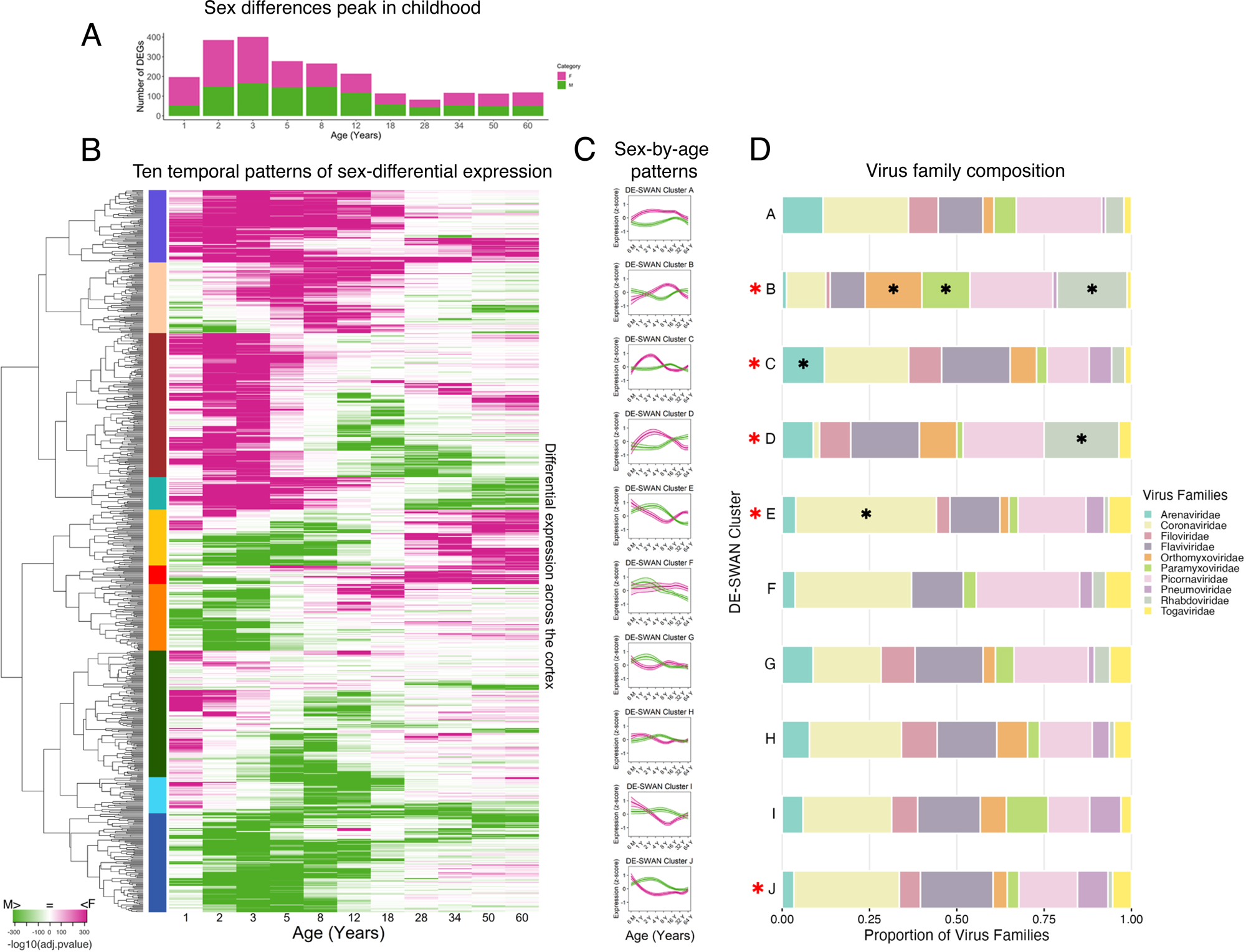
Sex-by-age differential virus receptor expression (DE-SWAN) in the cortex peaks during early childhood. (A) Stacked bar plot showing the number of sex-differentially expressed genes at each reference age (magenta, higher in females; green, higher in males), with a peak in early childhood (2–3 years) when more than half of the genes show differential expression. (B) Differential Expression–Sliding Window Analysis (DE-SWAN) heatmap illustrating the ages at which females (magenta) or males (green) show higher expression of virus receptor genes. Rows represent gene–brain area combinations across the 11 cortical areas, and columns correspond to reference ages. Hierarchical clustering identifies 10 distinct clusters of sex-by-age expression patterns. (C) LOESS trajectories for females (magenta) and males (green) for each DE-SWAN cluster, illustrating the distinct lifespan expression profiles underlying the clustered patterns. (D) Stacked bar plots showing the relative contribution of virus families within each cluster. Clusters with virus family compositions that differ significantly from the global distribution were identified using Jensen–Shannon divergence (SD > 2; red asterisk), and significantly enriched virus families within clusters are indicated with a black asterisk.

The DE-SWAN results were visualized as a heatmap (Fig. 4B), with hierarchical clustering grouping virus receptor genes that shared similar sex- and age-related differential expression patterns. This analysis identified 10 distinct DE-SWAN clusters, representing different temporal patterns of sex bias across the lifespan (Fig. 4C, Supplementary Table 4). While some clusters showed consistently higher expression in one sex, such as higher expression in females (Cluster A) or males (Cluster J), most clusters exhibited dynamic patterns in which the sex with higher expression changed across development. For example, Cluster D switched from females higher in childhood to males higher in adulthood. In contrast, Cluster E was the opposite: males were higher in childhood, and females were higher in adulthood.

To identify which virus families align with these developmental programs, we examined the representation of virus families within each DE-SWAN cluster. Overall, virus families were broadly distributed across clusters, indicating that sex-by-age expression patterns are not exclusive to individual families (Fig. 4D). However, Jensen–Shannon enrichment analysis identified five clusters (B, C, D, E, and J) with strong (>2 SD) to very strong (>3 SD) enrichment for specific virus families. Notably, Clusters B and D, despite exhibiting different sex-by-age expression patterns, were both enriched for Rhabdoviridae receptors, whereas Cluster E was selectively enriched for Coronaviridae receptors. These results indicate that while sex-specific developmental patterns of viral receptor expression are broadly conserved across virus families, a subset of temporal expression programs shows selective alignment with specific viral lineages.

### Selective cell-type associations characterize a subset of DE-SWAN clusters

Previous studies have identified a diverse range of virus-host cell interactions in the brain. For example, glial interactions include the production of pro-inflammatory cytokines, leading to neuroinflammation, or may play a more important role in infection, such as astrocytes in ZIKA [44]. Neuronal interactions include cell lysis and synapse dysfunction, and certain viruses, such as RABV, have stronger associations with neurons, leading to GABA dysregulation [45] and molecular changes that impact neuronal function [46].

To examine the cellular context in which the various DE-SWAN clusters are most likely to be exposed to, or responsive to, we inferred cell-type associations for the virus receptor genes using Fidelity scores [47]. We used the Fidelity score because it quantifies the relative specificity and enrichment of gene expression across the 4 major cortical cell-types (astrocytes, microglia, neurons, oligodendrocytes), enabling us to infer which cell populations are most strongly associated with virus receptor genes independent of absolute expression level. The Fidelity scores do not imply productive infection or cell-type abundance but rather the cellular context in which the receptors are expressed.

We first examined the overall distribution of Fidelity scores for all virus receptor genes across cortical areas to determine any biases in which cell-types express these receptors. Estimation statistics indicated that, overall, virus receptor genes showed greater enrichment for microglial expression signatures and lower enrichment for neuronal signatures (Fig. 5A). Next, we examined the cell-type associations across DE-SWAN clusters. We plotted the cell-types for clusters in which the median unpaired differences for the Fidelity scores were over- or under-represented relative to the overall distribution of Fidelity scores per cluster (Fig. 5B). Astrocytes were only over-represented in Cluster C, and oligodendrocytes in Cluster D. In contrast, microglia were over-represented in 3 Clusters (E, H and J) including Cluster E that is enriched for Coronaviridae receptors. Neurons were over-represented in Cluster B, which is also enriched for Rhabdoviridae receptors, but neurons were under-represented across 5 of the other Clusters.

**Figure 5.**
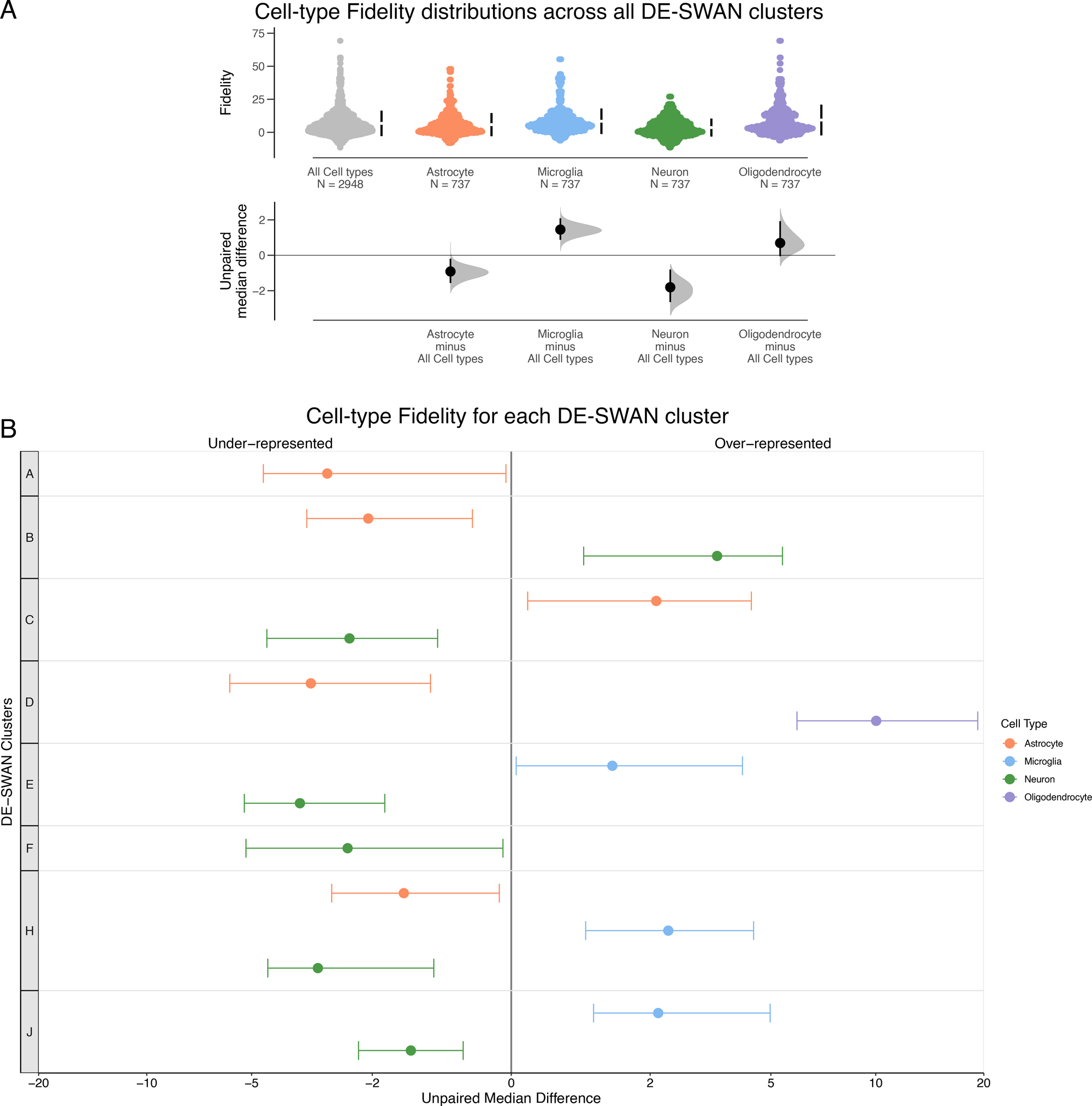
Fidelity-based cell-type associations of virus receptor genes across DE-SWAN clusters. (A) Swarm plots showing the distribution of Fidelity scores for all virus receptor genes across four major cortical cell types (astrocytes, microglia, neurons, oligodendrocytes). The left-most distribution represents the combined Fidelity scores across all cell types and serves as a reference. Estimation plots show unpaired median differences for each cell type relative to this reference, with dots indicating median differences, horizontal bars indicating 95% confidence intervals, and grey shading representing bootstrapped sampling distributions. Overall, virus receptor genes show higher Fidelity for microglia and lower Fidelity for neurons. (B) Lollipop plot summarizing cluster-specific cell-type associations across the 10 DE-SWAN clusters. Dots represent the unpaired median difference in Fidelity score for each cell type relative to the cluster-specific reference distribution (all trajectories within that cluster), with horizontal bars indicating 95% confidence intervals. Only clusters with confidence intervals excluding zero are shown; positive values indicate over-representation and negative values indicate under-representation. Neurons are over-represented in Cluster B, astrocytes in Cluster C, oligodendrocytes in Cluster D, and microglia in Clusters E, H, and J.

We specifically examined the Zika virus candidate entry receptor AXL, which was found in two of the DE-SWAN clusters: Cluster C (ITC, M1C, MFC, STC, V1C, VFC), with astrocyte over-representation, and Cluster H (A1C, DFC, IPC, OFC, S1C), with microglia over-representation. In addition, these clusters exhibited distinct sex-by-age expression profiles (Fig. 4C), suggesting that different glial cell types may contribute to Zika-associated vulnerability at different developmental stages.

These analyses show that although virus families and putative cell-type associations are broadly distributed across developmental programs, a subset of sex-by-age expression patterns shows selective enrichment for particular virus families and more restricted cell-type profiles. These results define a structured landscape of sex-specific developmental variation in virus receptor expression in the human cortex.

### Prenatal expression of Zika virus candidate entry receptors

We next asked whether sex differences in Zika virus candidate entry receptors are already present before birth. To address this question, we examined prenatal cortical expression of the TAM receptors TYRO3, AXL, and MERTK together with the lectins CD209 and CLEC4M. Prenatal analyses were conducted separately from the postnatal trajectory analyses because prenatal cortical development is characterized by neurogenesis, progenitor proliferation, neuronal migration, cortical lamination, and early gliogenesis. Using prenatal cortical samples from the Kang et al. dataset (304 tissue samples; 162 female, 142 male; 13–37 post-conception weeks; 17 donors), we compared expression between the sexes using estimation statistics. Among the five genes examined, AXL and CD209 showed higher expression in males, TYRO3 showed higher expression in females, whereas MERTK and CLEC4M showed similar expression between the sexes (Fig. 6A–E). These findings indicate that sex differences in Zika virus candidate entry receptors are already established during prenatal cortical development and occur in both TAM receptor and lectin-mediated entry pathways.

**Figure 6.**
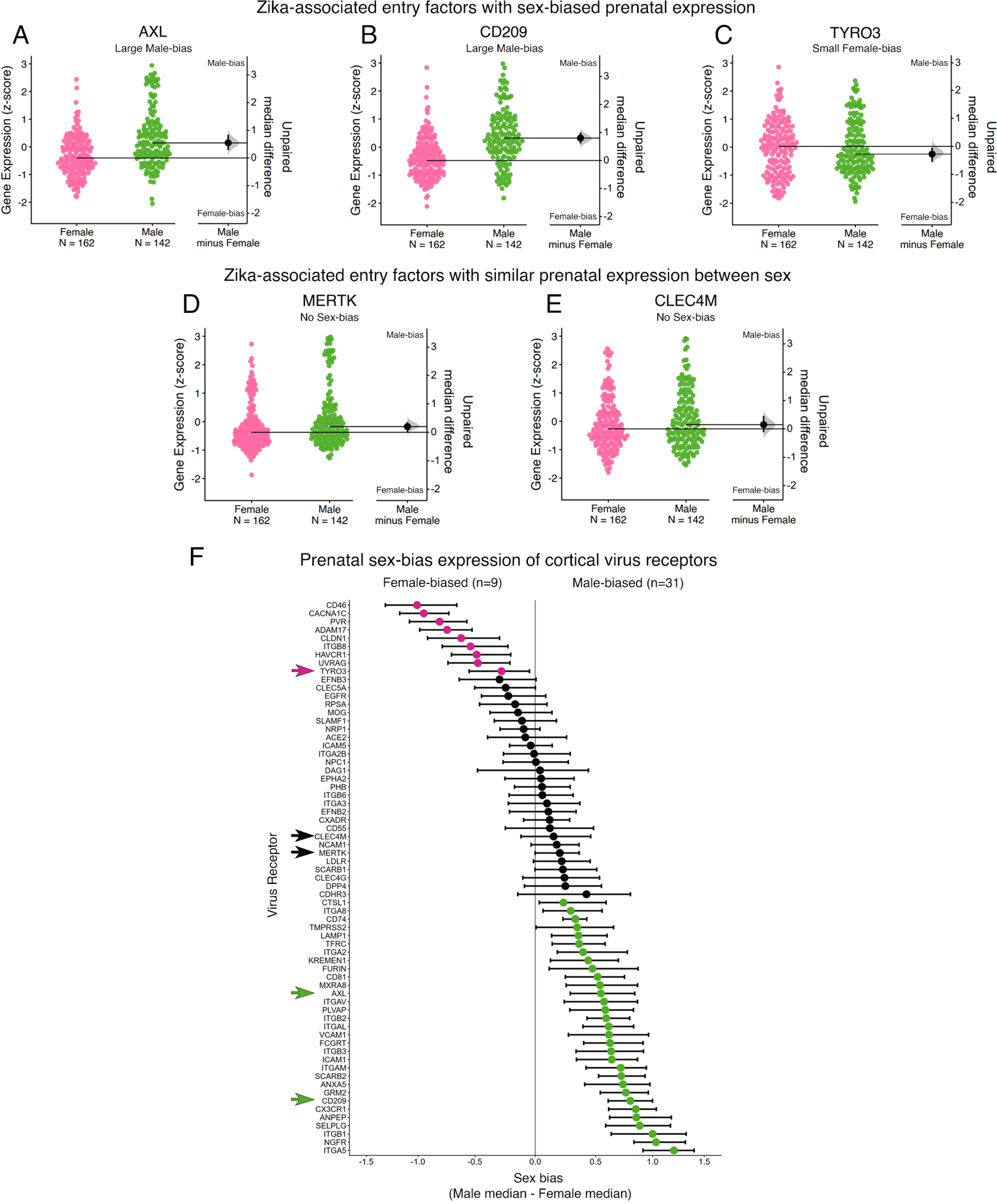
Prenatal expression of virus receptors in the cortex of females and males. (A-E) Swarm plots showing the expression of Zika virus candidate entry receptors with sex-biased prenatal expression (AXL, CD209, TYRO3; A–C) and similar prenatal expression between sexes (MERTK, and CLEC4M; D–E). Black dots represent the unpaired median difference between sexes, horizontal bars indicate 95% confidence intervals, and grey shading shows the bootstrapped sampling distribution. (F) Estimation statistics analysis of sex bias for all 67 genes. Virus receptor genes identified as female-biased are coloured magenta (n=9), male-biased are coloured green (n=31), and no-sex-bias are coloured black (n=27). The arrows identify the Zika virus candidate entry receptors shown in A–E, with the arrow colour indicating the sex-bias category.

### Prenatal cortical virus receptor expression is predominantly male-biased

To determine whether this pattern was unique toZika virus candidate entry receptors or reflected a broader feature of prenatal cortical development, we next examined all 67 virus receptor genes. Because several prenatal hemispheres did not contain the complete set of cortical regions available in the postnatal dataset, expression values were collapsed across cortical areas for this analysis. We used estimation statistics to analyze for sex biases in the expression of all 67 genes in the cortex. Prenatal cortical virus receptor expression exhibited a marked predominance of male-biased genes, with 31 receptors showing higher expression in males compared with only 9 showing higher expression in females (Fig. 6F). This prenatal male bias contrasts with the predominantly female-biased expression observed during the first postnatal year in the DE-SWAN analysis (Fig. 4), indicating that the direction of sex bias undergoes a major developmental transition across the prenatal-to-postnatal period.

Together, these findings indicate that sex-biased virus receptor expression is already established during prenatal cortical development but undergoes a marked developmental transition after birth, shifting from a predominantly male-biased molecular landscape during corticogenesis to predominantly female-biased expression during early postnatal cortical maturation.

## Discussion

This study provides the first developmental atlas of viral entry receptor expression in the human brain and demonstrates that sex differences are a fundamental feature of this system. Virus receptors follow non-linear developmental trajectories, many of which diverge by sex. These sex-biased cortical programs are sufficiently distinct at the population level to predict the sex of individual cases, yet they change dynamically across the lifespan, with divergence occurring most often during early childhood. Although virus families and inferred cell-types are broadly distributed across these developmental programs, a subset of sex-by-age expression patterns shows selective enrichment for specific viral lineages and more restricted, often glial-associated, cellular profiles. Together, these findings suggest that vulnerability and resilience to neuroinfectious disease may arise from the interplay among sex, developmental stage, cellular context, and molecular programs, rather than from receptor abundance alone.

### Sex-Biased Developmental Programs vs. Receptor Abundance Models

In many peripheral tissues, the abundance of virus-host entry receptors is a key determinant of tissue tropism and pathogenesis, as increased receptor expression can enhance viral entry and infection [48,49]. While this model has strong experimental support in peripheral tissues and transgenic systems, there are challenges explaining neurological outcomes in the human brain, where viral burden can be low or regionally restricted [5], and disease severity is not consistently mirrored by receptor abundance or widespread infection. Moreover, viral infection can alter neuronal activity and neurotransmitter release, leading to downstream disruption of synaptic plasticity and circuit function [9,16,50,51].

Our findings help refine this framework by showing that virus receptor expression in the cortex follows sex-biased developmental trajectories that vary across the lifespan, with some alignment for virus families and cell-type programs. Importantly, our findings do not address altered infection or replication; instead, they suggest that vulnerability may be shaped by the timing and cellular context of receptor expression rather than by receptor density alone.

### Dynamic trajectories and developmental windows

A central implication of our findings is that virus receptor expression in the cortex is sex-biased and developmentally regulated, with lifespan changes that are often undulating rather than monotonic or static. The widespread presence of non-linear trajectories, together with the high accuracy of the Random Forest classifier, indicates that these developmental programs reflect system-level organization rather than gene-level noise. Importantly, the classifier was not trained directly on individual cases but on population-level developmental trajectories and was subsequently challenged to classify individual expression profiles. Its successful performance demonstrates that these trajectories capture reproducible biological organization that generalizes from population-level developmental programs to individual brains, rather than simply describing average group trends. Together, these findings indicate that population-level developmental trajectories capture fundamental organizational principles of virus receptor expression that are preserved at the level of individual brains.

Many virus entry receptors participate in adhesion, chemokine signalling, or immune–synaptic crosstalk, placing them within transcriptional programs that regulate immune–neural interactions during circuit assembly. The dynamic trajectories imply age-dependent windows during which cortical circuits may be differentially susceptible to viral interaction. The coincidence of peak sex differences with sensitive periods of cortical development in childhood is notable and suggests that sex-biased organization of viral receptors may be particularly relevant during early circuit refinement. These receptors are not inert entry points but are embedded within neuroimmune and synaptic signalling networks that are themselves developmentally regulated and sexually dimorphic [52–58]. Many are expressed by microglia, which are central regulators of postnatal experience-dependent circuit maturation, positioning receptor trajectories at the interface of immune signalling and cortical circuit development. Such trajectories likely reflect coordinated regulation across interacting systems that sculpt developing cortical circuits. The timing of receptor expression relative to neurodevelopmental processes may therefore influence whether viral exposure produces transient perturbation, persistent circuit remodelling, or clinically significant pathology. Furthermore, recent transcriptomic analyses of human fetal brain explants further demonstrate that Zika infection suppresses synaptic signalling programs and disrupts the developmental transition from neurogenesis to astrogliogenesis [59], consistent with the idea that viral interactions occur within broader developmental molecular programs rather than through receptor abundance alone.

The targeted prenatal analyses further suggest that these developmental programs extend across the transition from corticogenesis to postnatal circuit maturation. During prenatal cortical development, virus receptor expression exhibited a marked predominance of male-biased genes, whereas the early postnatal period was characterized by predominantly female-biased expression. This developmental shift indicates that sex bias is not a static property of virus receptor expression but is itself developmentally regulated. Rather than maintaining a constant female or male advantage, the molecular landscape encountered by neurotropic viruses appears to be reorganized as the brain transitions from neurogenesis and cortical lamination before birth to synaptogenesis, experience-dependent plasticity, myelination, and neuroimmune maturation after birth. The predominance of male-biased prenatal receptor expression is consistent with emerging evidence from non-human primate models that congenital Zika infection produces sexually dimorphic outcomes. Male fetuses exhibit higher amniotic viral RNA concentrations and greater impairments in postnatal growth, whereas females show distinct alterations in early postnatal social behaviour [60].

One potential contributor to this developmental transition is the changing hormonal environment across the prenatal and early postnatal periods. Late gestation is characterized by increased androgen exposure in males, whereas the early postnatal period encompasses mini-puberty, during which estradiol and other gonadal hormones undergo dynamic changes [61]. Together with sex-chromosome effects and developmental changes in neuroimmune signalling, these endocrine transitions may contribute to the reorganization of viral receptor programs throughout early brain development.

This interpretation aligns with broader evidence that antiviral immune responses are sexually dimorphic and shift across developmental stages. Experimental and clinical studies demonstrate that sex hormones, chromosomal complement, and innate immune signalling pathways shape antiviral immunity differently in females and males [62,63]. Importantly, these immune differences are not static but evolve across childhood, puberty, and reproductive life. Epidemiological data similarly show that sex differences in viral susceptibility and severity vary across the lifespan, with male predominance often observed in early childhood infections (e.g., viral meningitis; [31,32]) and shifts emerging during adolescence and adulthood. Our findings provide a cortical molecular framework that may help contextualize how these systemic immune differences intersect with brain-specific developmental programs.

### Developmental cell-type programs and neurotropic disease

In our study, virus-family–enriched developmental programs aligned with distinct cellular contexts, suggesting that viral entry receptors are organized within cell-type-specific molecular programs. Consistent with these observations, the Zika virus candidate entry receptor AXL in our dataset was distributed across two distinct sex-by-age developmental programs. One showed higher expression in females during early childhood and was associated with astrocytes, whereas the other showed higher expression in males during late childhood and was associated with microglia. This partitioning mirrors experimentally observed shifts in Zika cell-type engagement across development and suggests that developmental reorganization of receptor expression may create distinct age- and sex-specific windows of cortical vulnerability.

Previous studies of Zika virus provide a particularly clear example of how developmental timing and cell-type specificity shape cortical vulnerability. Congenital infection disrupts neurogenesis and cortical development, resulting in microcephaly and other structural abnormalities [64]. Although early postnatal infection does not cause severe anatomical abnormalities, it does alter brain growth, neuronal and glial development and the development of emotional, social, motor, and cognitive function [11,64]. These developmental outcomes are sexually dimorphic in both mice and rhesus macaques, with prenatal and postnatal infection producing distinct sex-dependent developmental trajectories [60,65,66].

At the molecular level, transcriptomic and experimental studies demonstrate that Zika infection perturbs developmental gene-expression programs governing synaptic signalling, gliogenesis, and myelination [59,67]. In the developing human cortex, the candidate entry receptor AXL is highly expressed by astrocytes, microglia, radial glia, and endothelial cells [67], and experimental studies show that Zika exhibits age- and cell-type-specific tropism, initially targeting astrocytes and neural progenitors before infecting neurons [45]. In adult models, Zika can replicate in neurons, alter synaptic function, and induce glial dysfunction associated with aging-like phenotypes [12,66]. Collectively, these studies suggest that vulnerability to neurotropic viruses is determined not simply by the presence of an entry receptor, but by the developmental program in which that receptor is embedded.

Our finding that sex-biased trajectories are particularly prominent in the cortex further suggests that, when differences occur, viral interactions may intersect with mechanisms underlying experience-dependent development of cognitive and perceptual systems [11]. Although these concepts are well illustrated by Zika virus, they likely extend to other neurotropic viruses. For example, microglia are key mediators of synapse loss and cognitive impairment following West Nile virus infection [68], and also play an important role in sculpting neural circuits during the sensitive period of refinement in cortical development [72]. The higher incidence of neuroinvasive West Nile disease in males [69–71] further illustrates how age-, cell type-, and sex-dependent molecular programs may interact to influence neurological outcome following infection.

In addition to neurons and glia, many viral entry receptors are expressed by endothelial cells and pericytes that form the blood-brain barrier. Impairment of the blood-brain barrier by neurotropic viruses such as West Nile virus and Japanese encephalitis virus can lead to a rapid increase in permeability to viral particles [8,68,69]. Alternatively, disruption may induce indirect neurotoxic effects on astrocytes via infected pericytes, triggering downstream neuroimmune responses [70]. Future studies characterizing developmental changes in this system in females and males will be essential for understanding how vascular and parenchymal mechanisms converge to shape brain vulnerability.

### Limitations

Several limitations should be considered when interpreting these findings. First, the analyses are based on postmortem, cross-sectional transcriptomic data. Although the dataset spans a wide age range and includes rigorous quality control (PMI, pH, RIN, dissection score), the developmental inferences are derived from population-level patterns. As with all bulk transcriptomic datasets, the measured mRNA levels do not directly reflect protein abundance, receptor surface localization, or functional capacity [71,72]. Moreover, immune status, hormonal state, and socioeconomic variables were unavailable, limiting the ability to link receptor expression patterns to prior viral exposure or environmental context.

Second, although we inferred cellular context using Fidelity scores derived from single-cell datasets [47], these associations do not imply productive infection, viral replication, or cell-type abundance. Rather, they indicate the relative specificity of receptor gene expression across major cortical cell classes. In addition, finer cellular subclasses are not available in the Fidelity database, so subtle sex differences within subclasses could not be assessed.

Despite these limitations, the study has several important strengths. The dataset represents one of the most comprehensive publicly available human lifespan transcriptomic resources, spanning infancy to older adulthood across multiple cortical and subcortical regions [73]. Our analytic framework integrates sex as a biological variable at every stage of analysis and employs a series of complementary quantitative approaches. The convergence of these strategies strengthens confidence that the observed sex-biased developmental programs reflect structured biological organization rather than analytical artifact.

These strengths and limitations frame the current findings as a high-resolution atlas of sex-biased developmental organization of virus receptor expression in the human brain, while highlighting the need for future studies incorporating spatial transcriptomics, proteomic validation, and sex-balanced experimental models to directly test how these molecular programs influence neuroinfectious disease outcomes.

## Conclusions

As climate-driven expansion of neurotropic viruses increases global exposure [4,74,75], understanding the biological determinants of neurological severity becomes increasingly urgent. Our findings advance a framework in which developmentally regulated, sex-biased receptor programs define the molecular terrain upon which neurotropic viral exposure unfolds. Rather than viewing receptor abundance as a static determinant of infection, this work supports a model in which vulnerability emerges at the intersection of developmental stage, cellular maturation and synaptic refinement. Within this framework, sex-biased viral receptor programs are themselves developmentally reorganized across the lifespan, defining distinct molecular landscapes that may influence vulnerability to neurotropic viruses at different stages of life. By defining normative developmental trajectories of virus receptor expression, this work establishes a reference framework against which future studies of neurotropic viral infection can evaluate disease-associated molecular changes..

## Supporting Information

**Supplementary Figure 1** *Lifespan trajectories for 67 virus receptor genes across 15 brain areas.* Locally estimated scatterplot smoothing (LOESS) curves are shown for females (magenta) and males (green) for all 67 genes across all 15 brain areas studied. Z-scored expression values are plotted against age on a logarithmic scale. Shaded bands indicate the 95% confidence interval.

**Supplementary Figure 2** *Leave*-*one*-*case-out resampling to assess trajectory robustness and the stability of the sex differences.* For a random subset of 100 gene–area combinations, female and male LOESS trajectories were repeatedly refit across 10,000 leave-one-case-out iterations, each time omitting one female and one male case. In each iteration, the Fréchet distance between refit female and male trajectories was recalculated, generating a distribution of simulated distances for each combination. Estimation statistics were used to compare the original Fréchet distance with the mean simulated value, quantified as the unpaired median difference shown as a black dot with its 95% confidence interval; grey shading represents the bootstrap sampling distributions.

**Supplementary Figure 3** *Determining the number of clusters.* Elbow plot showing the within-cluster sum of squares plotted against the number of clusters (*k*). The optimal k-value (k = 25) is indicated as the inflection point in the curve, corresponding to the point of diminishing returns.

**Supplementary Figure 4** *Between-cluster estimation statistics of developmental trajectories.* Fréchet distances were calculated between each of the 25 developmental trajectory clusters to quantify similarity in trajectory shape. A–D) Swarm plots show the distributions of Fréchet distances and unpaired median differences for each cluster relative to a reference cluster (left-most distribution in each plot). A) Comparison of four clusters to reference cluster F83:M17; B) four clusters to reference F63:M37; C) all 25 clusters to reference F13:M87; D) all 25 clusters to reference F81:M19. Dots indicate median differences, horizontal bars represent 95% confidence intervals and the grey shading represents the bootstrapped sampling distributions. Larger median differences indicate greater dissimilarity in trajectory shape relative to the reference.

**Supplementary Figure 5** *Within-cluster estimation statistics of developmental trajectories*. A) Swarm plots show the Fréchet distances within each cluster, and estimation plots display the median differences relative to the distribution of all within-cluster distances. Black dots indicate the Hodges–Lehmann median difference, and black bars represent 95% confidence intervals. B) Distribution of within-cluster Fréchet distances, with the vertical lines indicating the median (blue) and the 97.5^th^ percentile (orange). The median of this within-cluster Fréchet distance distribution was taken as the reference point for defining similarity. Values below the median are considered developmentally similar, values between the median and 97.5^th^ percentile are within expected variation, and values greater than the 97.5^th^ percentile are developmentally dissimilar.

**Supplementary Figure 6** *Comparison of within*-*and between*-*cluster Fréchet distance distributions.* Distributions of Fréchet distances calculated within clusters and between clusters are shown, with the between-cluster distribution serving as the reference. The Hodges–Lehmann estimate of the median difference between distributions is indicated by a black dot, and horizontal lines denote the median of each distribution.

**Supplementary Figure 7** *den-SNE by sex and brain area.* Density-preserving t-SNE (den-SNE) projections of high-dimensional LOESS trajectory curves into two-dimensional space. Each curve represents a gene–brain area trajectory and is coloured by brain region and sex. Saturated pink and green denote female and male cortical areas, respectively, whereas pale pink and green denote female and male subcortical structures.

**Supplementary Figure 8** *Assessing robustness of DE-SWAN.* The number of significantly differentially expressed genes was quantified across age under different analysis parameters. A) Varying uncorrected p-value thresholds. B) Using q-values after Benjamini–Hochberg false discovery rate (FDR) correction. C) Varying sliding window sizes to assess robustness. Inflection points from these plots were used to define the window centers.

**Supplementary Figure 9** *Cell-type fidelity for DE-SWAN clusters*. Lollipop plot shows the unpaired median difference Fidelity score for clusters with under- or over-representation of a cell type. Dots represent median differences and horizontal lines indicate 95% confidence intervals.

**Supplementary Table 1.** *Donor information from Kang et al 2011.* Summary table of donors from the Kang et al 2011 dataset across 15 brain areas, including age, sex, and cause of death.

**Supplementary Table 2.** *Virus Receptor Gene List information.* List of viral receptor genes used in this study. Includes names, short forms, and gene symbols with associated viral species.

**Supplementary Table 3.** *Members of the 25 developmental trajectory clusters.* Table listing the developmental trajectories within each of the 25 clusters, as determined by hierarchical clustering of the 2,010 lifespan trajectories (67 virus receptor genes × 15 brain areas × 2 sexes).

**Supplementary Table 4.** *Members of the 10 DE-SWAN clusters.* Table listing the Gene-Area trajectories within each of the 10 DE-SWAN clusters (67 virus receptor genes × 11 brain areas).

## Author Contributions

LM, NH, KM Designed research; LM, NH Performed research; JL, PW Analyzed data; LM, NH, KM Wrote the first draft of the paper; MP, NB Edited the paper; LM, NH, MP, NB, KM Wrote the paper.

## Acknowledgements

We recognize that our McMaster University laboratory is located on the traditional territories of the Mississauga and Haudenosaunee nations and within the lands protected by the “Dish with One Spoon” wampum agreement. We thank Rachel Kwan, Brendan Kumagai, Ewalina Jeyanesan, Luke Bai and Adam Gee for help with coding and Dr. Robyn Klein for feedback on the manuscript.

## Data and Code accessibility

The processed dataset analyzed in the current study, along with the R code and supplementary materials (figures and tables), are available on the Open Science Framework (Monteiro et al., 2025) https://osf.io/4fgka/.

The transcriptomic dataset is from Kang et. al., (2011) and is available on the Gene Omnibus: (GSE25219).

