## Supplementary Figures for "Sex- and age-specific development of ssRNA virus receptor expression in the human brain": Suppl Fig 1.pdf

**Supplementary Materials**

**Supplementary Figure 1** *Lifespan trajectories for 67 genes across 15 brain areas.* Locally estimated scatterplot smoothing (LOESS) curves are shown for females females (magenta) and males (green) for all 67 genes across all 15 brain areas studied. Z-scored expression values are plotted against age on a logarithmic scale. Shaded bands indicate the 95% confidence interval.

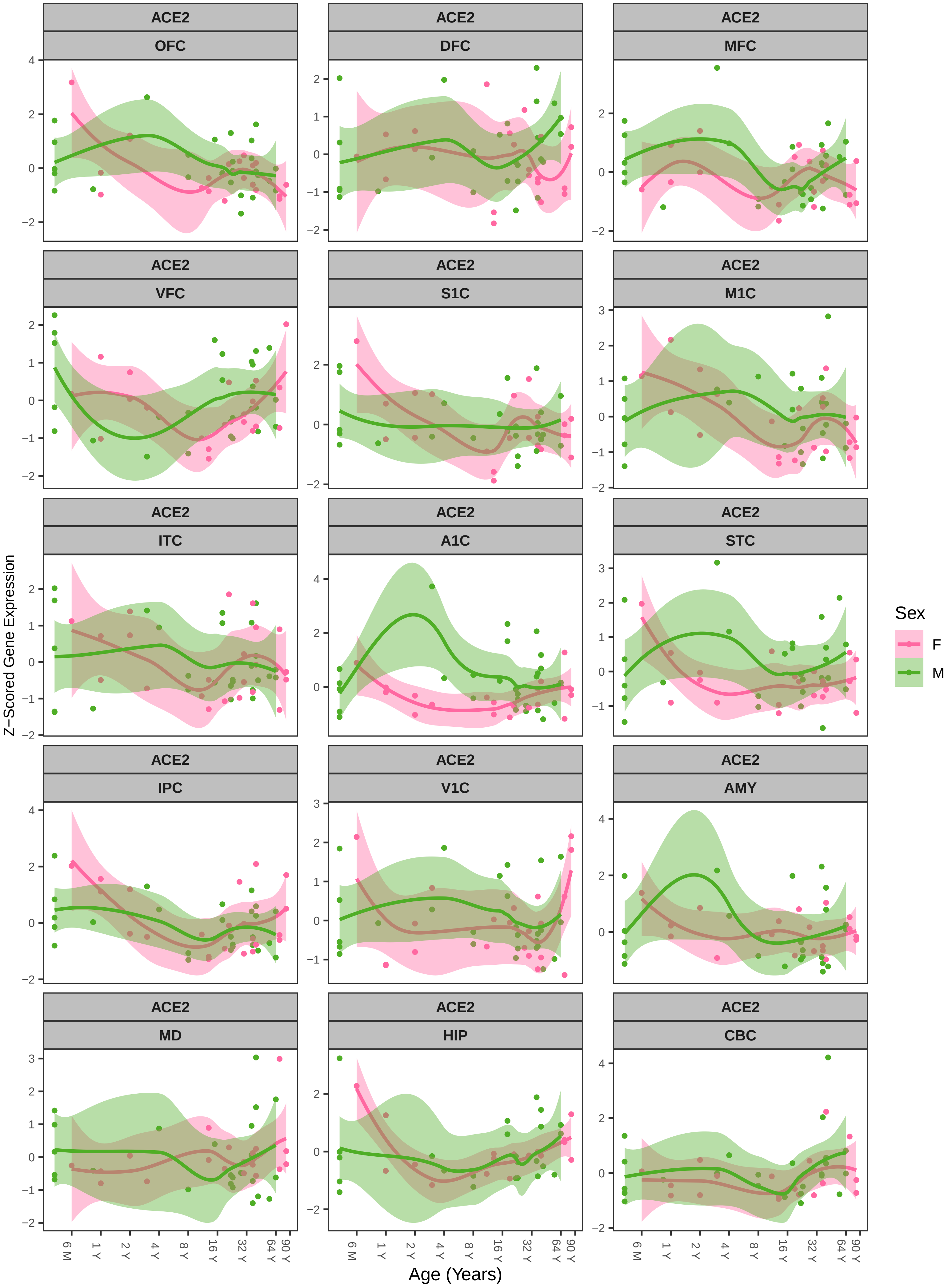

ACE2
OFC
ACE2
DFC
ACE2
MFC
ACE2
VFC
ACE2
S1C
ACE2
M1C
ACE2
ITC
ACE2
A1C
ACE2
STC
ACE2
IPC
ACE2
V1C
ACE2
AMY
ACE2
MD
ACE2
HIP
ACE2
CBC
Z−Scored Gene Expression
Age (Years)
6 M
1 Y
2 Y
4 Y
8 Y
16 Y
32 Y
64 Y
90 Y
Sex
F
M

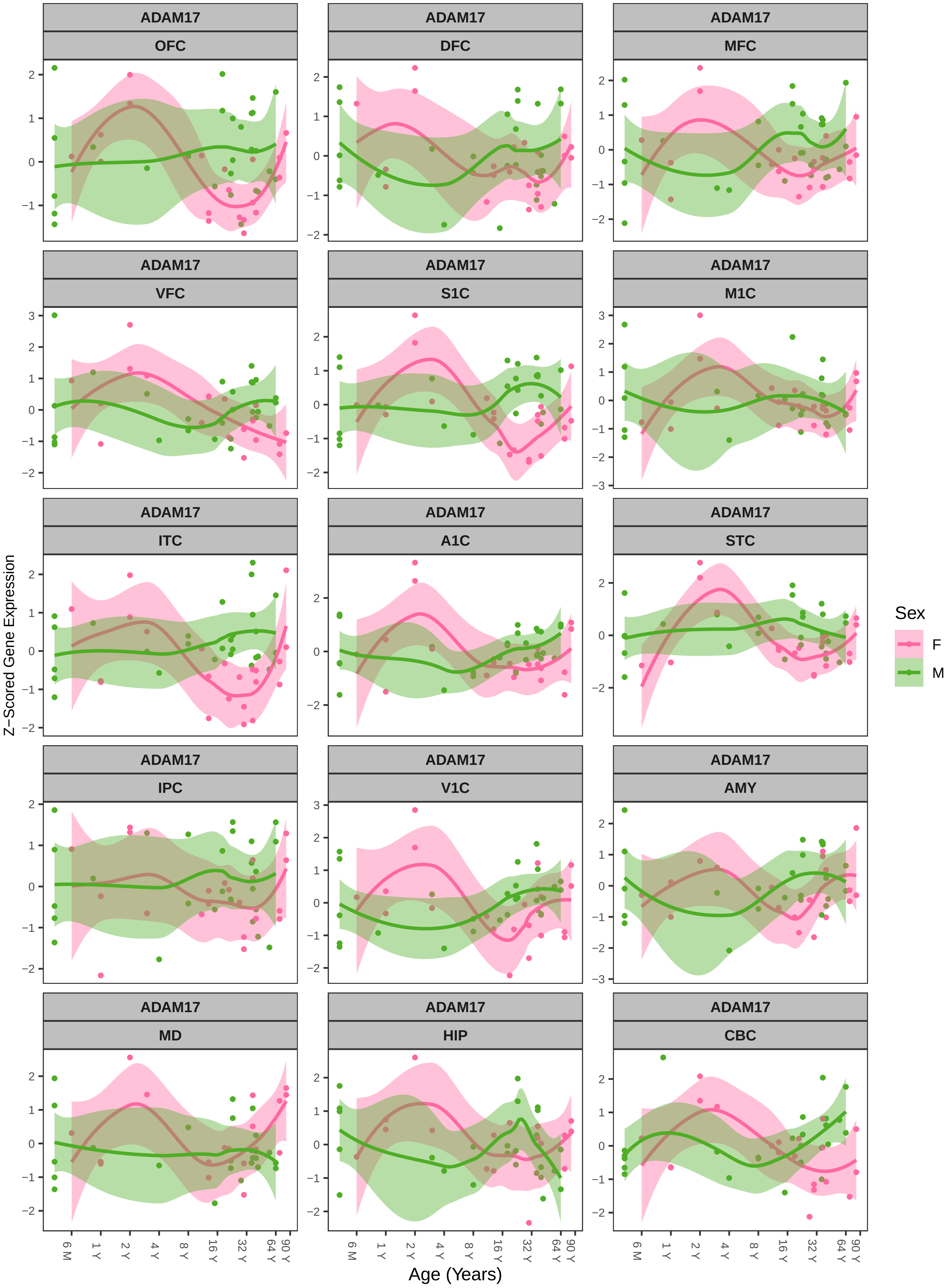
ADAM17
OFC
DFC
MFC
VFC
S1C
M1C
ITC
A1C
STC
IPC
V1C
AMY
MD
HIP
CBC
Z−Scored Gene Expression
Age (Years)
6 M
1 Y
2 Y
4 Y
8 Y
16 Y
32 Y
64 Y
90 Y
Sex
F
M

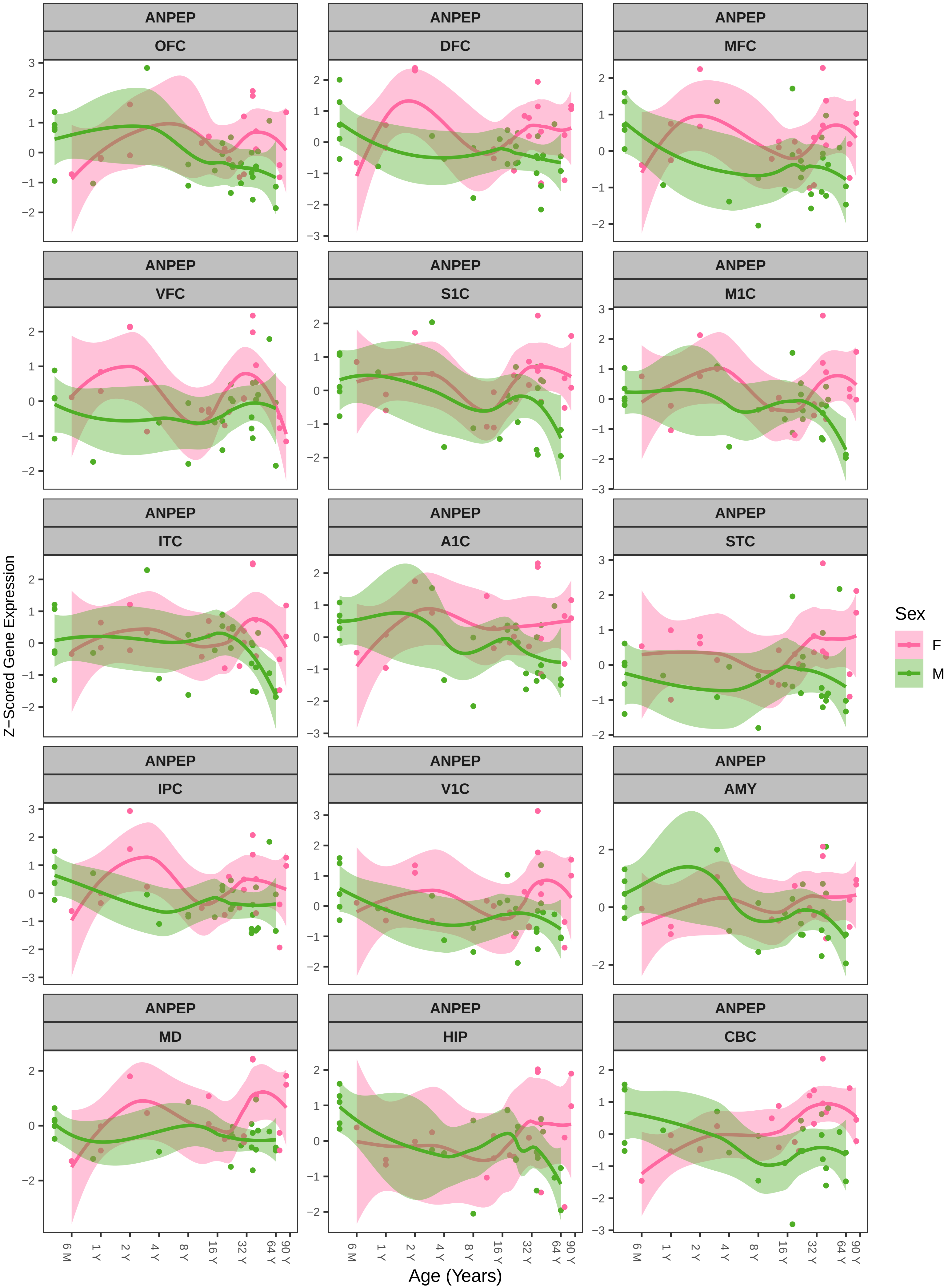
ANPEP
OFC
DFC
MFC
VFC
S1C
M1C
ITC
A1C
STC
IPC
V1C
AMY
MD
HIP
CBC
Z−Scored Gene Expression
Age (Years)
6 M
1 Y
2 Y
4 Y
8 Y
16 Y
32 Y
64 Y
90 Y
Sex
F
M

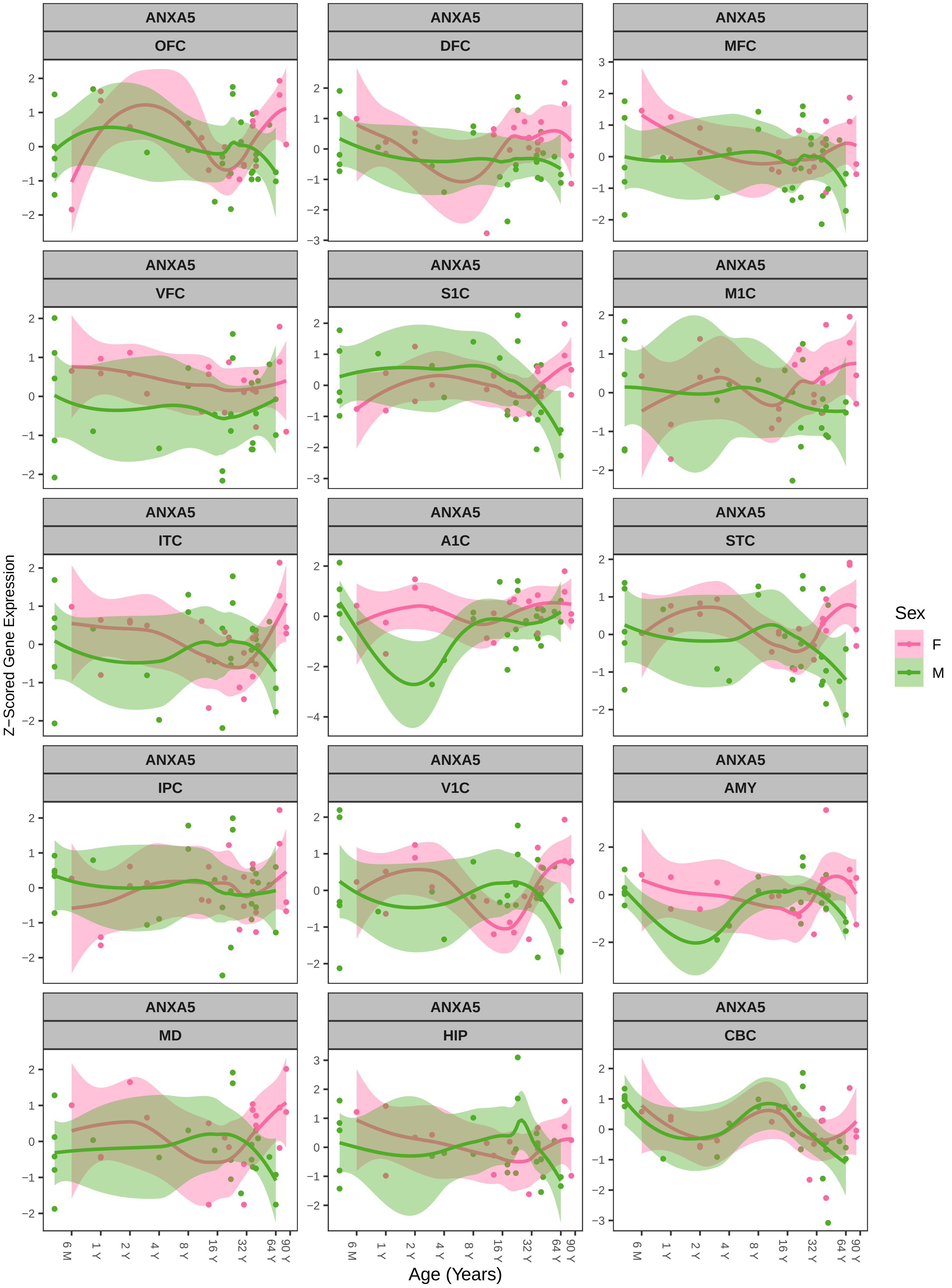
ANXA5
OFC
DFC
MFC
VFC
S1C
M1C
ITC
A1C
STC
IPC
V1C
AMY
MD
HIP
CBC
Z−Scored Gene Expression
Age (Years)
6 M
1 Y
2 Y
4 Y
8 Y
16 Y
32 Y
64 Y
90 Y
Sex
F
M

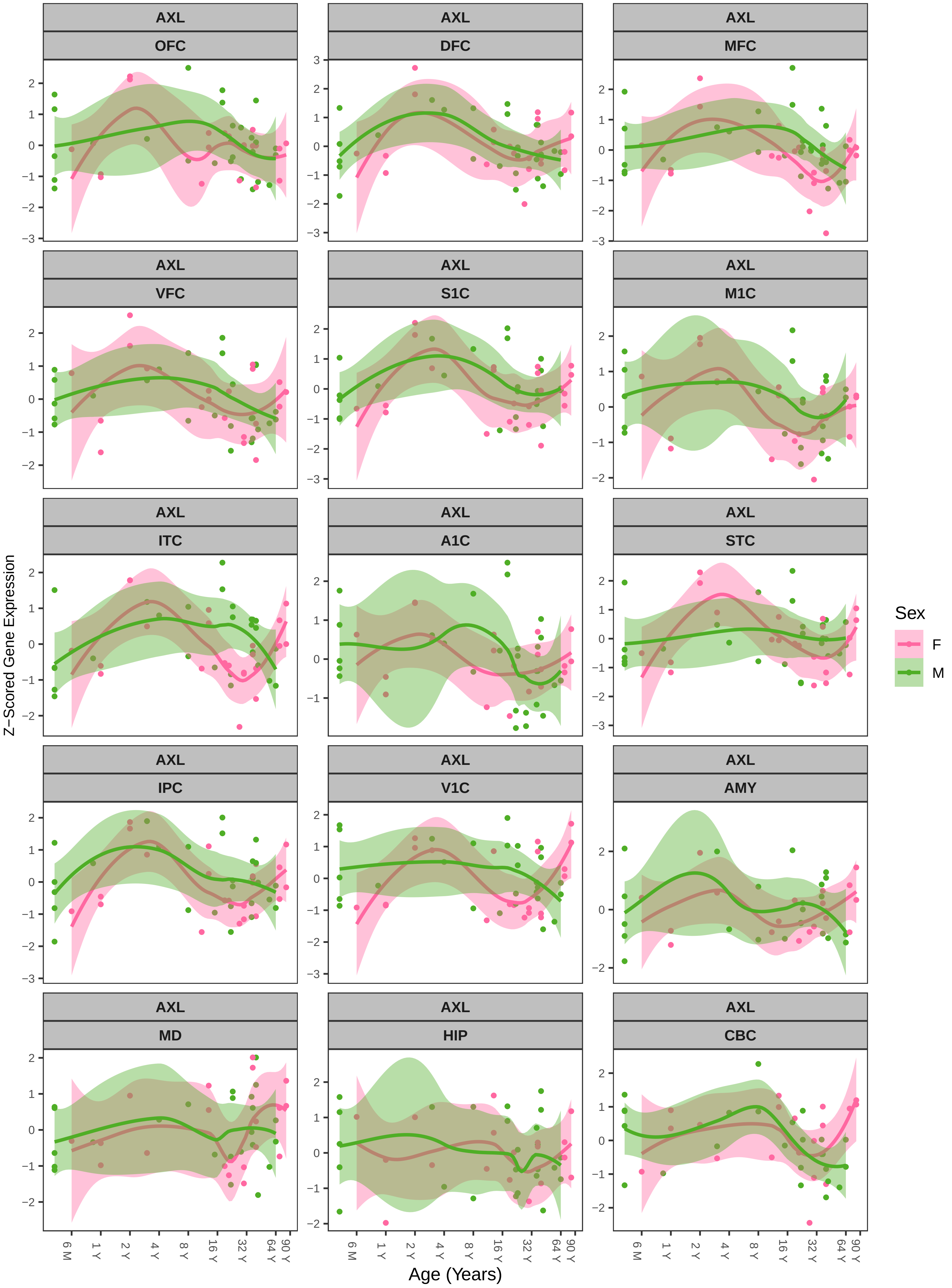

AXL
OFC
DFC
MFC
VFC
S1C
M1C
ITC
A1C
STC
IPC
V1C
AMY
MD
HIP
CBC
Z−Scored Gene Expression
Age (Years)
6 M
1 Y
2 Y
4 Y
8 Y
16 Y
32 Y
64 Y
90 Y
Sex
F
M

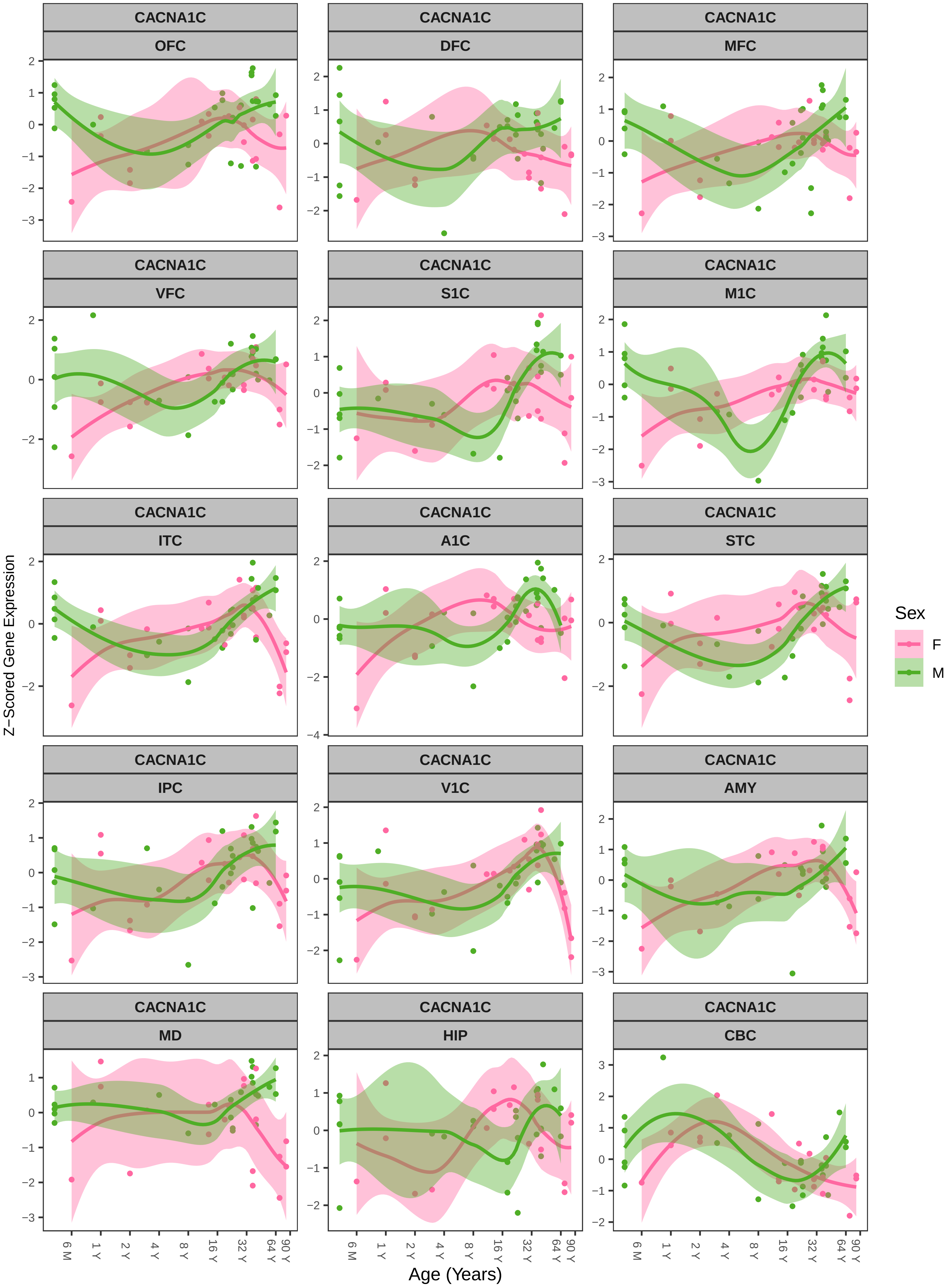

CACNA1C
OFC
DFC
MFC
VFC
S1C
M1C
ITC
A1C
STC
IPC
V1C
AMY
MD
HIP
CBC
Z−Scored Gene Expression
Age (Years)
6 M
1 Y
2 Y
4 Y
8 Y
16 Y
32 Y
64 Y
90 Y
Sex
F
M

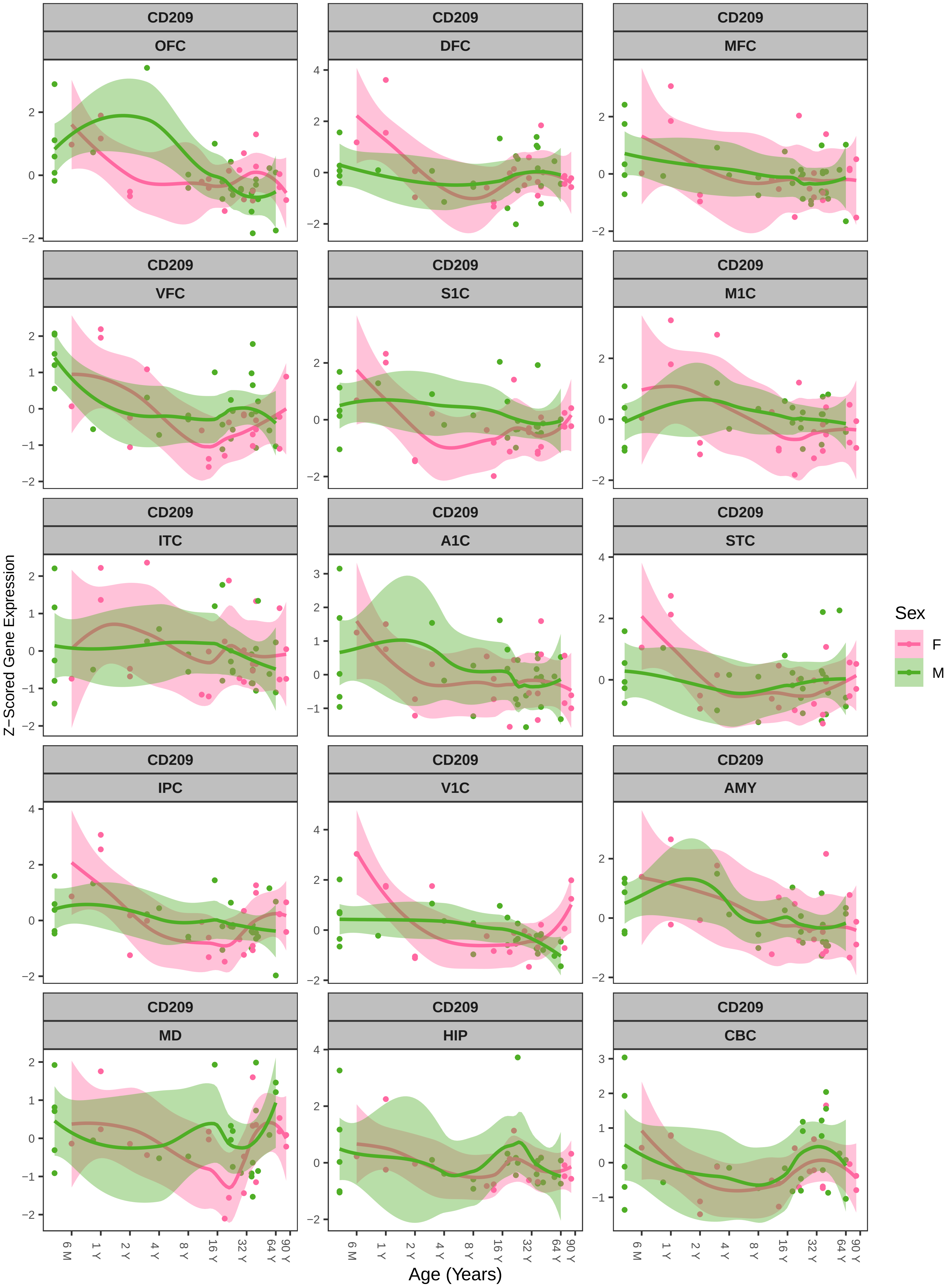

CD209
OFC
DFC
MFC
VFC
S1C
M1C
ITC
A1C
STC
IPC
V1C
AMY
MD
HIP
CBC
Z−Scored Gene Expression
Age (Years)
6 M
1 Y
2 Y
4 Y
8 Y
16 Y
32 Y
64 Y
90 Y
Sex
F
M

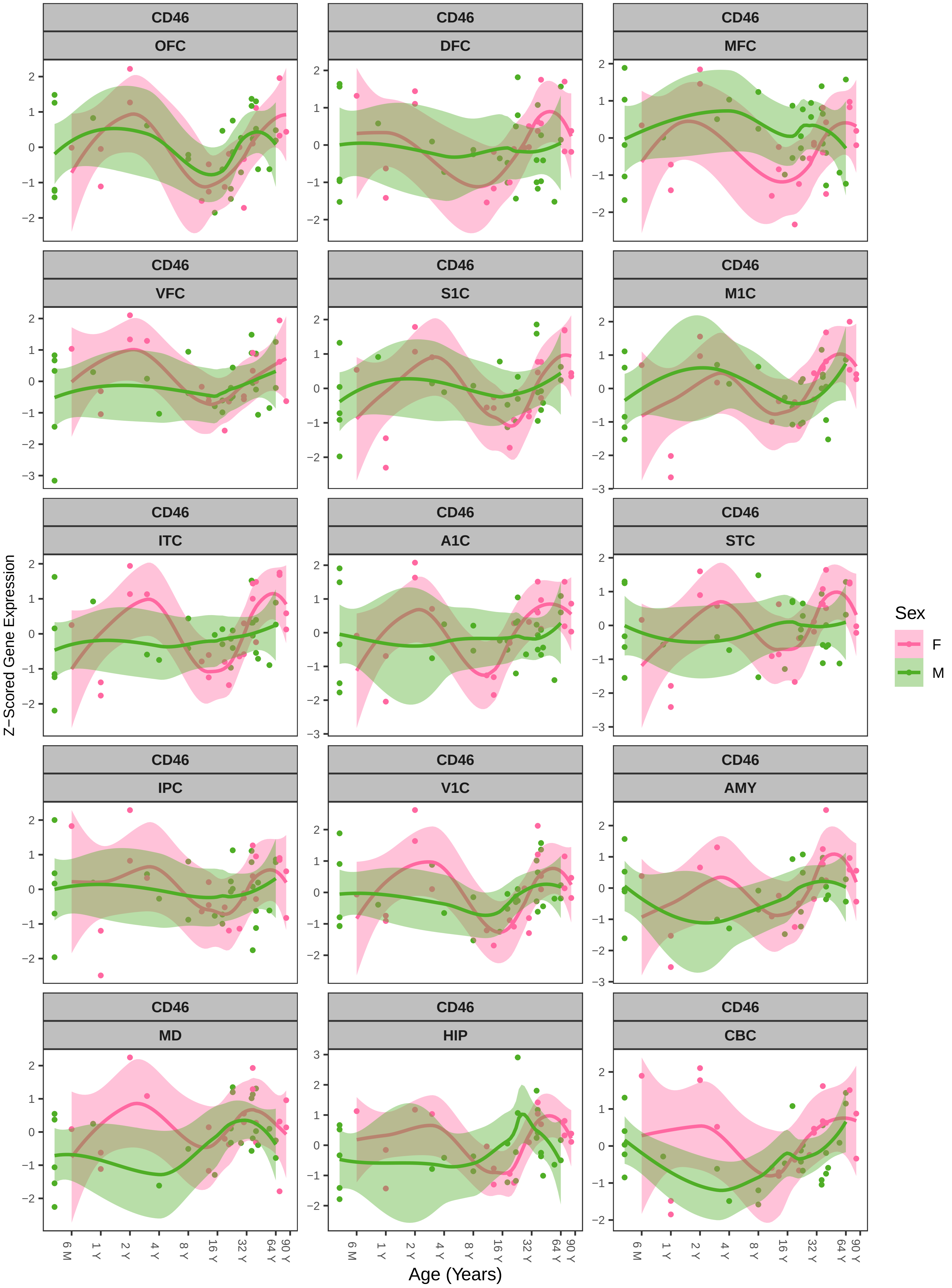
CD46
OFC
DFC
MFC
VFC
S1C
M1C
ITC
A1C
STC
IPC
V1C
AMY
MD
HIP
CBC
Z−Scored Gene Expression
Age (Years)
6 M
1 Y
2 Y
4 Y
8 Y
16 Y
32 Y
64 Y
90 Y
Sex
F
M

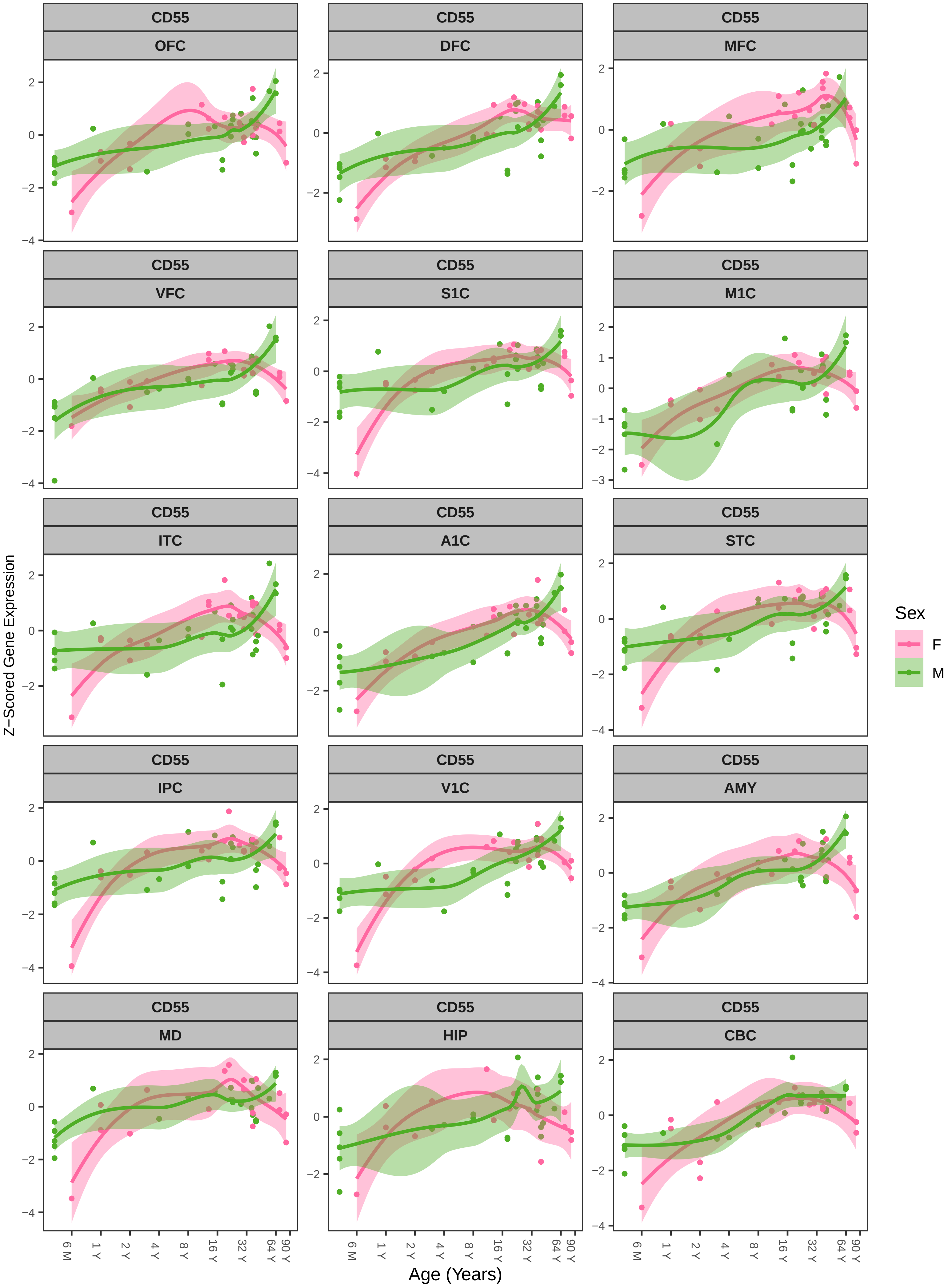
CD55
OFC
CD55
DFC
CD55
MFC
CD55
VFC
CD55
S1C
CD55
M1C
CD55
ITC
CD55
A1C
CD55
STC
CD55
IPC
CD55
V1C
CD55
AMY
CD55
MD
CD55
HIP
CD55
CBC
Z−Scored Gene Expression
Age (Years)
6 M
1 Y
2 Y
4 Y
8 Y
16 Y
32 Y
64 Y
90 Y
Sex
F
M

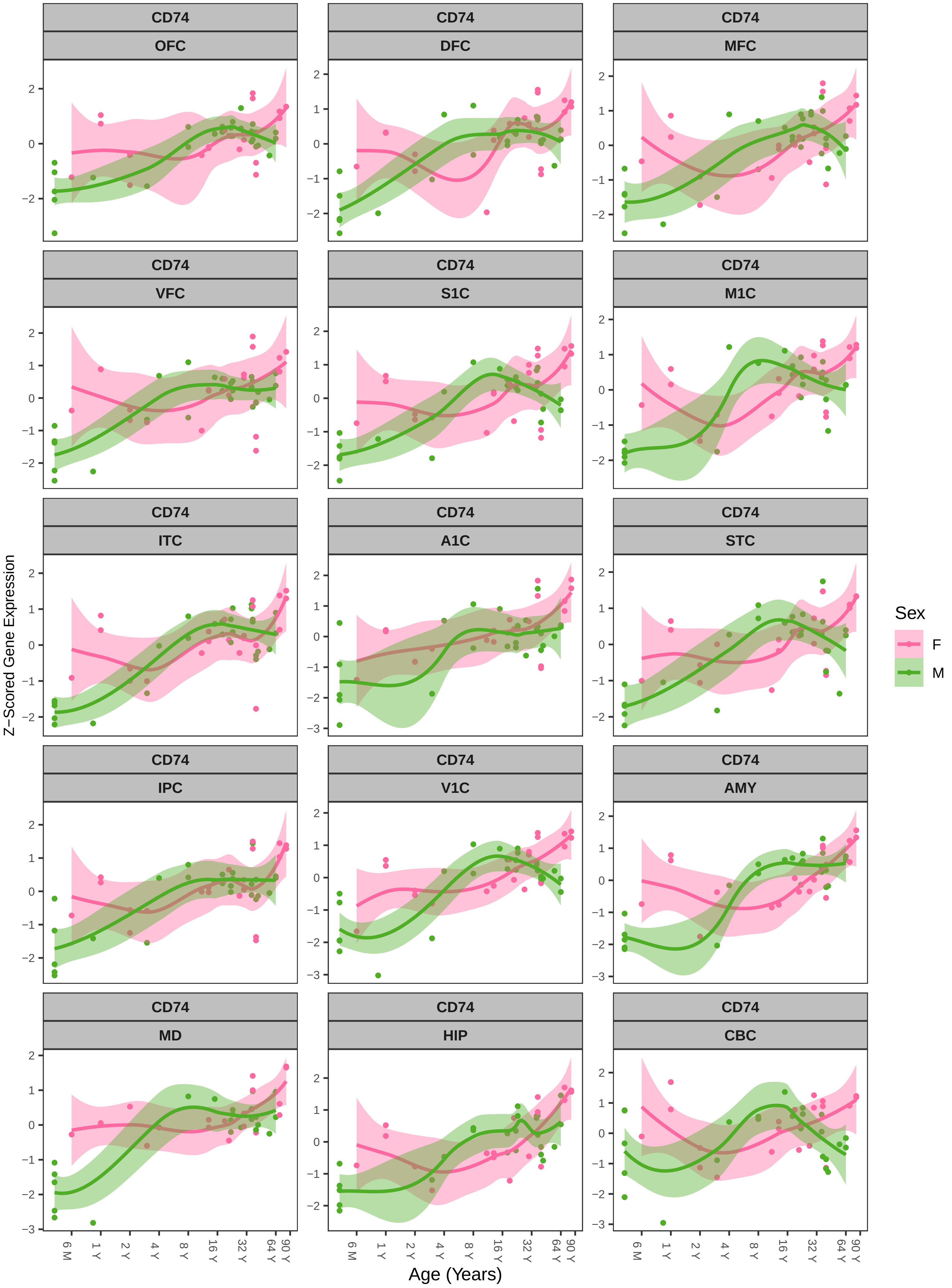

CD74
OFC
DFC
MFC
VFC
S1C
M1C
ITC
A1C
STC
IPC
V1C
AMY
MD
HIP
CBC
Z−Scored Gene Expression
Age (Years)
6 M
1 Y
2 Y
4 Y
8 Y
16 Y
32 Y
64 Y
90 Y
Sex
F
M

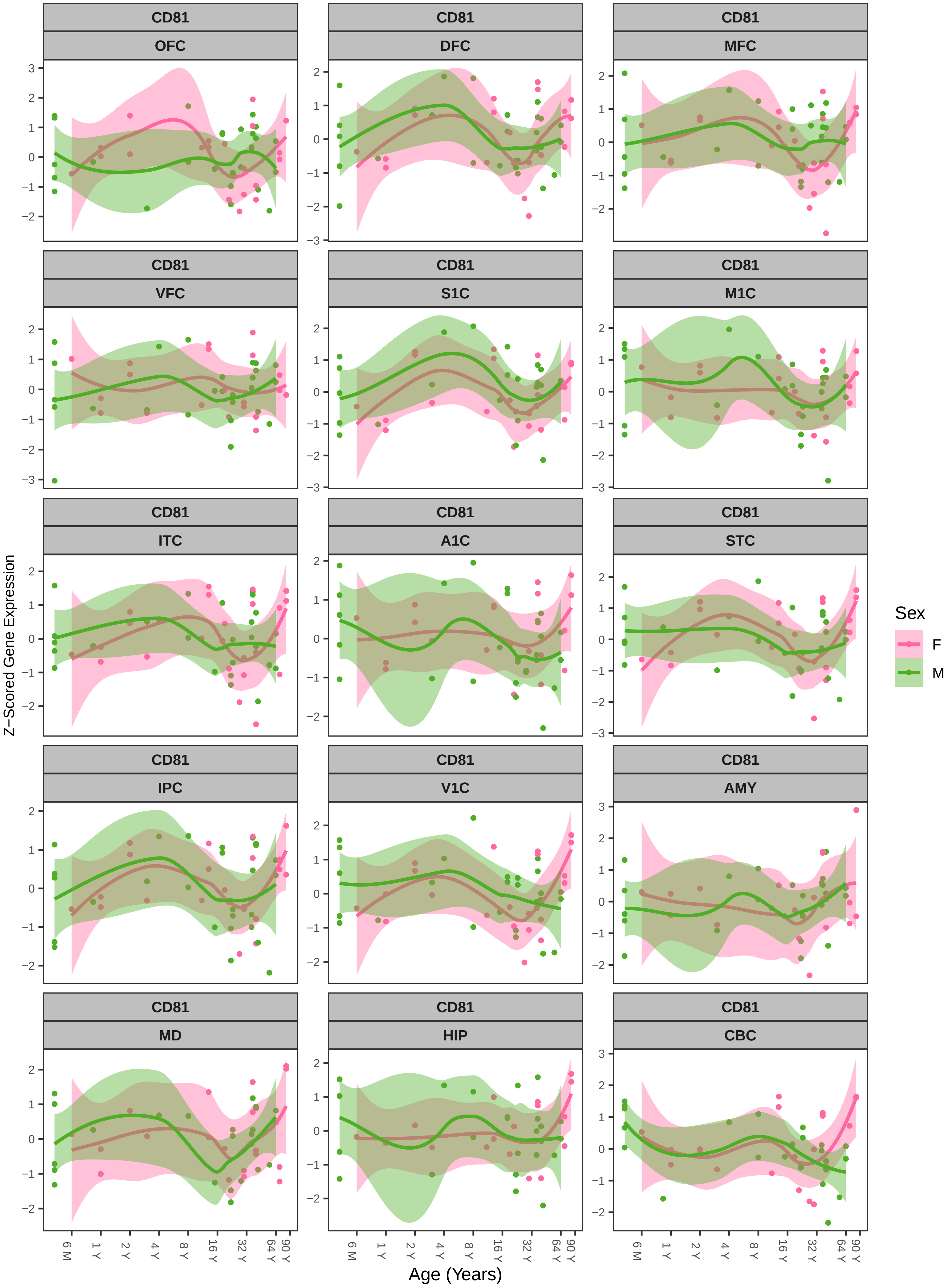
CD81
OFC
DFC
MFC
VFC
S1C
M1C
ITC
A1C
STC
IPC
V1C
AMY
MD
HIP
CBC
Z−Scored Gene Expression
Age (Years)
6 M
1 Y
2 Y
4 Y
8 Y
16 Y
32 Y
64 Y
90 Y
Sex
F
M

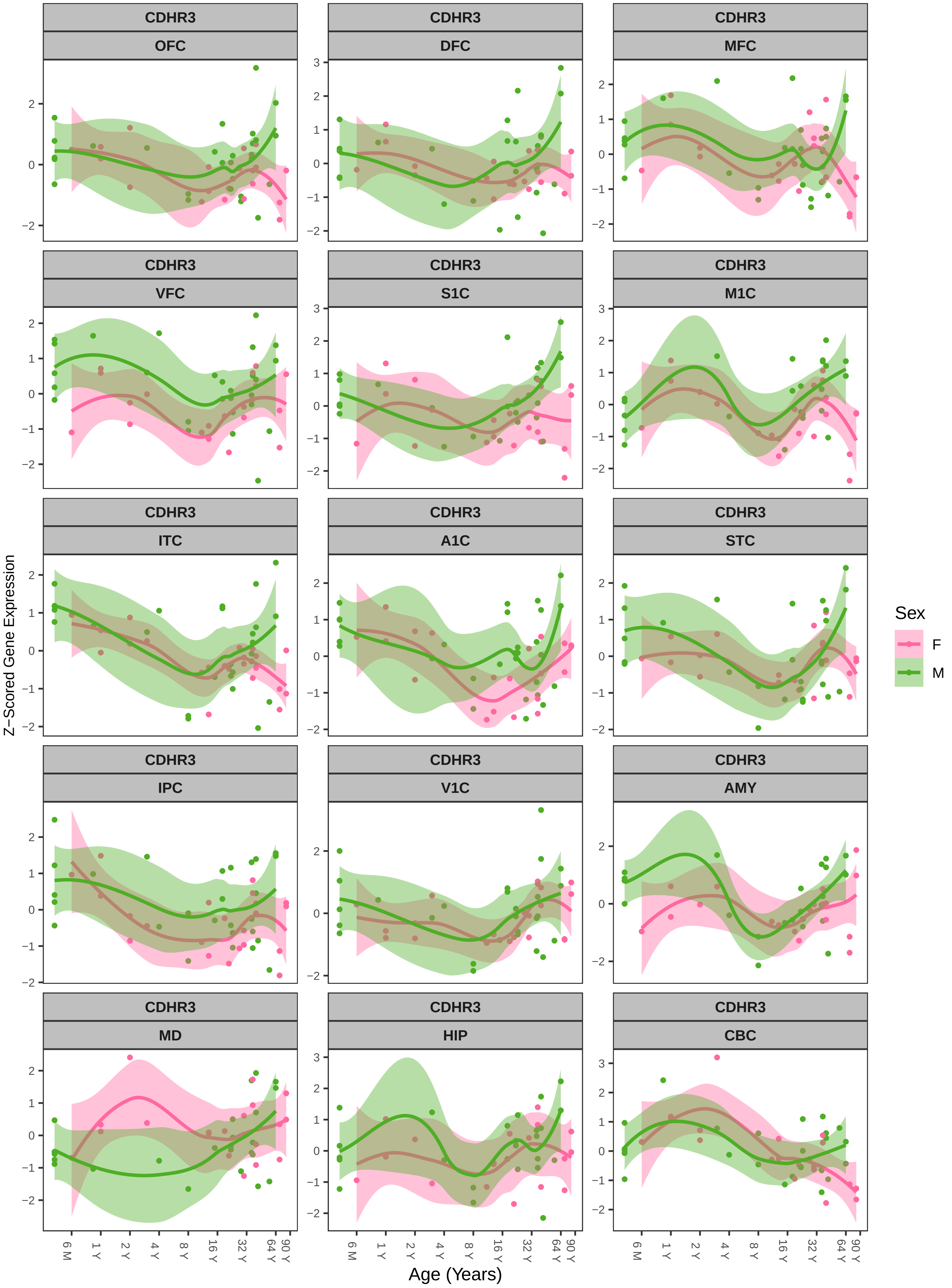
CDHR3
OFC
DFC
MFC
VFC
S1C
M1C
ITC
A1C
STC
IPC
V1C
AMY
MD
HIP
CBC
Z−Scored Gene Expression
Age (Years)
6 M
1 Y
2 Y
4 Y
8 Y
16 Y
32 Y
64 Y
90 Y
Sex
F
M

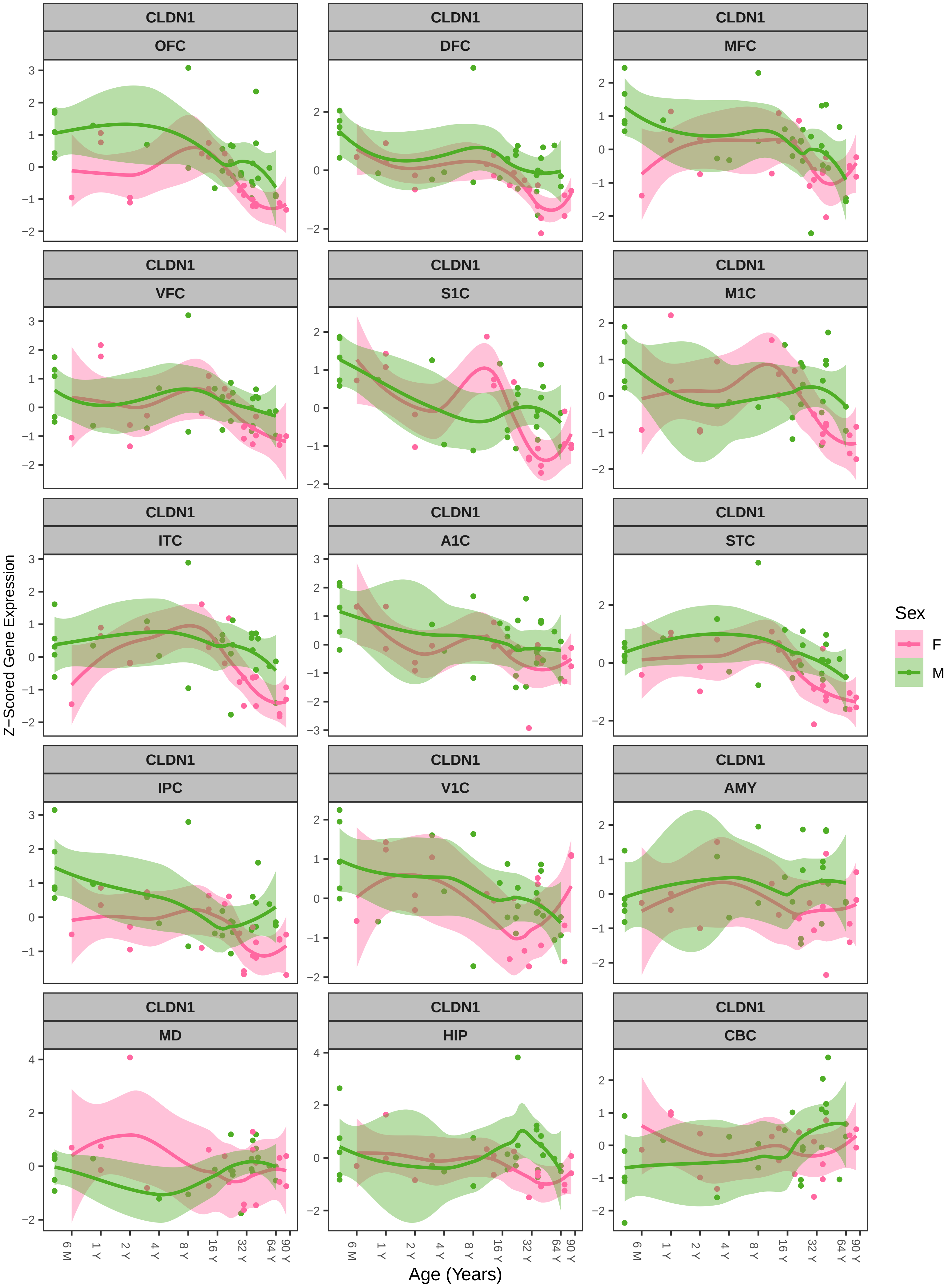
CLDN1
OFC
DFC
MFC
VFC
S1C
M1C
ITC
A1C
STC
IPC
V1C
AMY
MD
HIP
CBC
Z−Scored Gene Expression
Age (Years)
6 M
1 Y
2 Y
4 Y
8 Y
16 Y
32 Y
64 Y
90 Y
Sex
F
M

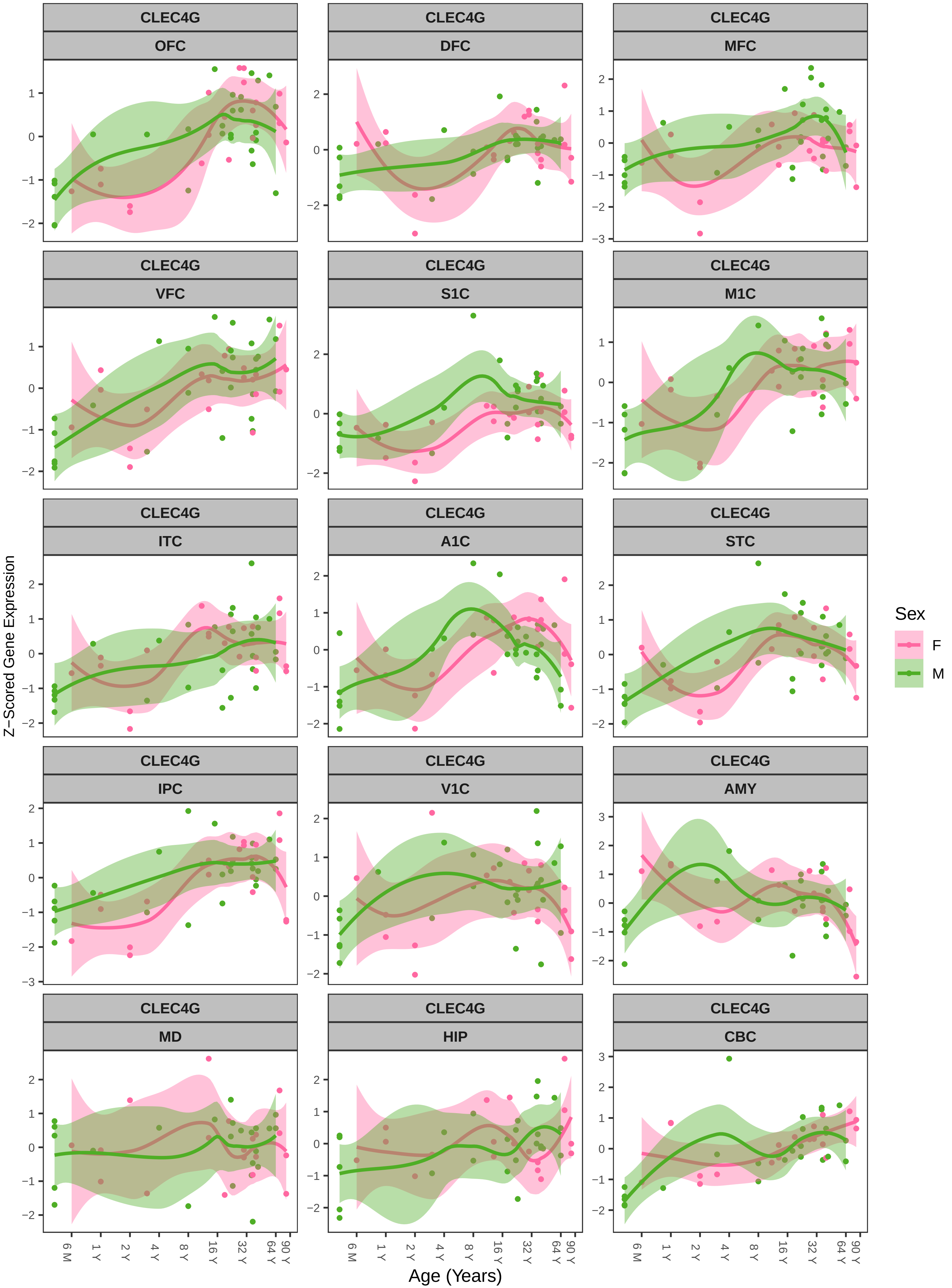

CLEC4G
OFC
DFC
MFC
VFC
S1C
M1C
ITC
A1C
STC
IPC
V1C
AMY
MD
HIP
CBC
Z−Scored Gene Expression
Age (Years)
6 M
1 Y
2 Y
4 Y
8 Y
16 Y
32 Y
64 Y
90 Y
Sex
F
M

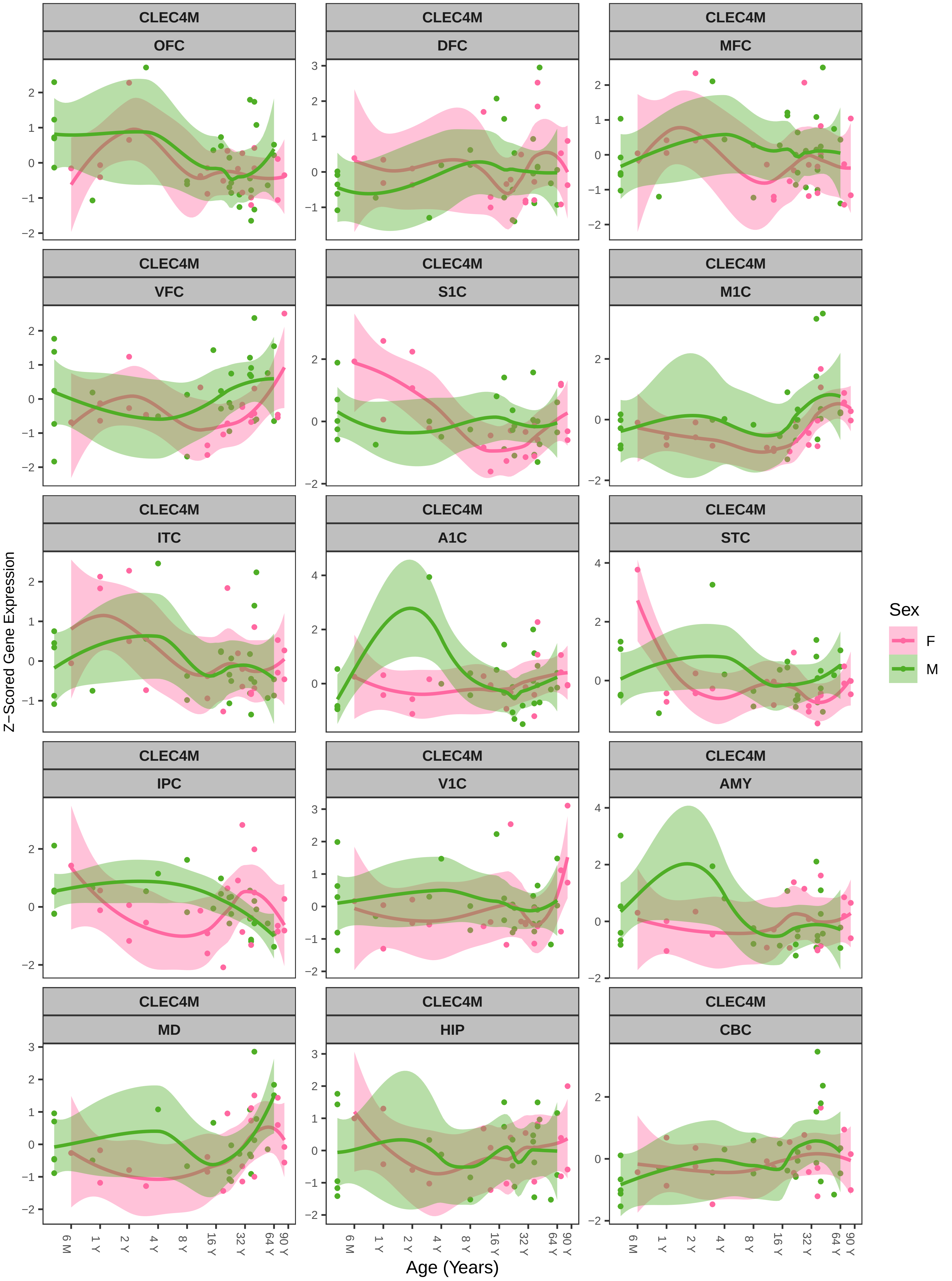

CLEC4M
OFC
DFC
MFC
VFC
S1C
M1C
ITC
A1C
STC
IPC
V1C
AMY
MD
HIP
CBC
Z−Scored Gene Expression
Age (Years)
6 M
1 Y
2 Y
4 Y
8 Y
16 Y
32 Y
64 Y
90 Y
Sex
F
M

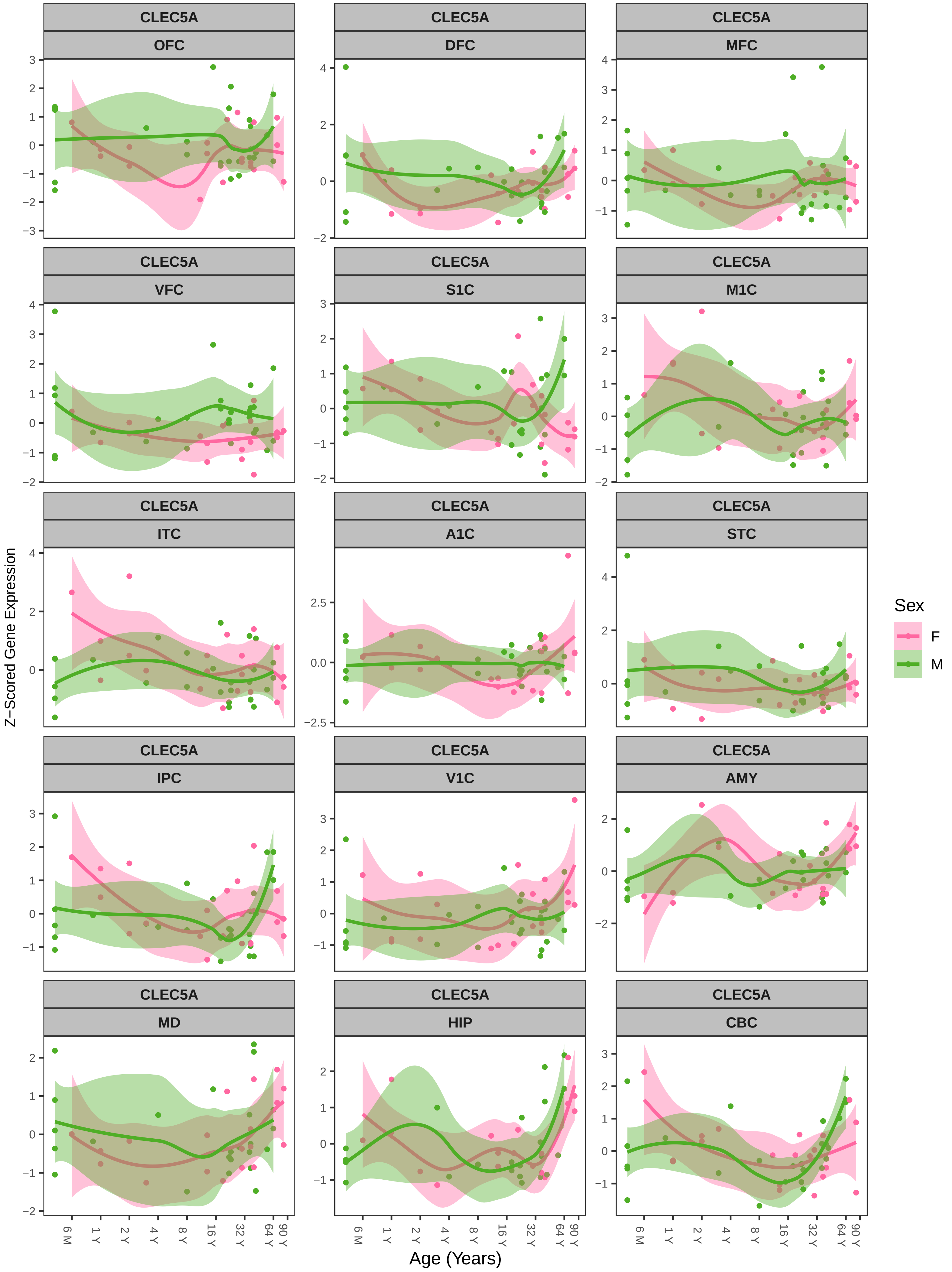
CLEC5A
OFC
DFC
MFC
VFC
S1C
M1C
ITC
A1C
STC
IPC
V1C
AMY
MD
HIP
CBC
Z−Scored Gene Expression
Age (Years)
6 M
1 Y
2 Y
4 Y
8 Y
16 Y
32 Y
64 Y
90 Y
Sex
F
M

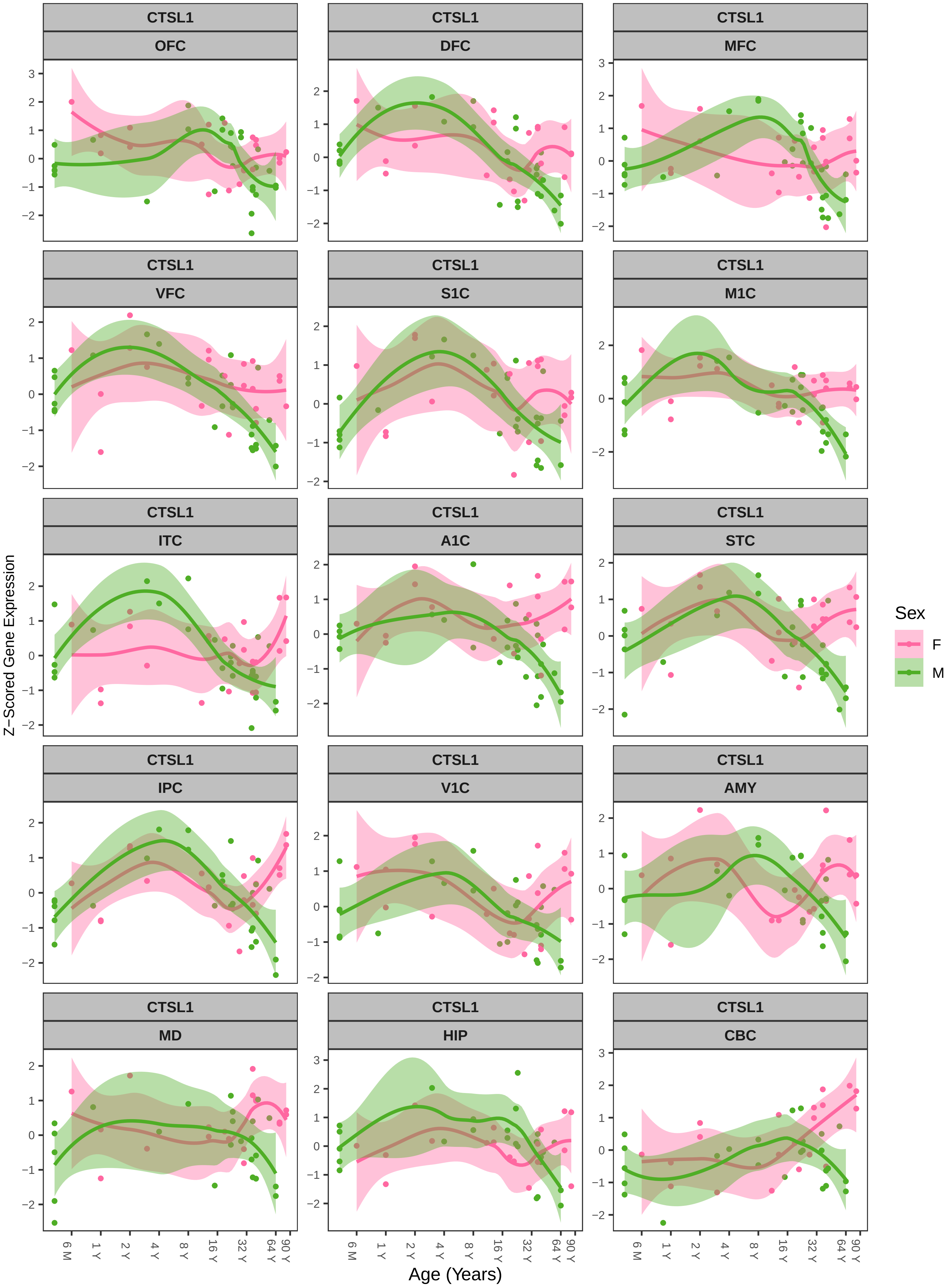
CTSL1
OFC
DFC
MFC
VFC
S1C
M1C
ITC
A1C
STC
IPC
V1C
AMY
MD
HIP
CBC
Z−Scored Gene Expression
Age (Years)
6 M
1 Y
2 Y
4 Y
8 Y
16 Y
32 Y
64 Y
90 Y
Sex
F
M

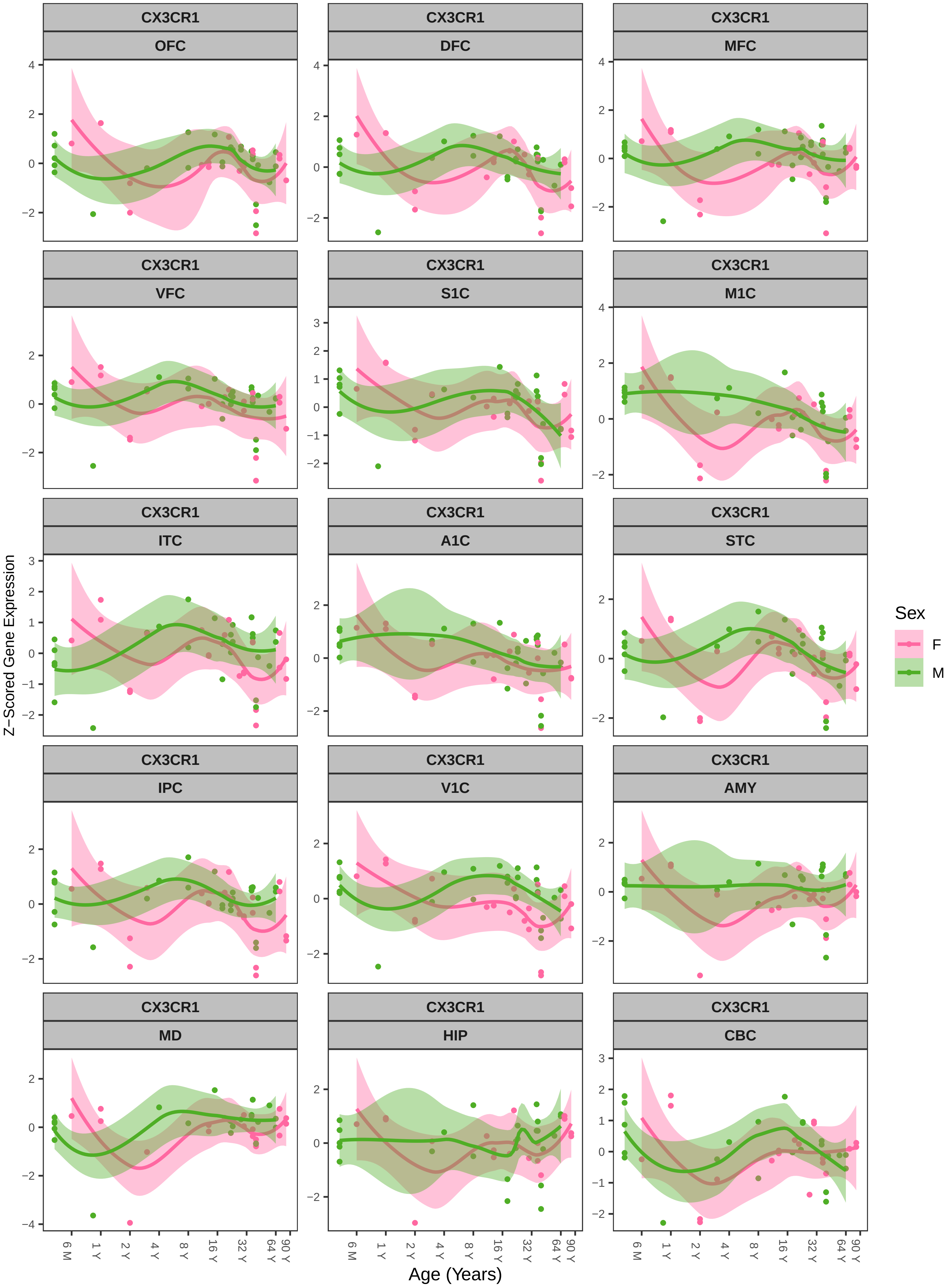
CX3CR1
OFC
DFC
MFC
VFC
S1C
M1C
ITC
A1C
STC
IPC
V1C
AMY
MD
HIP
CBC
Z−Scored Gene Expression
Age (Years)
6 M
1 Y
2 Y
4 Y
8 Y
16 Y
32 Y
64 Y
90 Y
Sex
F
M

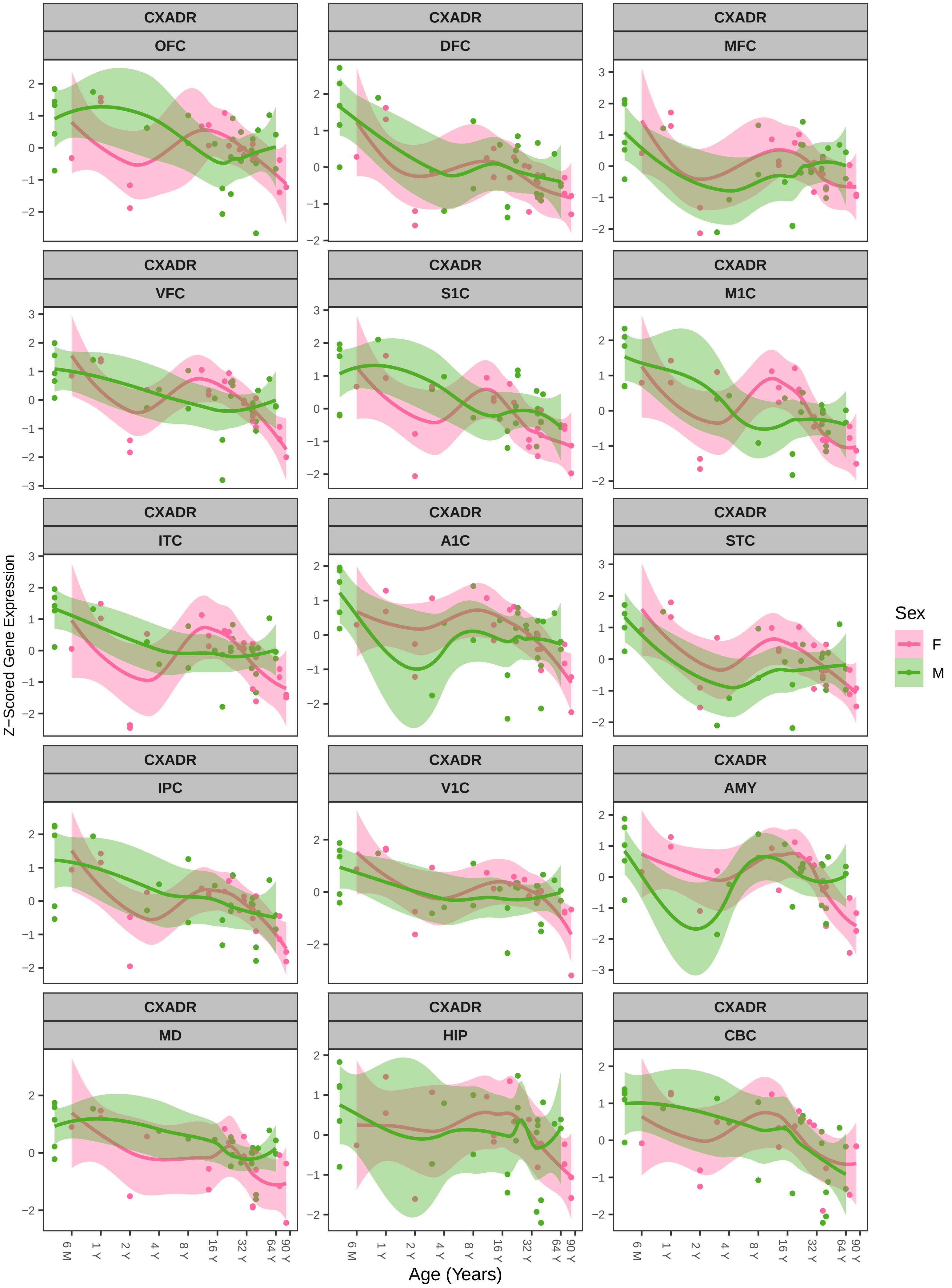
CXADR
OFC
CXADR
DFC
CXADR
MFC
CXADR
VFC
CXADR
S1C
CXADR
M1C
CXADR
ITC
CXADR
A1C
CXADR
STC
CXADR
IPC
CXADR
V1C
CXADR
AMY
CXADR
MD
CXADR
HIP
CXADR
CBC
Z−Scored Gene Expression
Age (Years)
6 M
1 Y
2 Y
4 Y
8 Y
16 Y
32 Y
64 Y
90 Y
Sex
F
M

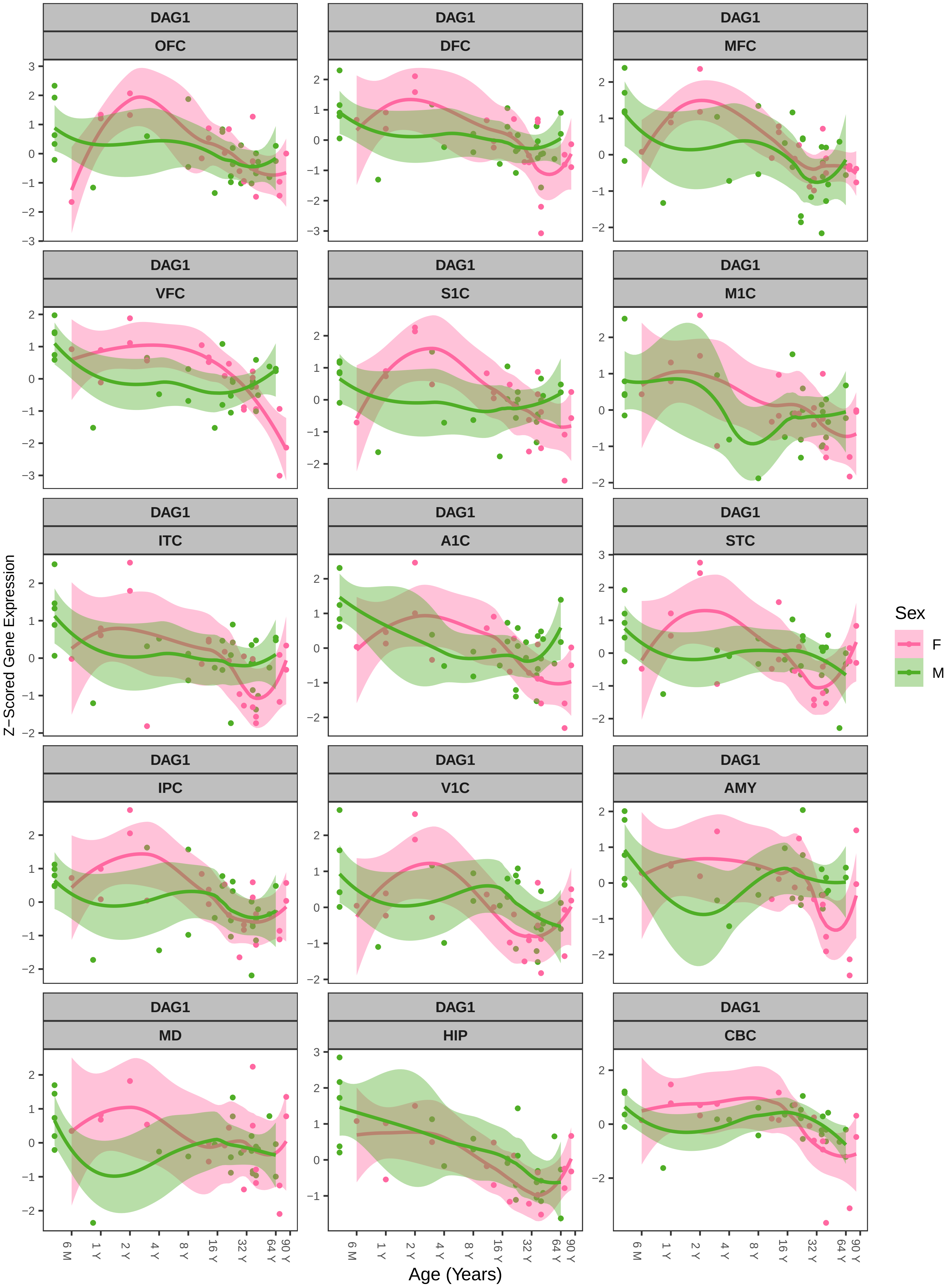
DAG1
OFC
DFC
MFC
VFC
S1C
M1C
ITC
A1C
STC
IPC
V1C
AMY
MD
HIP
CBC
Z−Scored Gene Expression
Age (Years)
6 M
1 Y
2 Y
4 Y
8 Y
16 Y
32 Y
64 Y
90 Y
Sex
F
M

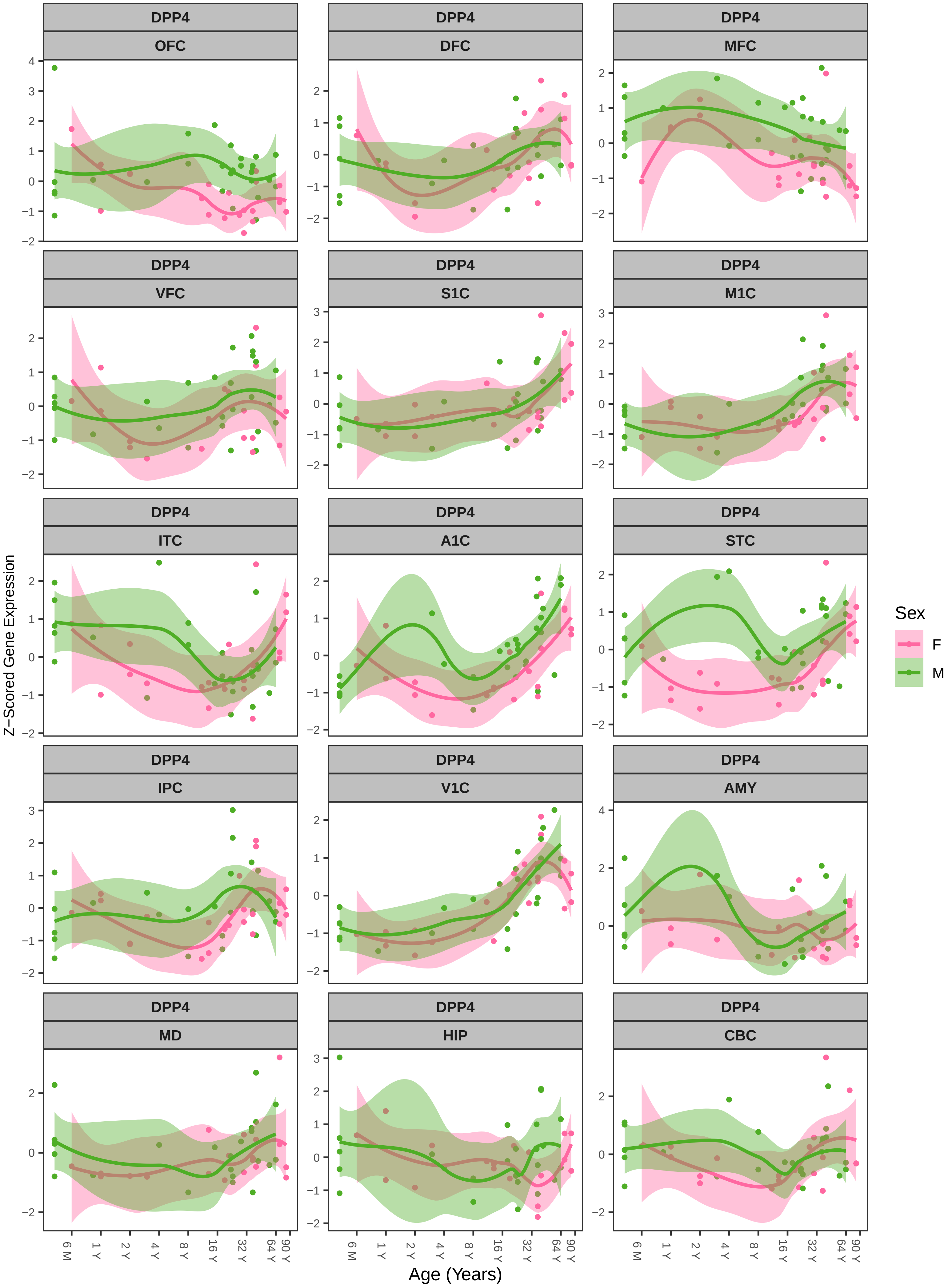
DPP4
OFC
DFC
MFC
VFC
S1C
M1C
ITC
A1C
STC
IPC
V1C
AMY
MD
HIP
CBC
Z−Scored Gene Expression
Age (Years)
6 M
1 Y
2 Y
4 Y
8 Y
16 Y
32 Y
64 Y
90 Y
Sex
F
M

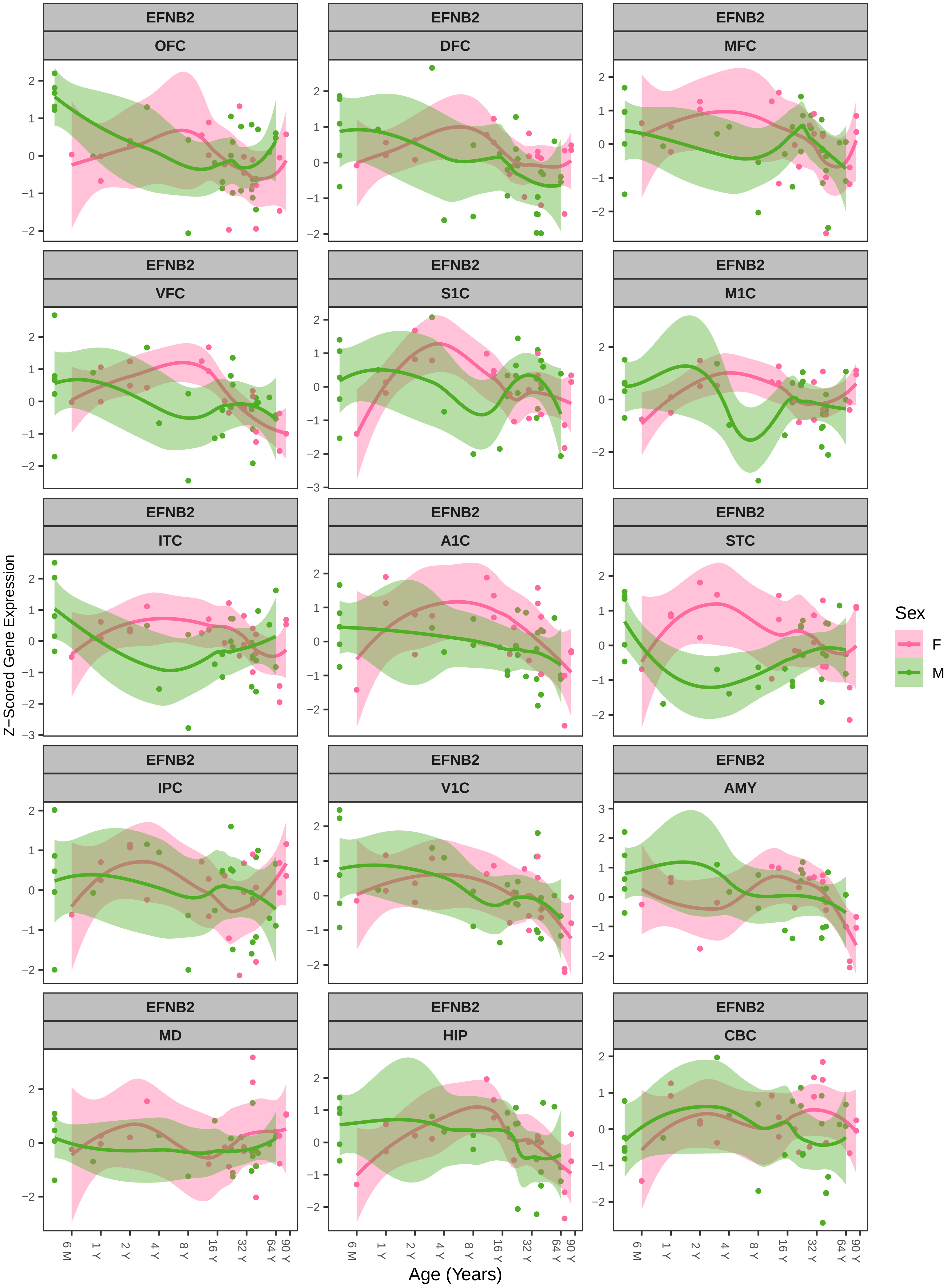
EFNB2
OFC
DFC
MFC
VFC
S1C
M1C
ITC
A1C
STC
IPC
V1C
AMY
MD
HIP
CBC
Z−Scored Gene Expression
Age (Years)
6 M
1 Y
2 Y
4 Y
8 Y
16 Y
32 Y
64 Y
90 Y
Sex
F
M

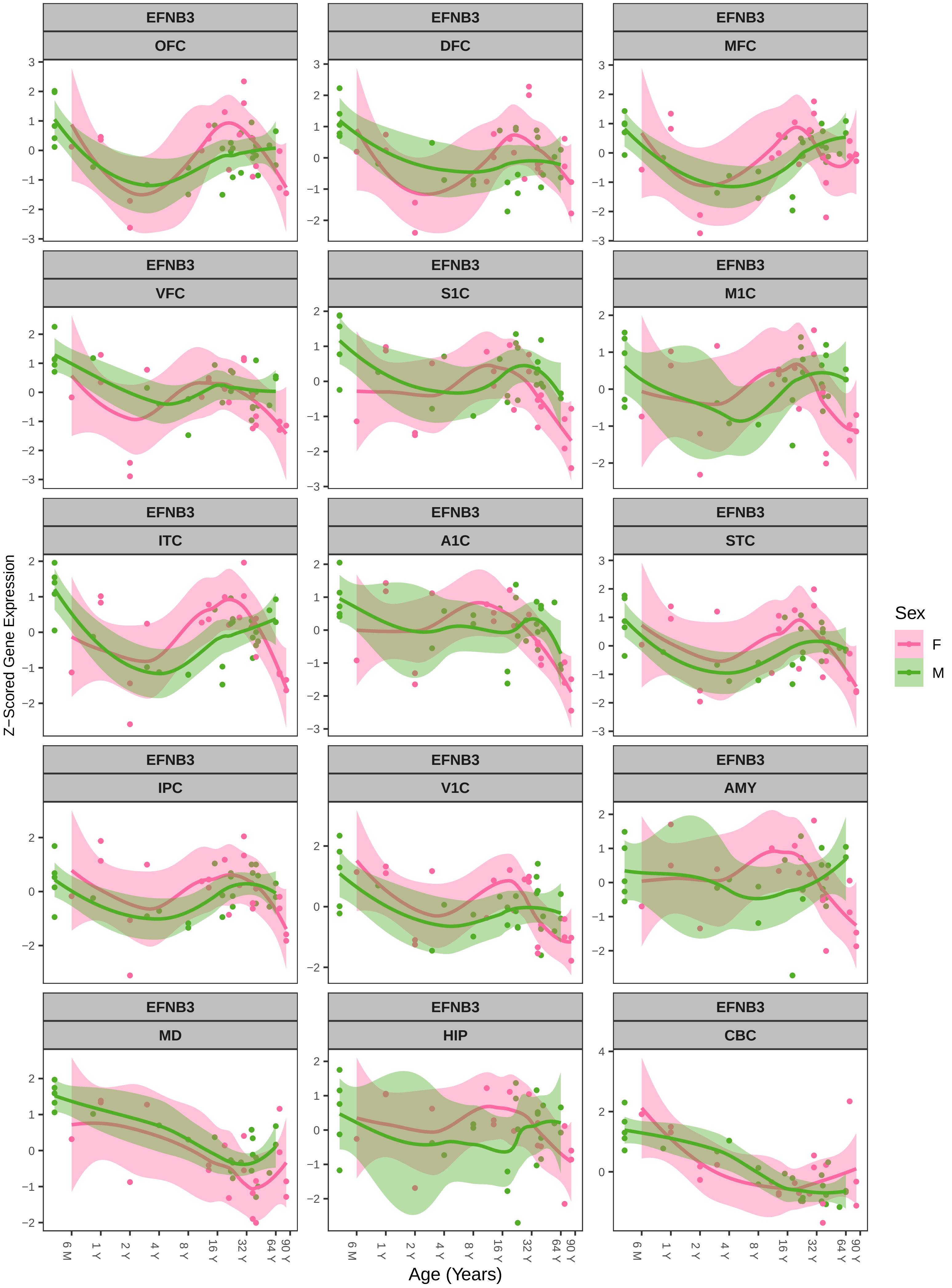
EFNB3
OFC
DFC
MFC
VFC
S1C
M1C
ITC
A1C
STC
IPC
V1C
AMY
MD
HIP
CBC
Z−Scored Gene Expression
Age (Years)
6 M
1 Y
2 Y
4 Y
8 Y
16 Y
32 Y
64 Y
90 Y
Sex
F
M

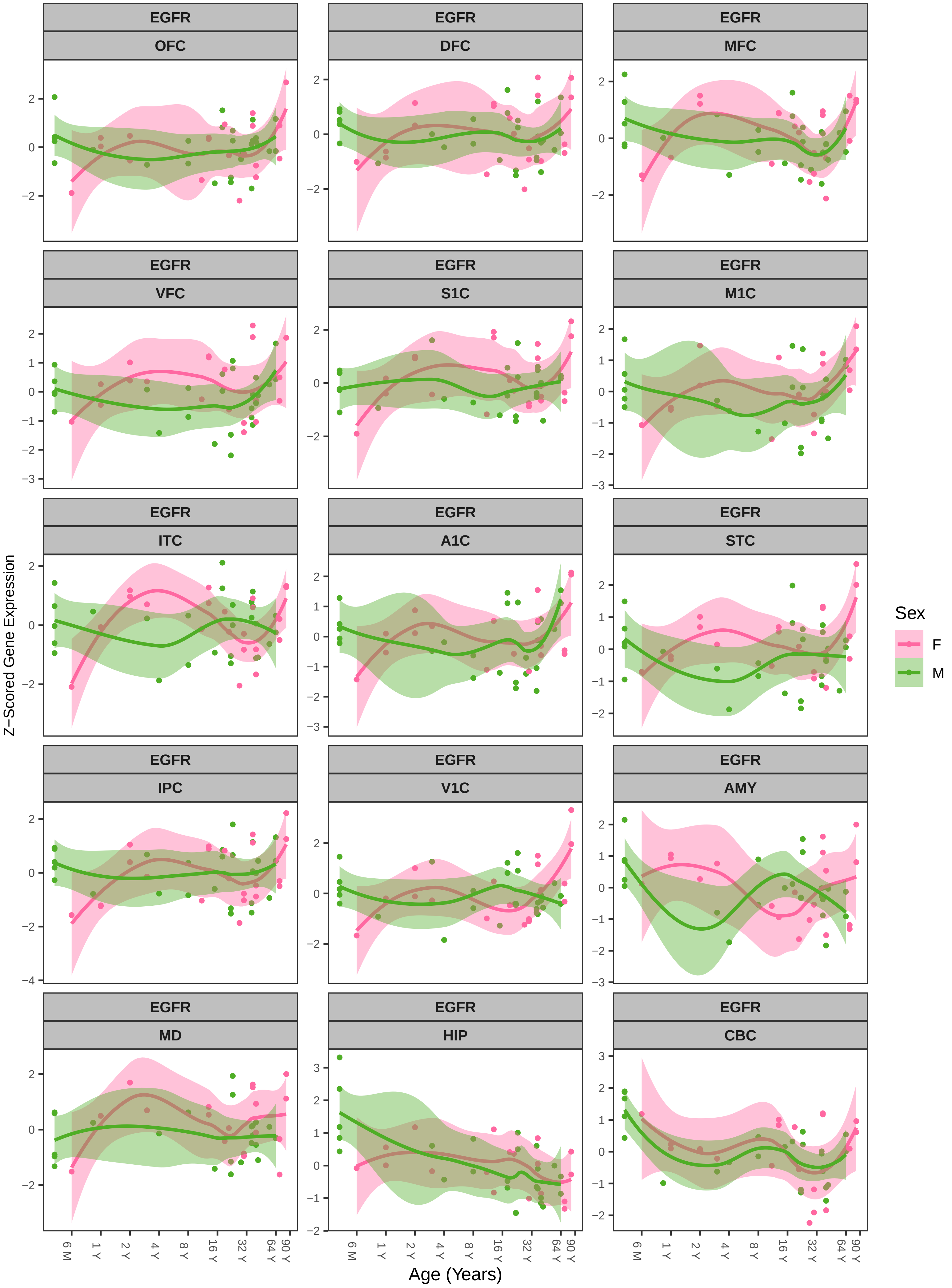
EGFR
OFC
DFC
MFC
VFC
S1C
M1C
ITC
A1C
STC
IPC
V1C
AMY
MD
HIP
CBC
Z−Scored Gene Expression
Age (Years)
6 M
1 Y
2 Y
4 Y
8 Y
16 Y
32 Y
64 Y
90 Y
Sex
F
M

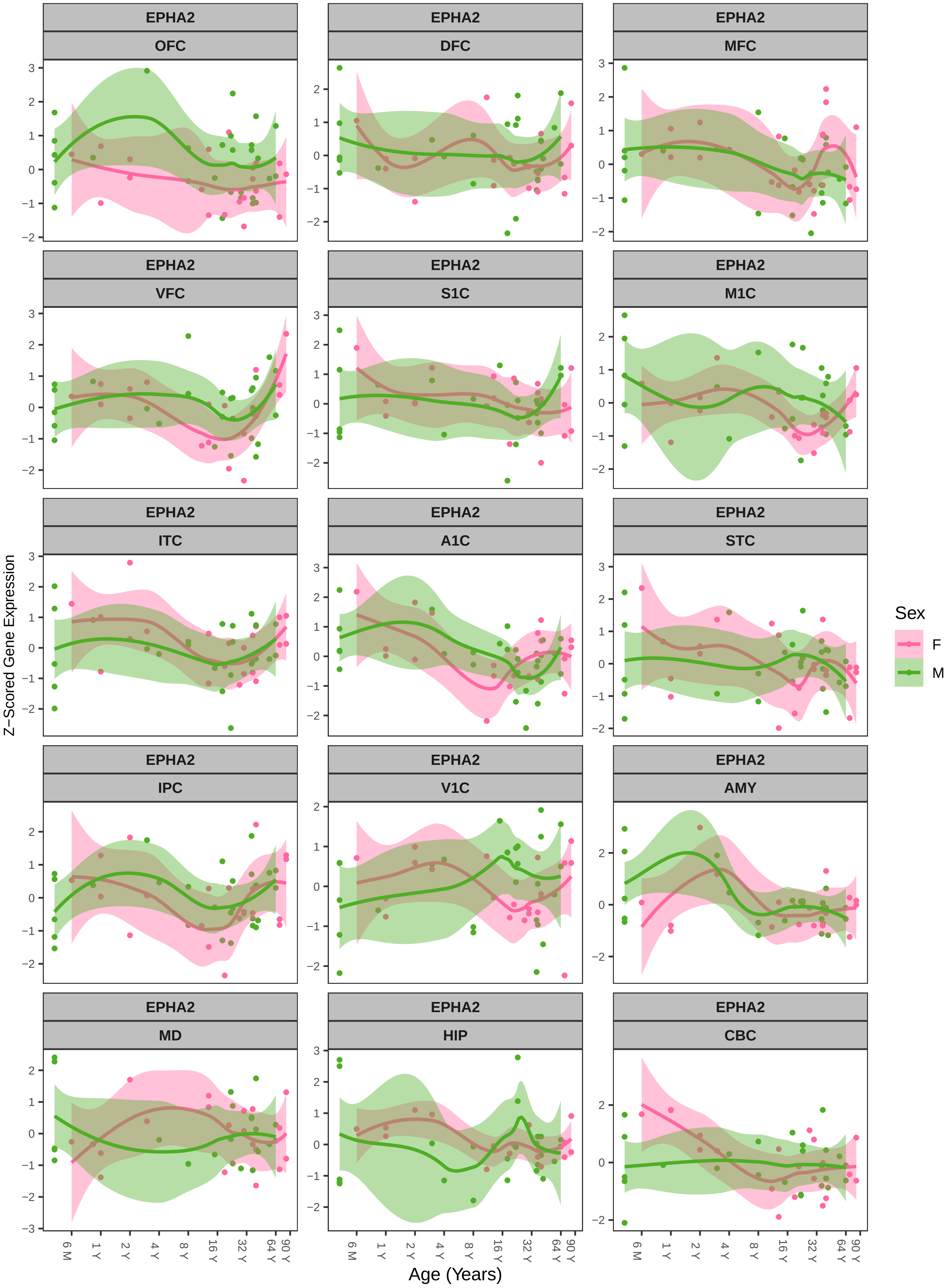

EPHA2
OFC
DFC
MFC
VFC
S1C
M1C
ITC
A1C
STC
IPC
V1C
AMY
MD
HIP
CBC
Z−Scored Gene Expression
Age (Years)
6 M
1 Y
2 Y
4 Y
8 Y
16 Y
32 Y
64 Y
90 Y
Sex
F
M

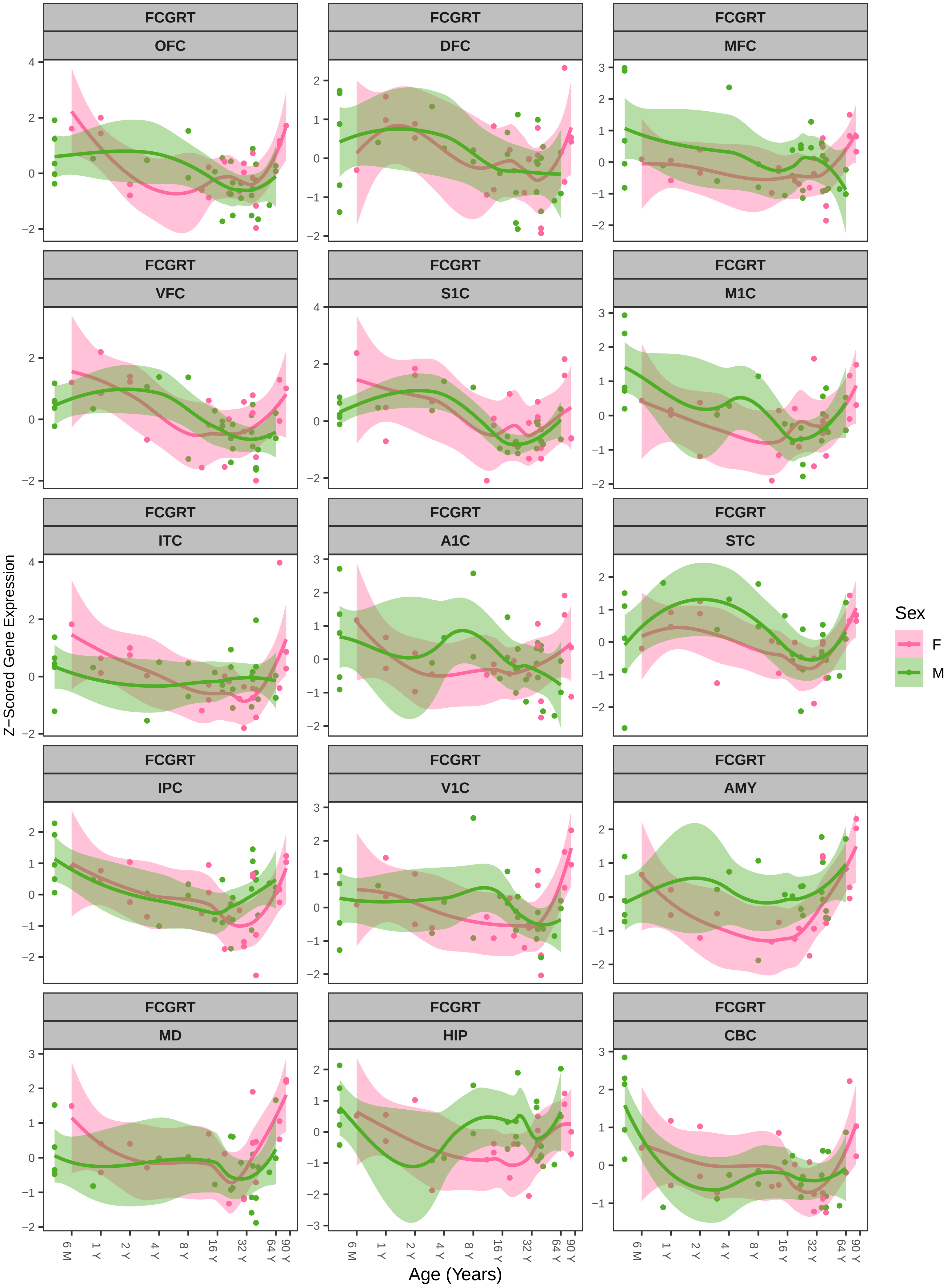
FCGRT
OFC
DFC
MFC
VFC
S1C
M1C
ITC
A1C
STC
IPC
V1C
AMY
MD
HIP
CBC
Z−Scored Gene Expression
Age (Years)
6 M
1 Y
2 Y
4 Y
8 Y
16 Y
32 Y
64 Y
90 Y
Sex
F
M

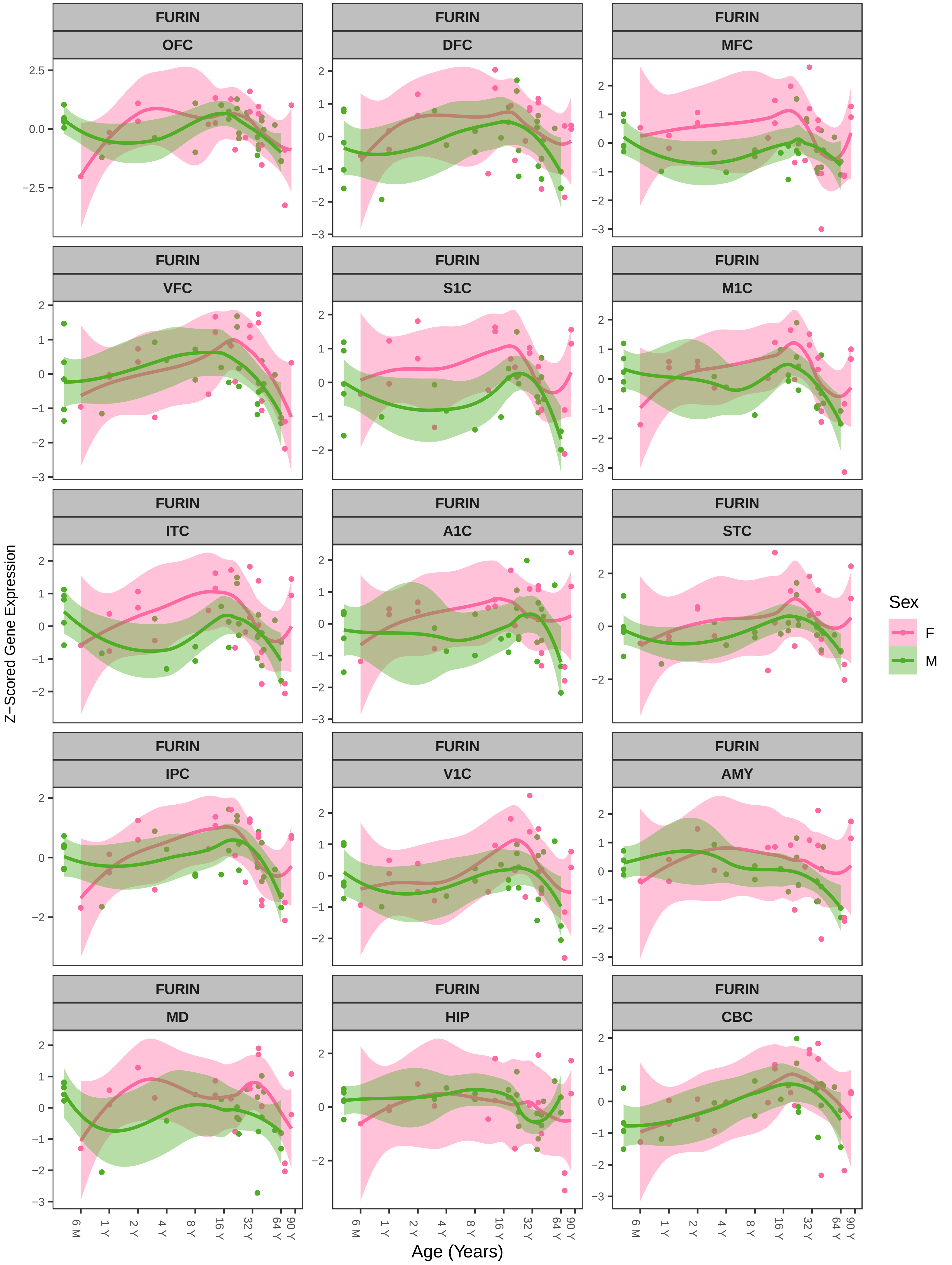

FURIN
OFC
DFC
MFC
VFC
S1C
M1C
ITC
A1C
STC
IPC
V1C
AMY
MD
HIP
CBC
Z−Scored Gene Expression
Age (Years)
6 M
1 Y
2 Y
4 Y
8 Y
16 Y
32 Y
64 Y
90 Y
Sex
F
M

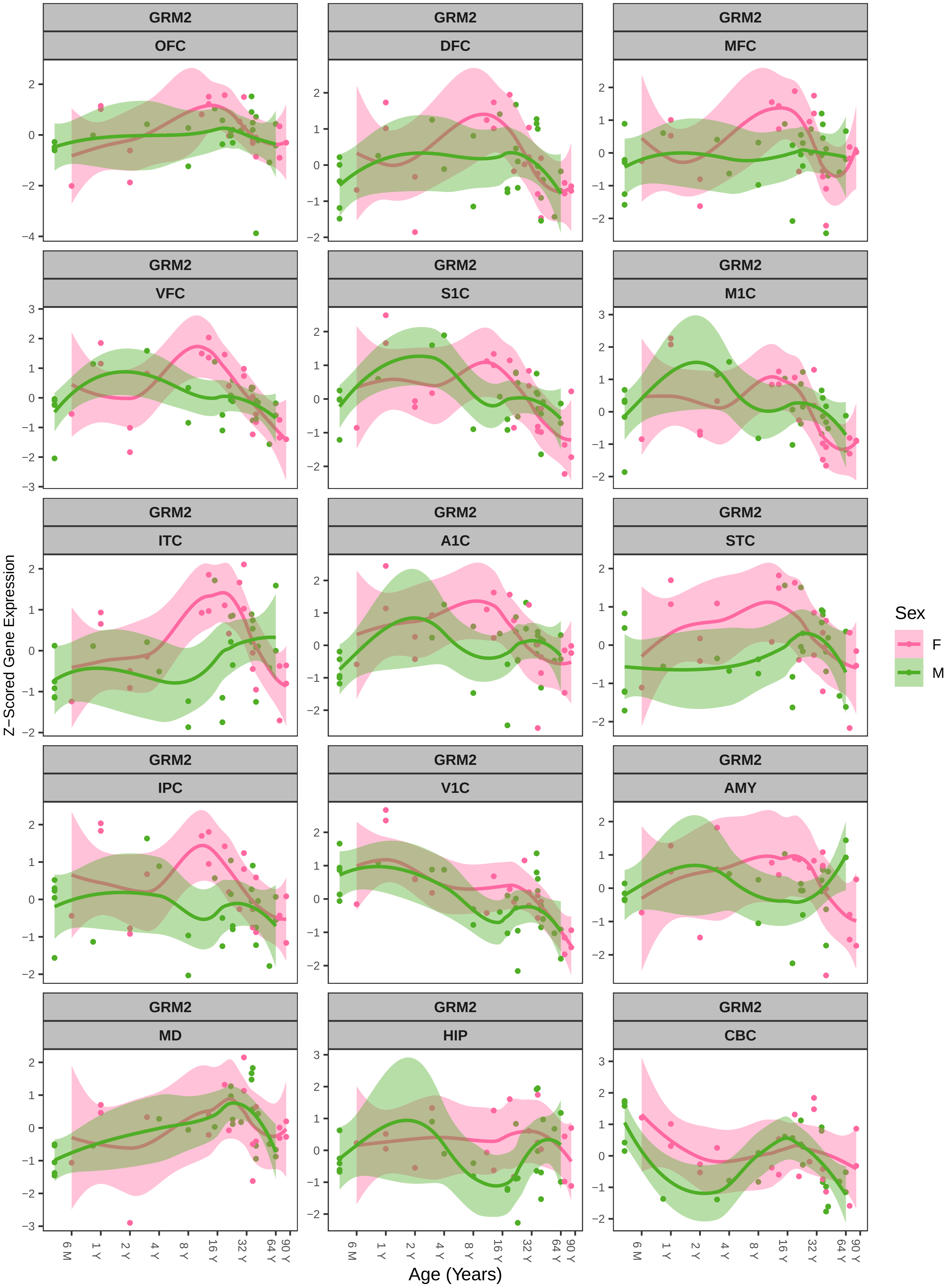
GRM2
OFC
DFC
MFC
VFC
S1C
M1C
ITC
A1C
STC
IPC
V1C
AMY
MD
HIP
CBC
Z−Scored Gene Expression
Age (Years)
6 M
1 Y
2 Y
4 Y
8 Y
16 Y
32 Y
64 Y
90 Y
Sex
F
M

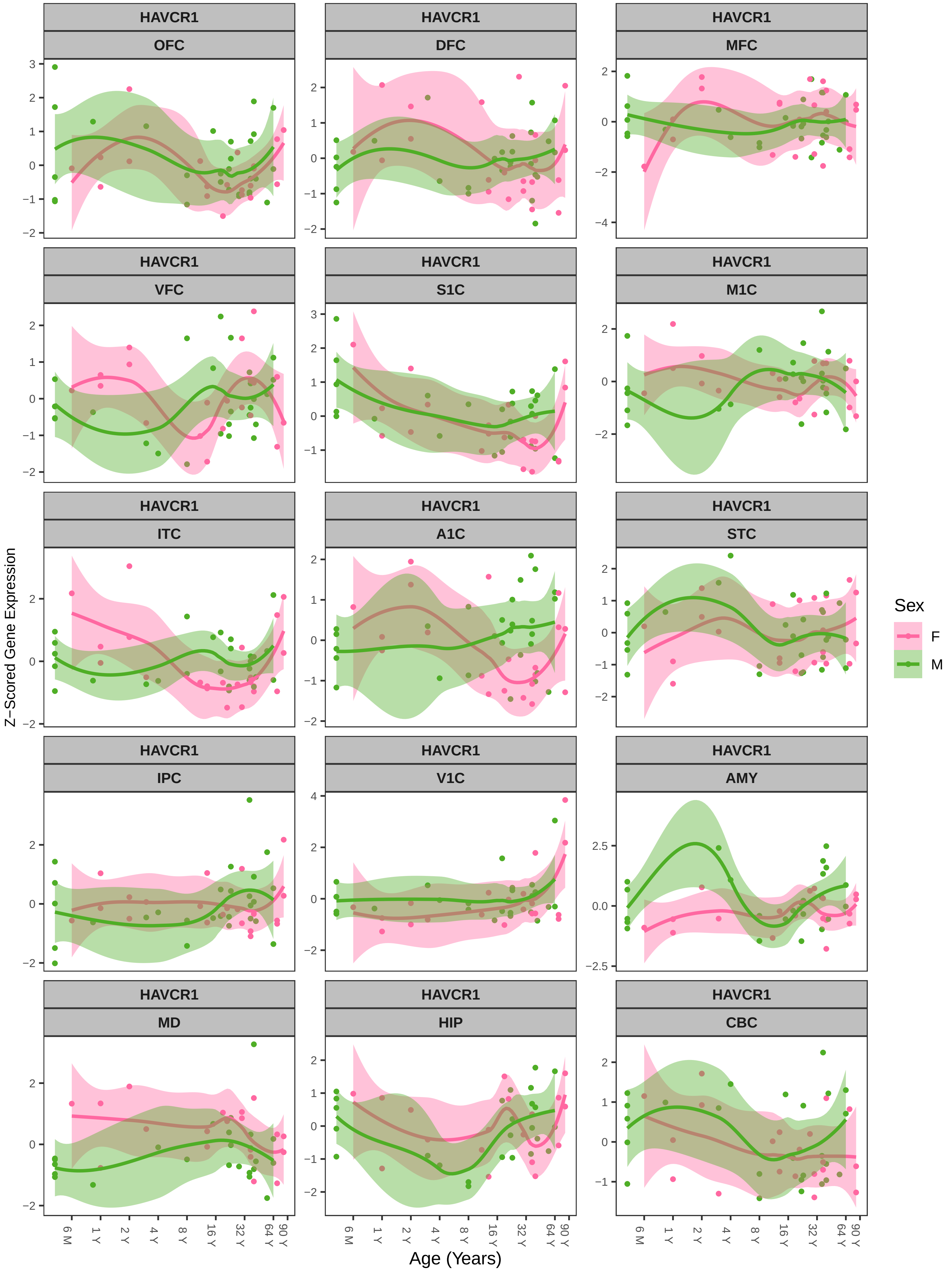
HAVCR1
OFC
DFC
MFC
VFC
S1C
M1C
ITC
A1C
STC
IPC
V1C
AMY
MD
HIP
CBC
Z−Scored Gene Expression
Age (Years)
6 M
1 Y
2 Y
4 Y
8 Y
16 Y
32 Y
64 Y
90 Y
Sex
F
M

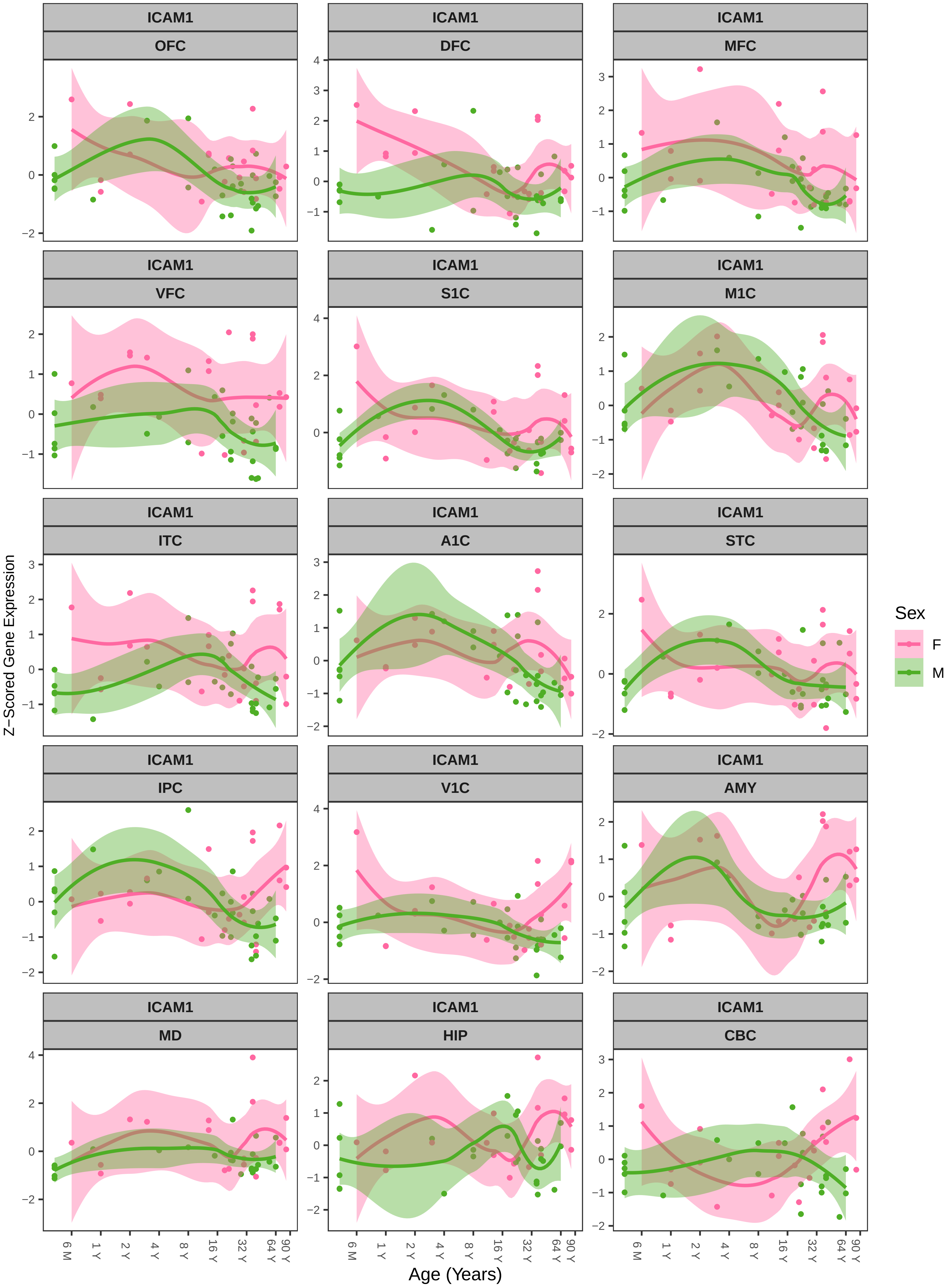
ICAM1
OFC
DFC
MFC
VFC
S1C
M1C
ITC
A1C
STC
IPC
V1C
AMY
MD
HIP
CBC
Z−Scored Gene Expression
Age (Years)
6 M
1 Y
2 Y
4 Y
8 Y
16 Y
32 Y
64 Y
90 Y
Sex
F
M

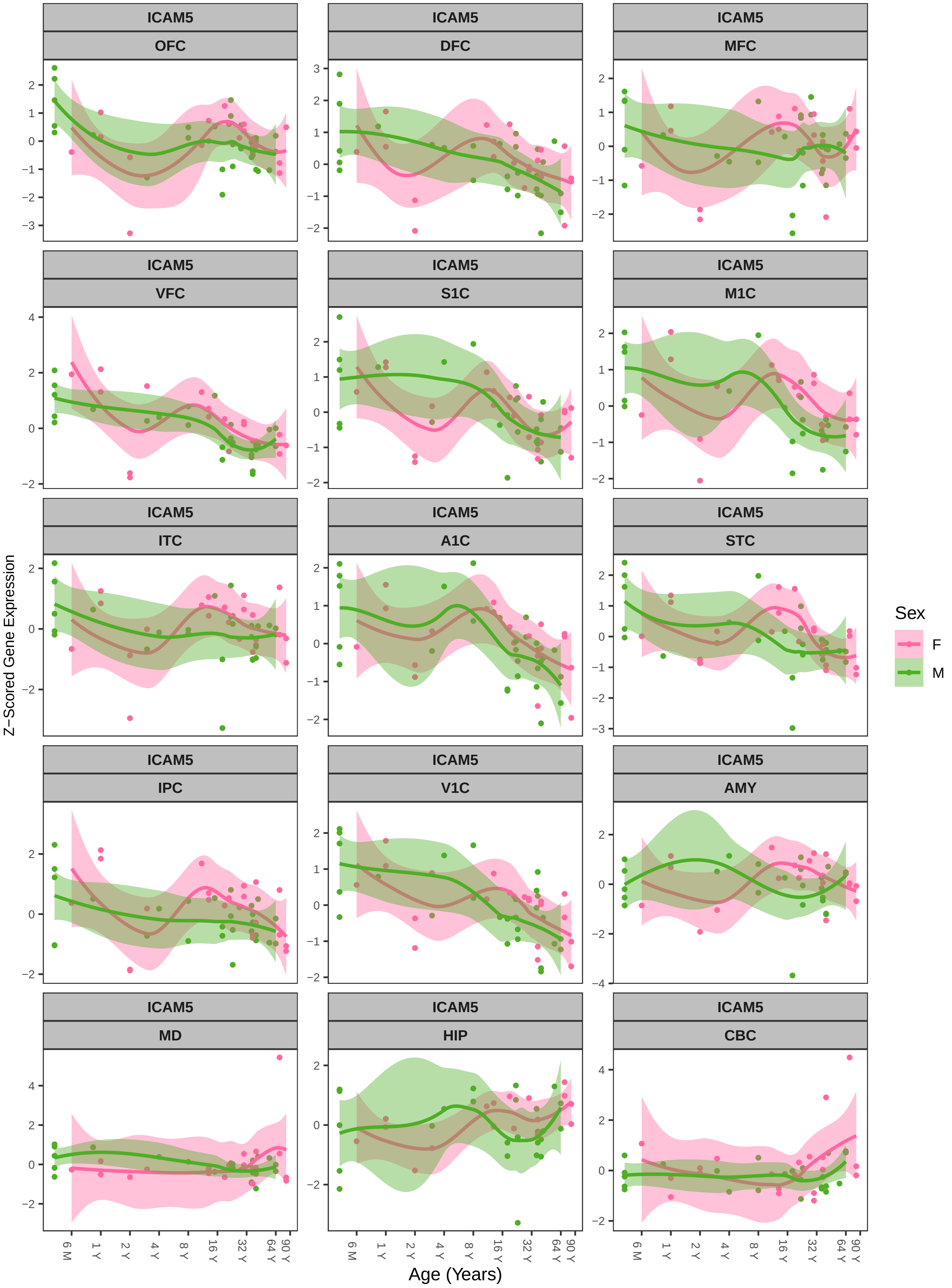
ICAM5
OFC
DFC
MFC
VFC
S1C
M1C
ITC
A1C
STC
IPC
V1C
AMY
MD
HIP
CBC
Z−Scored Gene Expression
Age (Years)
6 M
1 Y
2 Y
4 Y
8 Y
16 Y
32 Y
64 Y
90 Y
Sex
F
M

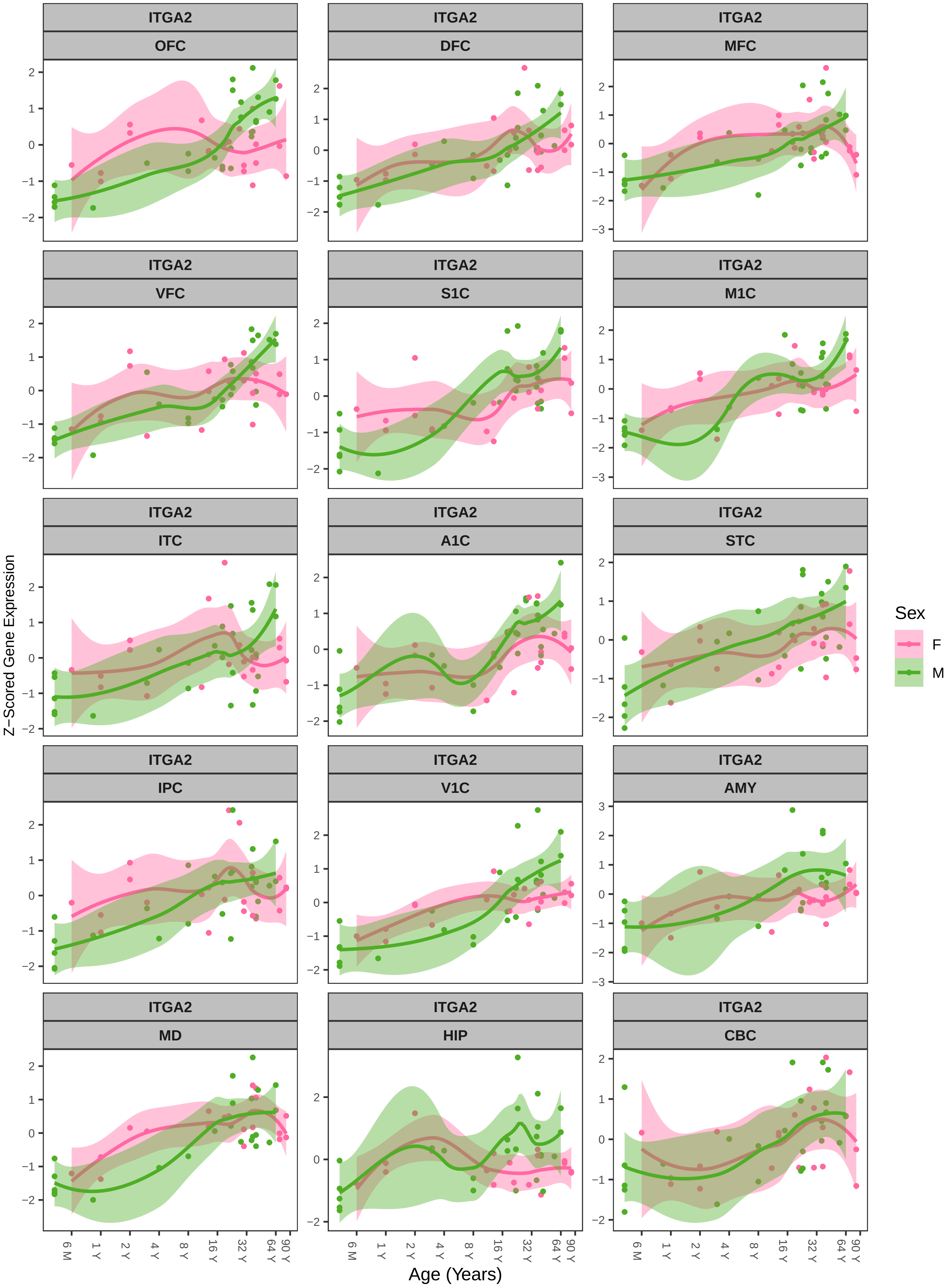
ITGA2
OFC
DFC
MFC
VFC
S1C
M1C
ITC
A1C
STC
IPC
V1C
AMY
MD
HIP
CBC
Z−Scored Gene Expression
Age (Years)
6 M
1 Y
2 Y
4 Y
8 Y
16 Y
32 Y
64 Y
90 Y
Sex
F
M

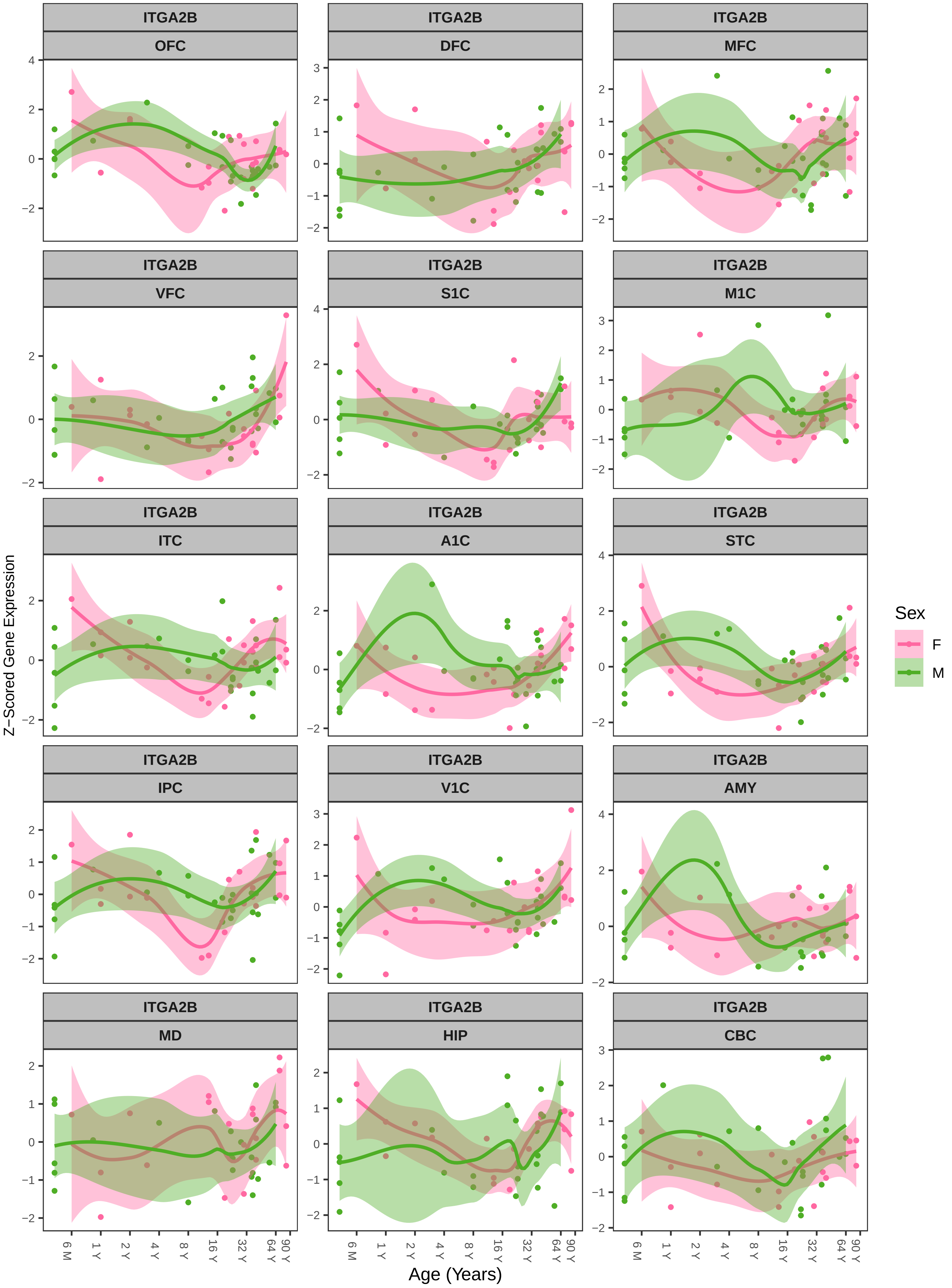

ITGA2B
OFC
DFC
MFC
VFC
S1C
M1C
ITC
A1C
STC
IPC
V1C
AMY
MD
HIP
CBC
Z−Scored Gene Expression
Age (Years)
6 M
1 Y
2 Y
4 Y
8 Y
16 Y
32 Y
64 Y
90 Y
Sex
F
M

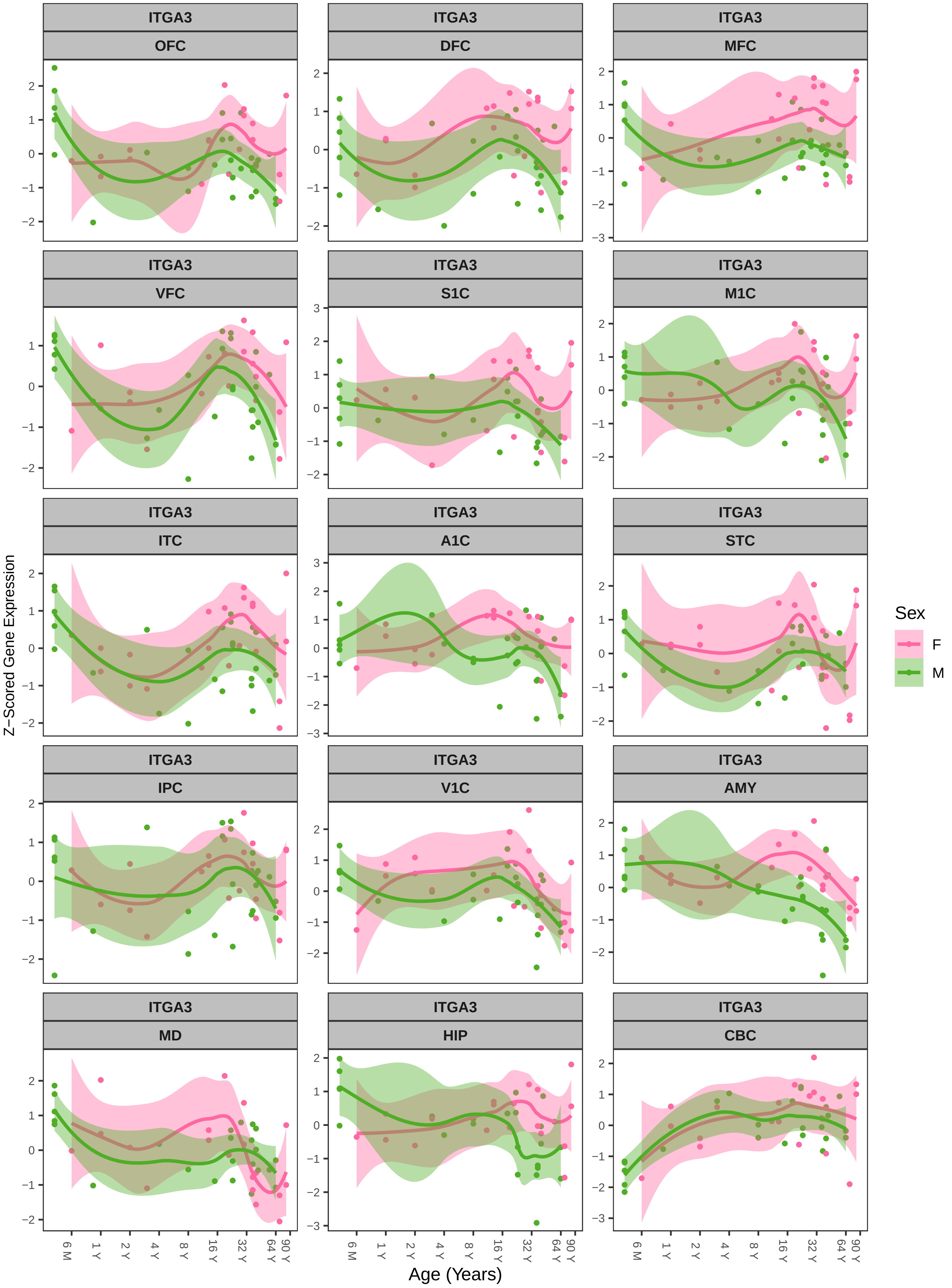

ITGA3
OFC
DFC
MFC
VFC
S1C
M1C
ITC
A1C
STC
IPC
V1C
AMY
MD
HIP
CBC
Z−Scored Gene Expression
Age (Years)
6 M
1 Y
2 Y
4 Y
8 Y
16 Y
32 Y
64 Y
90 Y
Sex
F
M

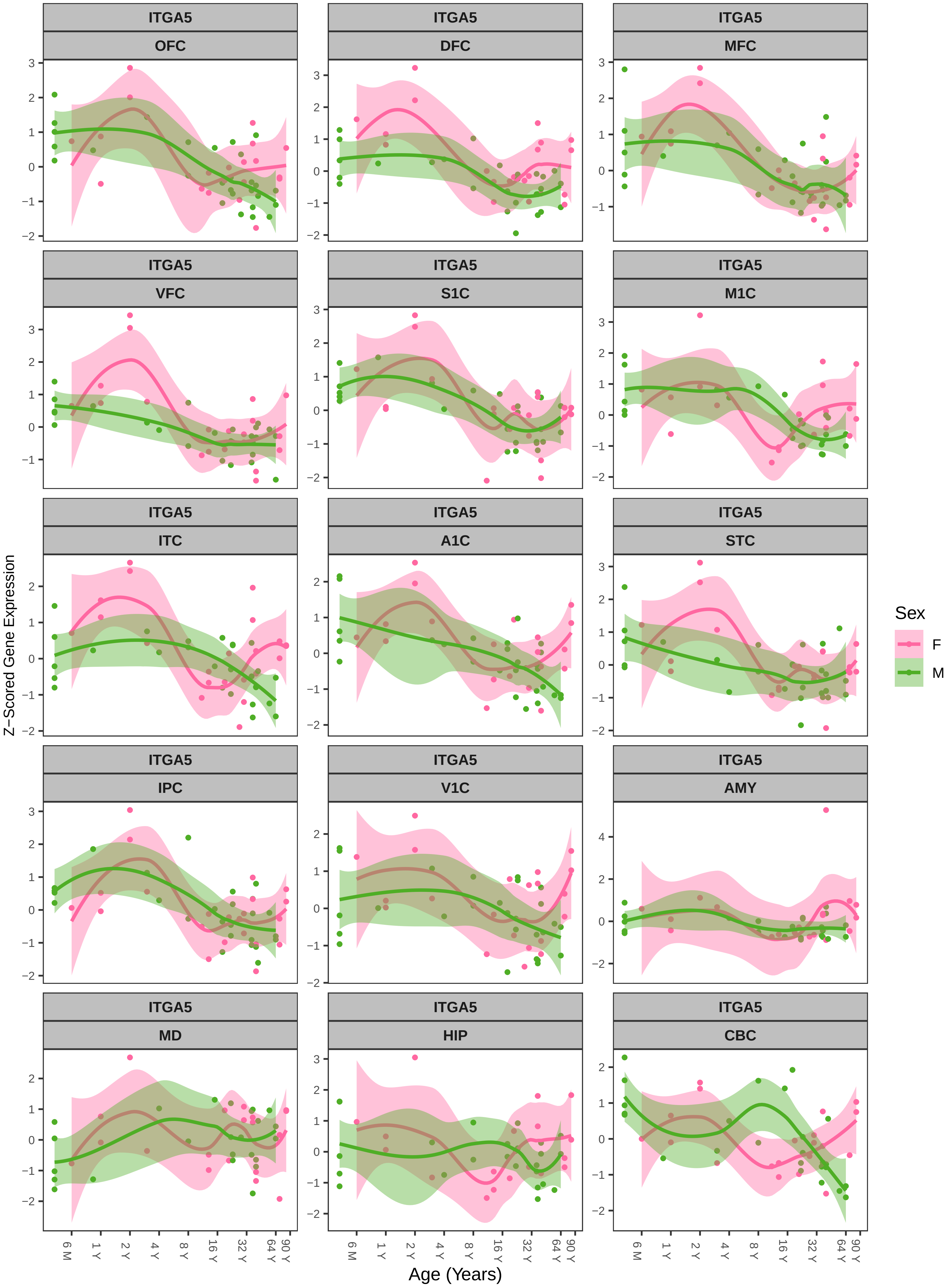

ITGA5
OFC
DFC
MFC
VFC
S1C
M1C
ITC
A1C
STC
IPC
V1C
AMY
MD
HIP
CBC
Z−Scored Gene Expression
Age (Years)
6 M
1 Y
2 Y
4 Y
8 Y
16 Y
32 Y
64 Y
90 Y
Sex
F
M

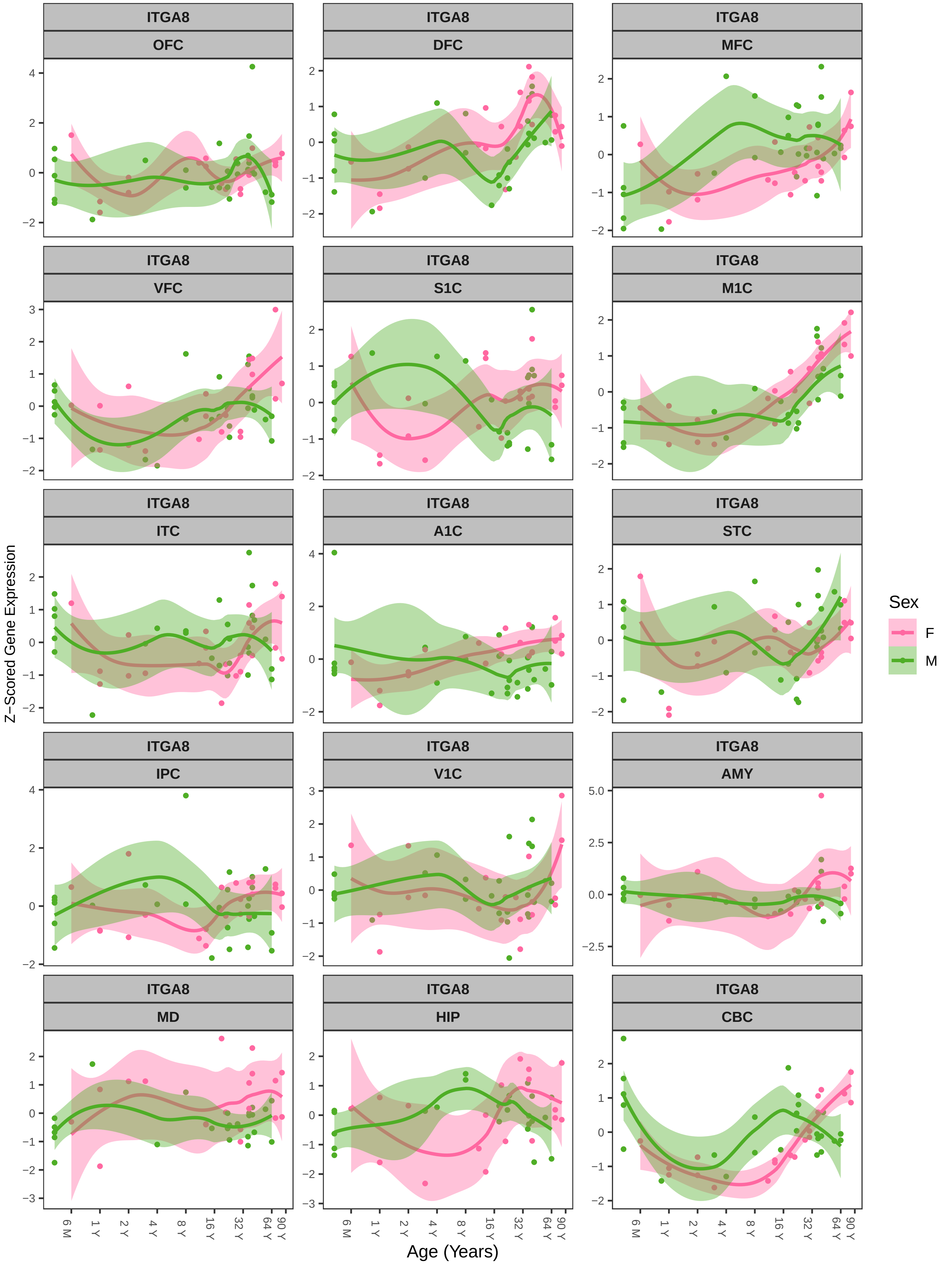

ITGA8
OFC
ITGA8
DFC
ITGA8
MFC
ITGA8
VFC
ITGA8
S1C
ITGA8
M1C
ITGA8
ITC
ITGA8
A1C
ITGA8
STC
ITGA8
IPC
ITGA8
V1C
ITGA8
AMY
ITGA8
MD
ITGA8
HIP
ITGA8
CBC
Z−Scored Gene Expression
Age (Years)
6 M
1 Y
2 Y
4 Y
8 Y
16 Y
32 Y
64 Y
90 Y
Sex
F
M

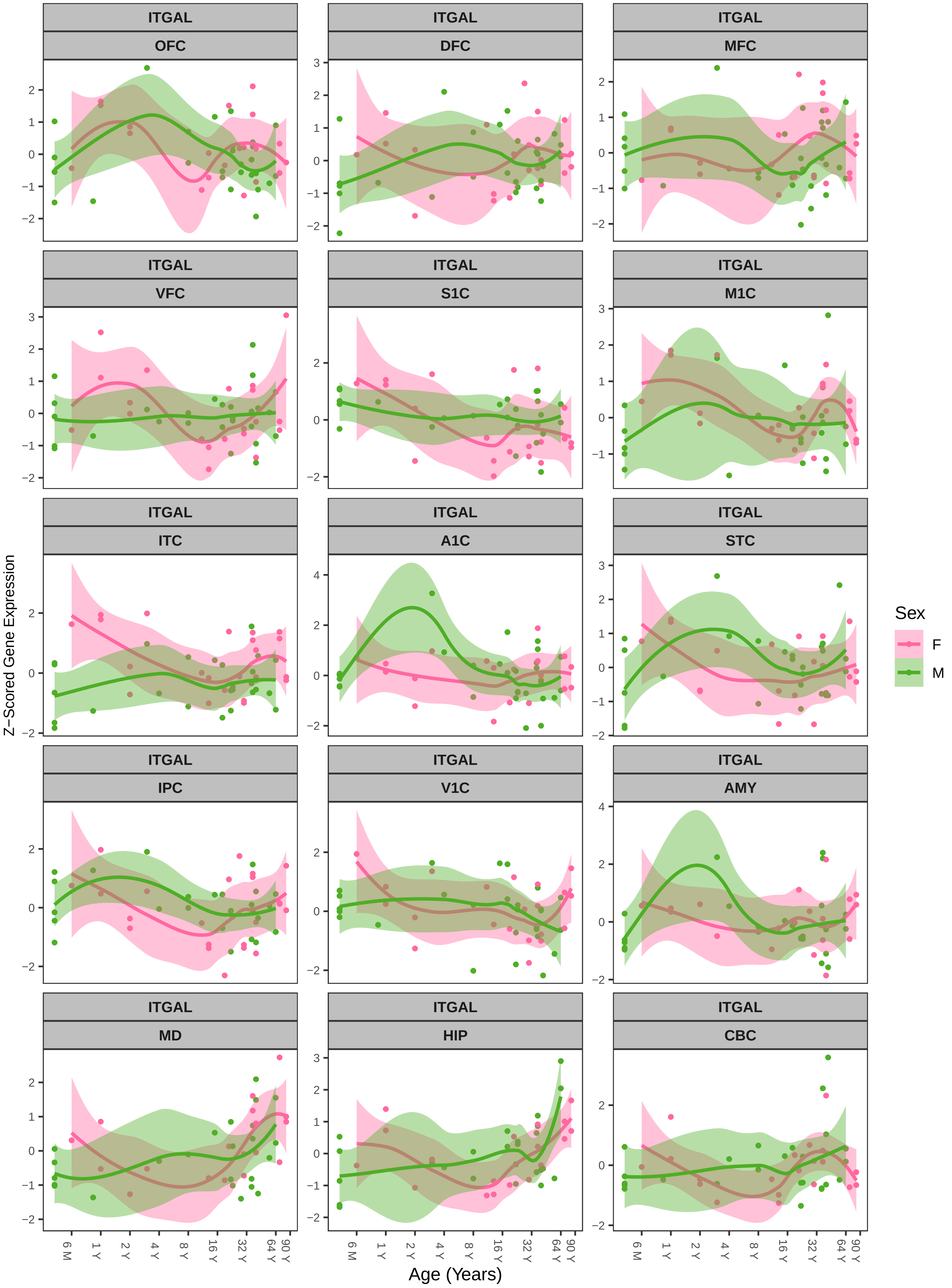

ITGAL
OFC
DFC
MFC
VFC
S1C
M1C
ITC
A1C
STC
IPC
V1C
AMY
MD
HIP
CBC
Z−Scored Gene Expression
Age (Years)
6 M
1 Y
2 Y
4 Y
8 Y
16 Y
32 Y
64 Y
90 Y
Sex
F
M

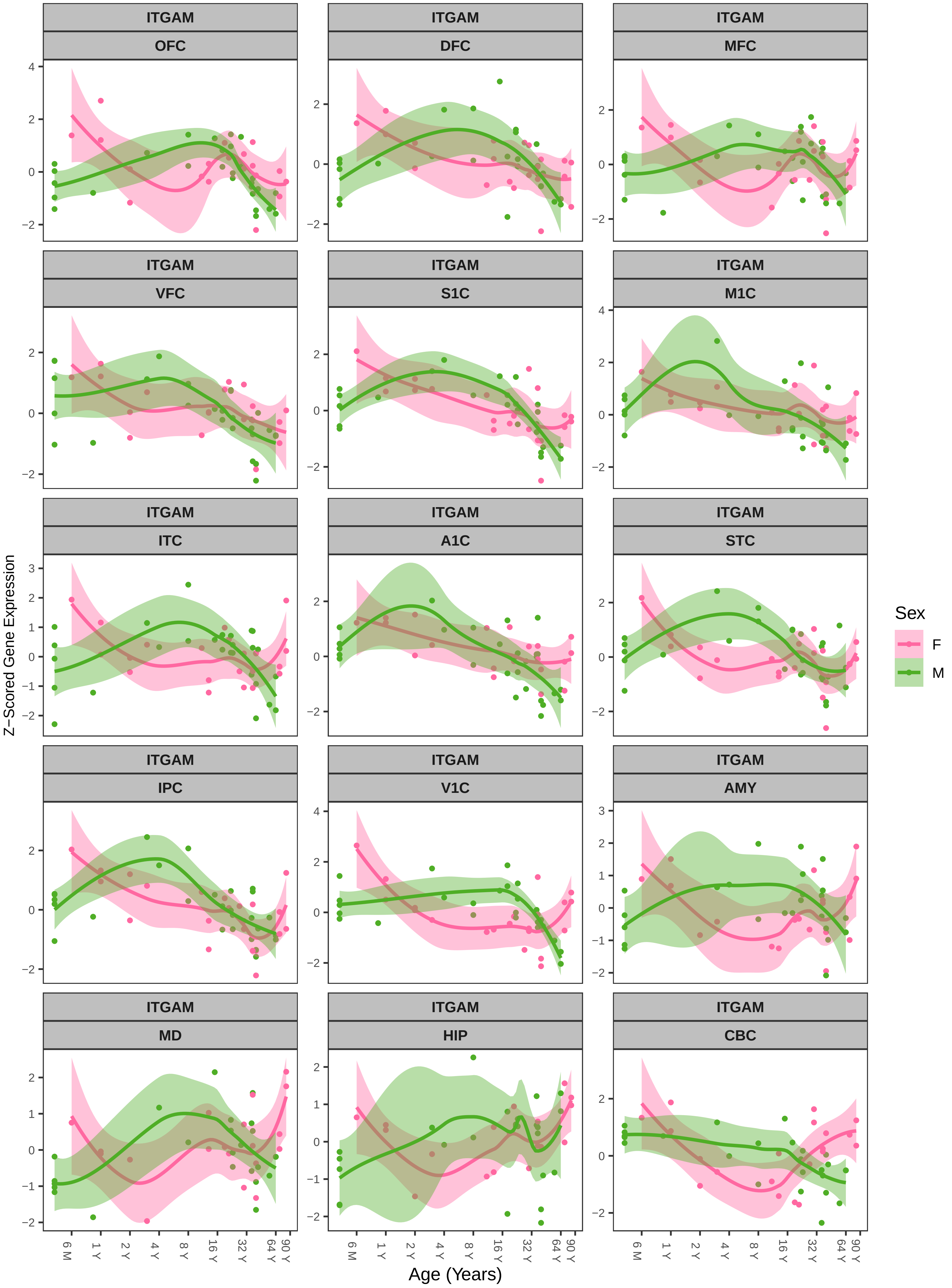

ITGAM
OFC
DFC
MFC
VFC
S1C
M1C
ITC
A1C
STC
IPC
V1C
AMY
MD
HIP
CBC
Z−Scored Gene Expression
Age (Years)
6 M
1 Y
2 Y
4 Y
8 Y
16 Y
32 Y
64 Y
90 Y
Sex
F
M

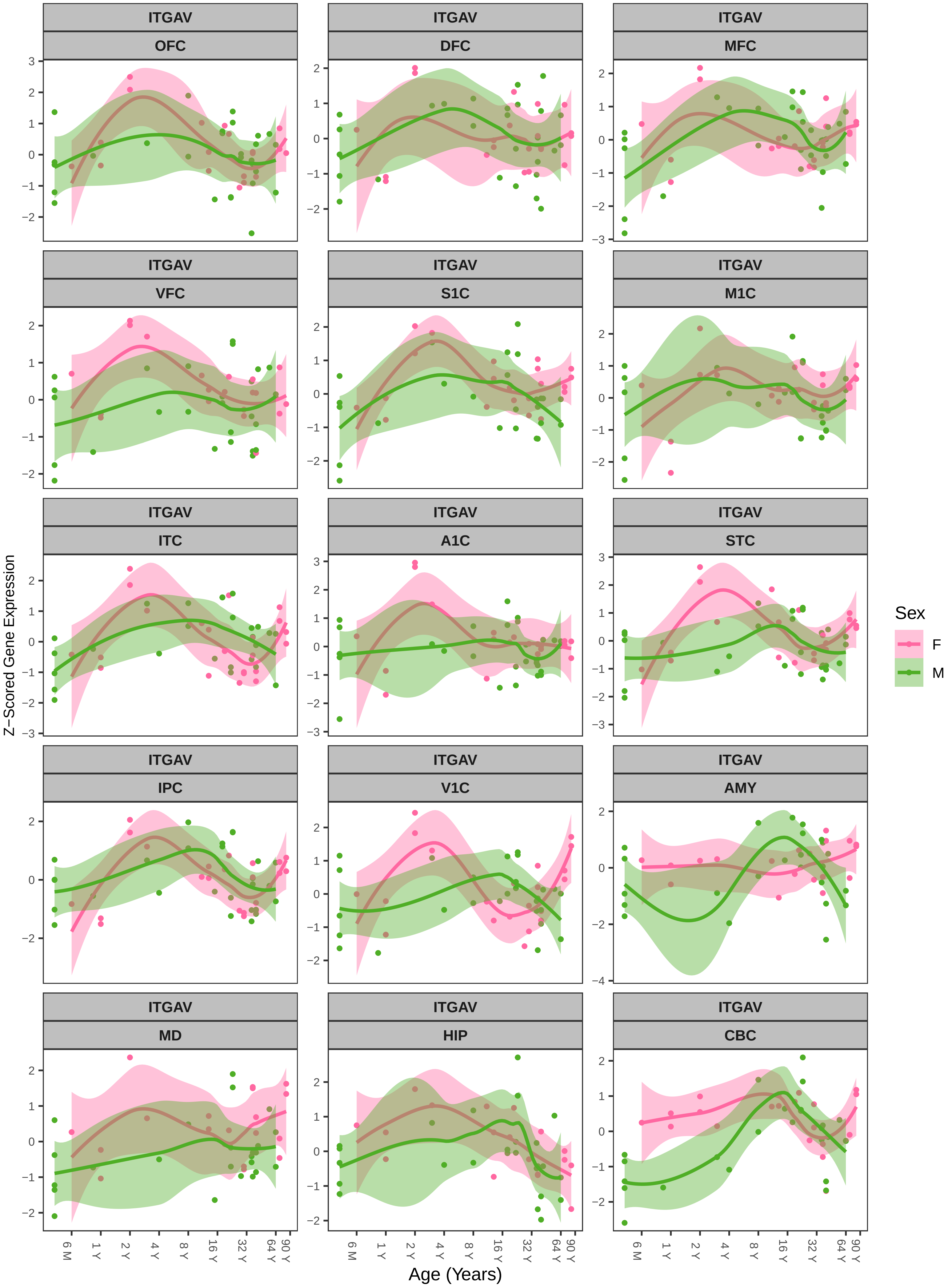

ITGAV
OFC
DFC
MFC
VFC
S1C
M1C
ITC
A1C
STC
IPC
V1C
AMY
MD
HIP
CBC
Z−Scored Gene Expression
Age (Years)
6 M
1 Y
2 Y
4 Y
8 Y
16 Y
32 Y
64 Y
90 Y
Sex
F
M

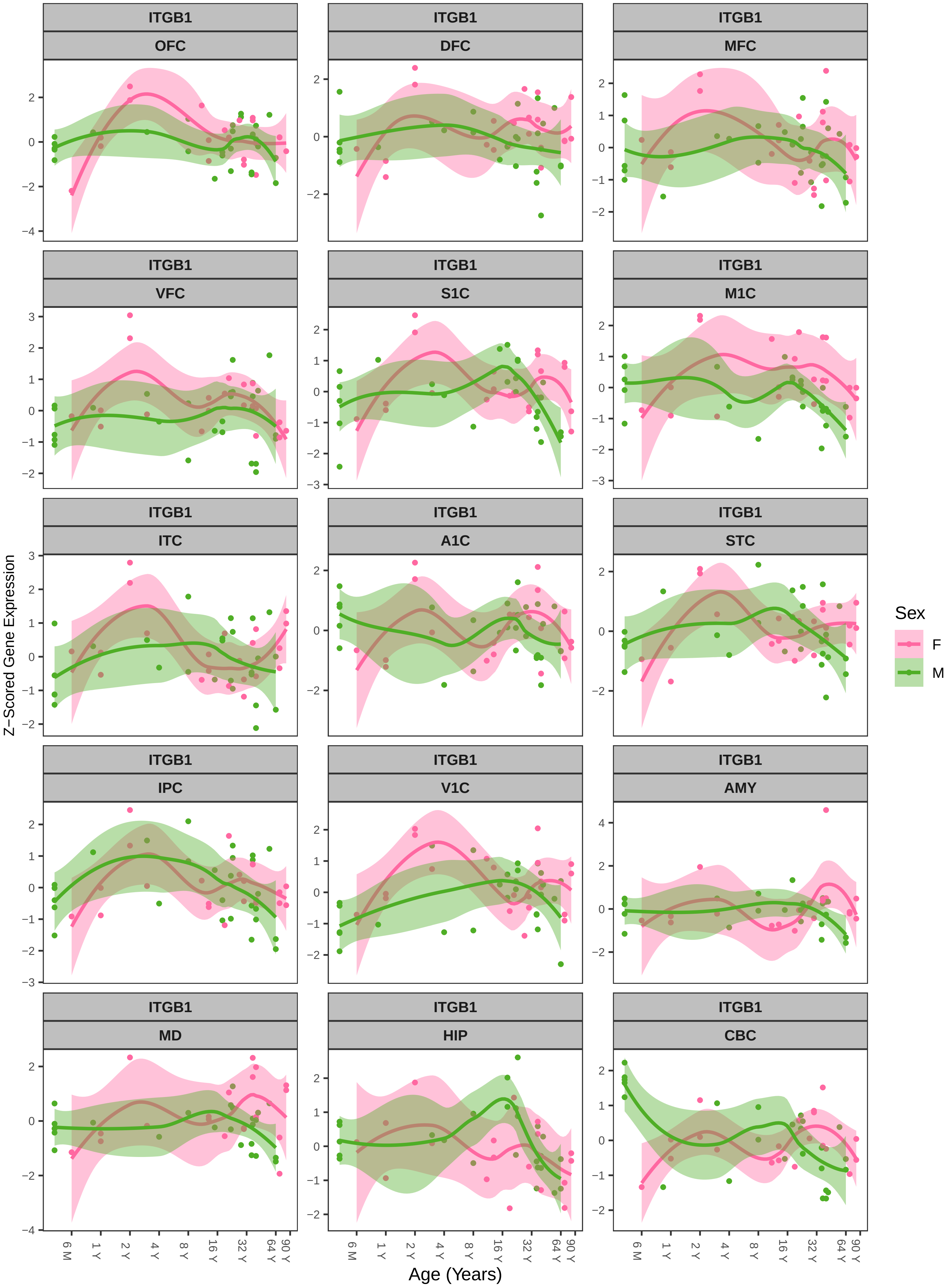
ITGB1
OFC
DFC
MFC
VFC
S1C
M1C
ITC
A1C
STC
IPC
V1C
AMY
MD
HIP
CBC
Z−Scored Gene Expression
Age (Years)
6 M
1 Y
2 Y
4 Y
8 Y
16 Y
32 Y
64 Y
90 Y
Sex
F
M

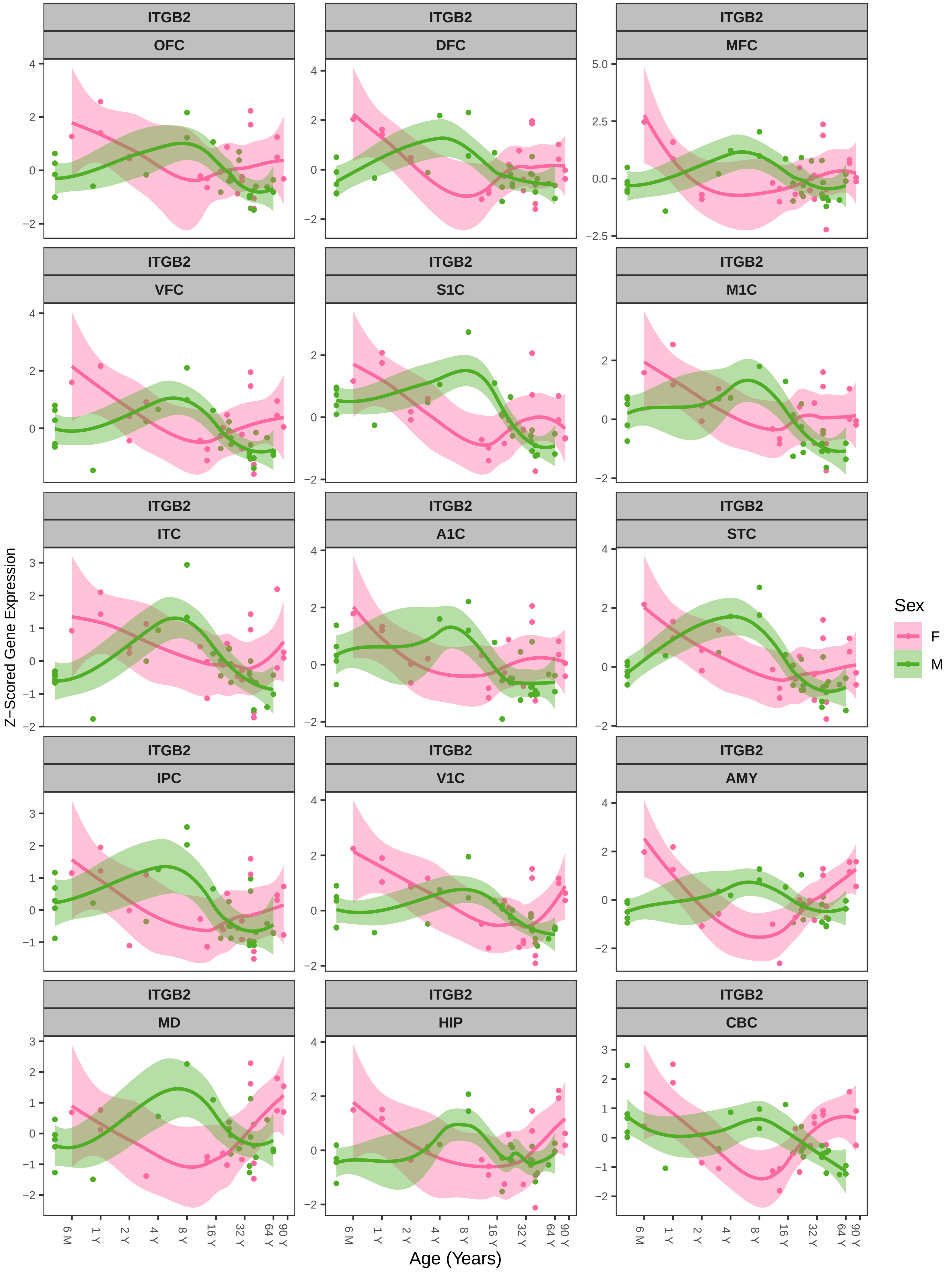
ITGB2
OFC
DFC
MFC
VFC
S1C
M1C
ITC
A1C
STC
IPC
V1C
AMY
MD
HIP
CBC
Z−Scored Gene Expression
Age (Years)
6 M
1 Y
2 Y
4 Y
8 Y
16 Y
32 Y
64 Y
90 Y
Sex
F
M

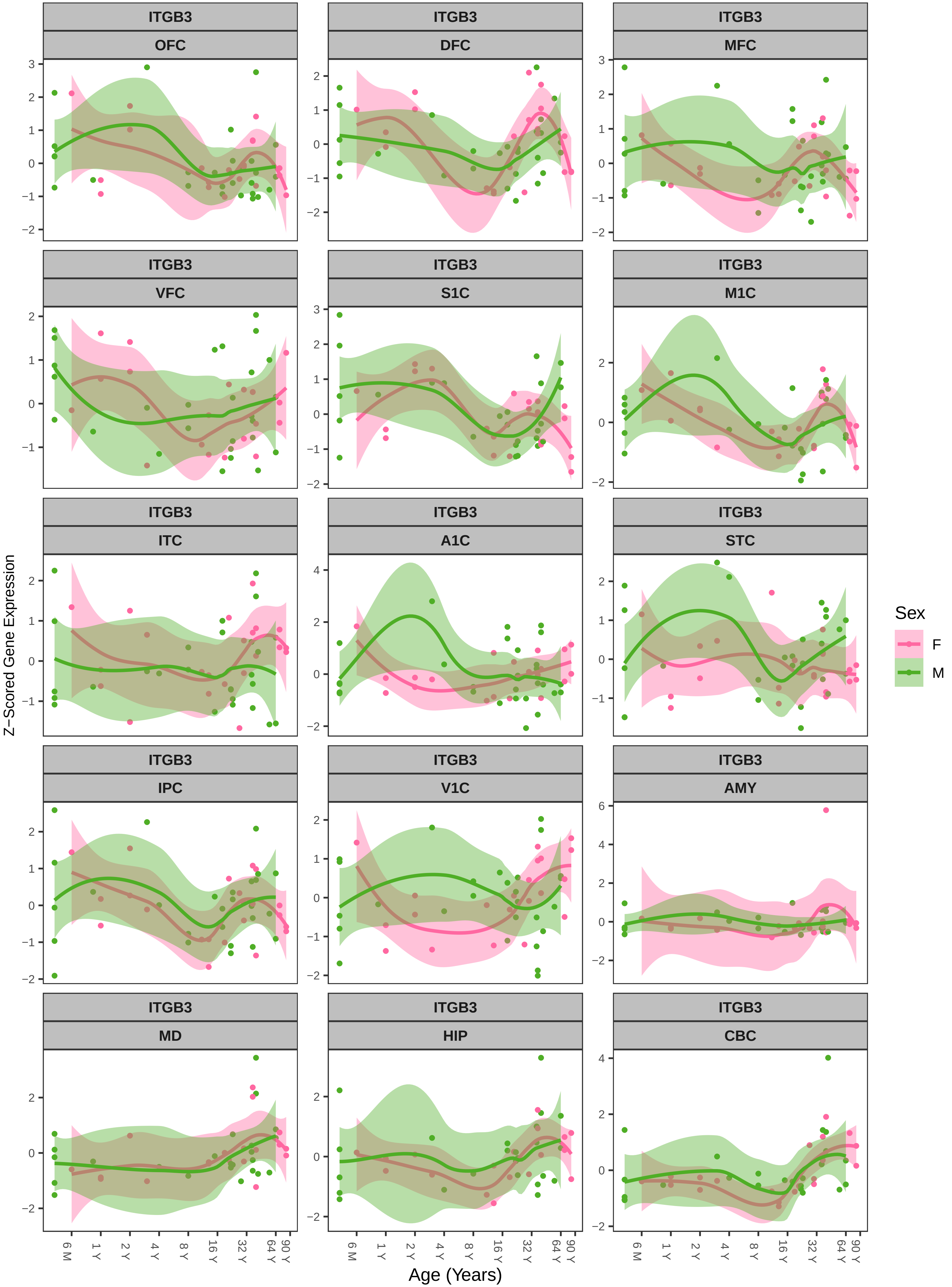
ITGB3
OFC
DFC
MFC
VFC
S1C
M1C
ITC
A1C
STC
IPC
V1C
AMY
MD
HIP
CBC
Z−Scored Gene Expression
Age (Years)
6 M
1 Y
2 Y
4 Y
8 Y
16 Y
32 Y
64 Y
90 Y
Sex
F
M

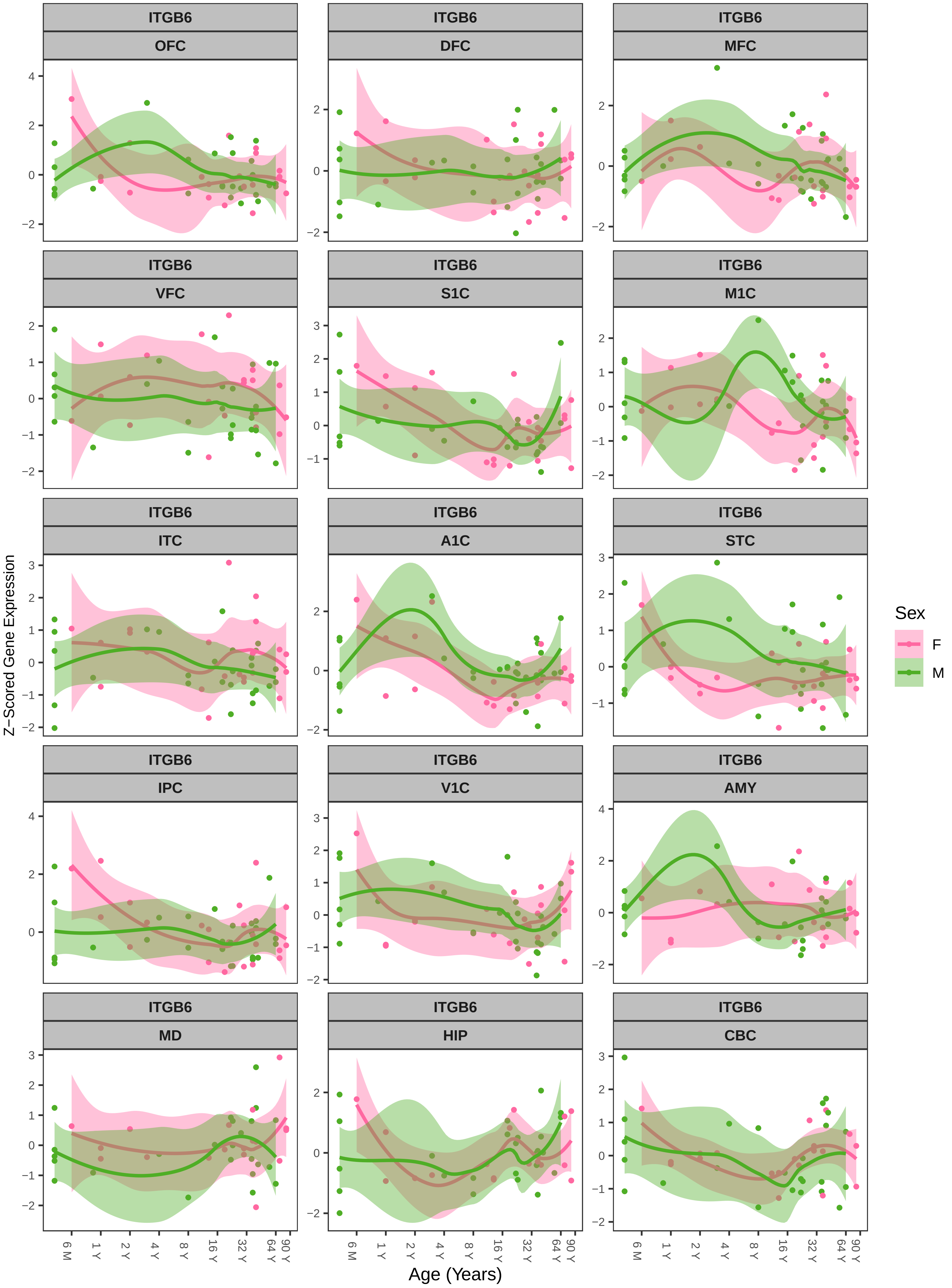
ITGB6
OFC
DFC
MFC
VFC
S1C
M1C
ITC
A1C
STC
IPC
V1C
AMY
MD
HIP
CBC
Z−Scored Gene Expression
Age (Years)
6 M
1 Y
2 Y
4 Y
8 Y
16 Y
32 Y
64 Y
90 Y
Sex
F
M

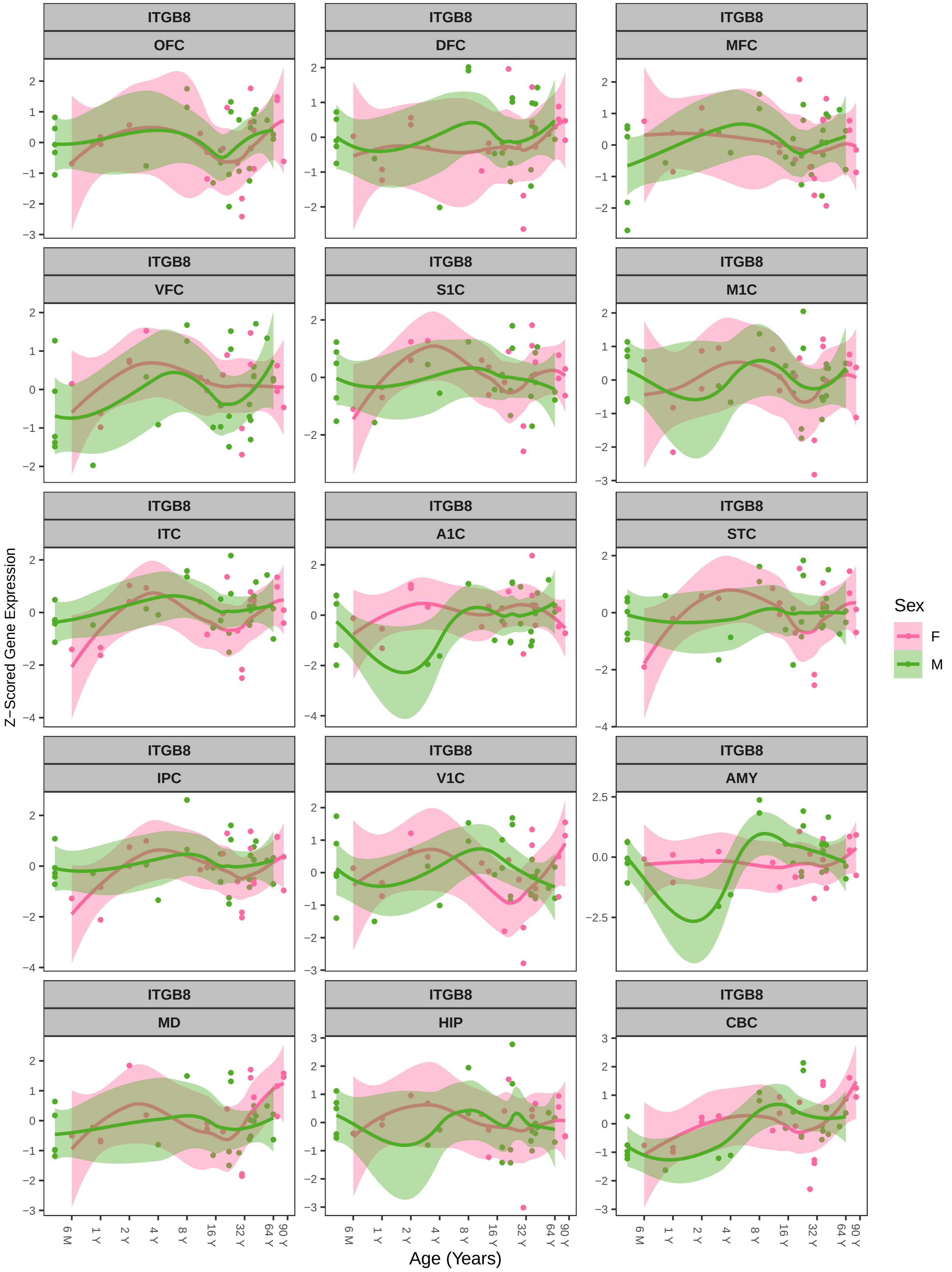
ITGB8
OFC
DFC
MFC
VFC
S1C
M1C
ITC
A1C
STC
IPC
V1C
AMY
MD
HIP
CBC
Z−Scored Gene Expression
Age (Years)
6 M
1 Y
2 Y
4 Y
8 Y
16 Y
32 Y
64 Y
90 Y
Sex
F
M

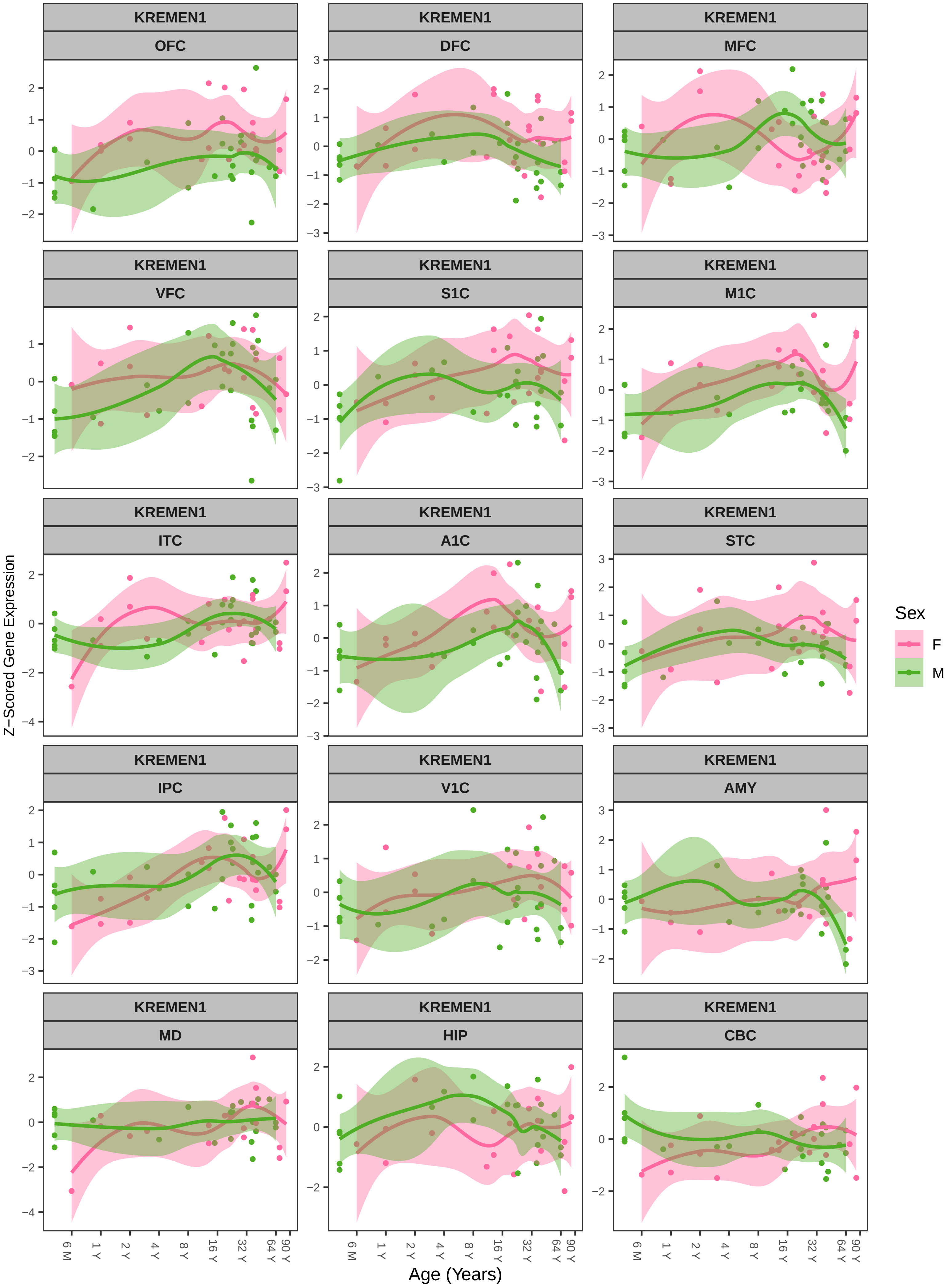
KREMEN1
OFC
DFC
MFC
VFC
S1C
M1C
ITC
A1C
STC
IPC
V1C
AMY
MD
HIP
CBC
Z−Scored Gene Expression
Age (Years)
6 M
1 Y
2 Y
4 Y
8 Y
16 Y
32 Y
64 Y
90 Y
Sex
F
M

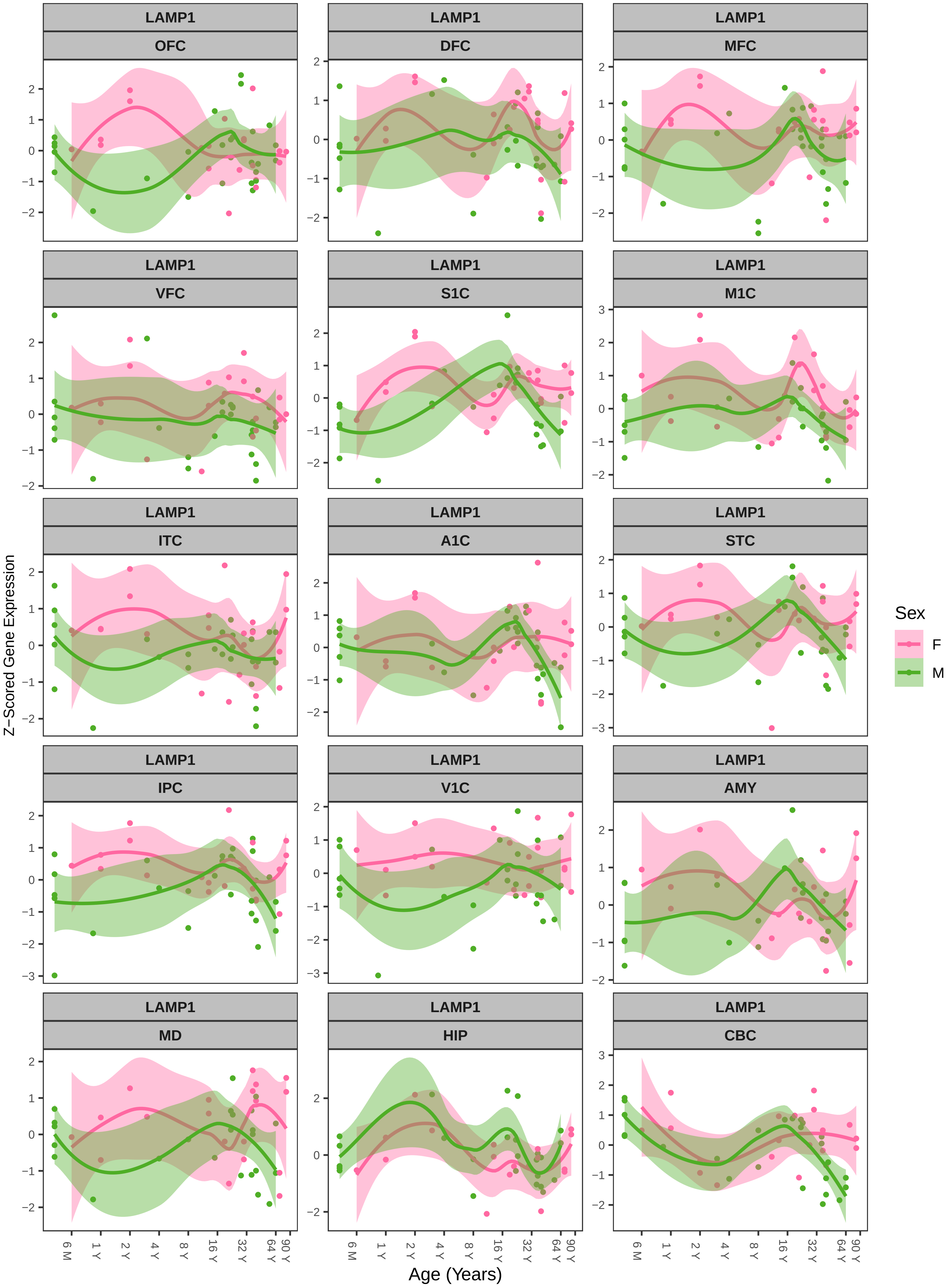

LAMP1
OFC
DFC
MFC
VFC
S1C
M1C
ITC
A1C
STC
IPC
V1C
AMY
MD
HIP
CBC
Z−Scored Gene Expression
Age (Years)
6 M
1 Y
2 Y
4 Y
8 Y
16 Y
32 Y
64 Y
90 Y
Sex
F
M

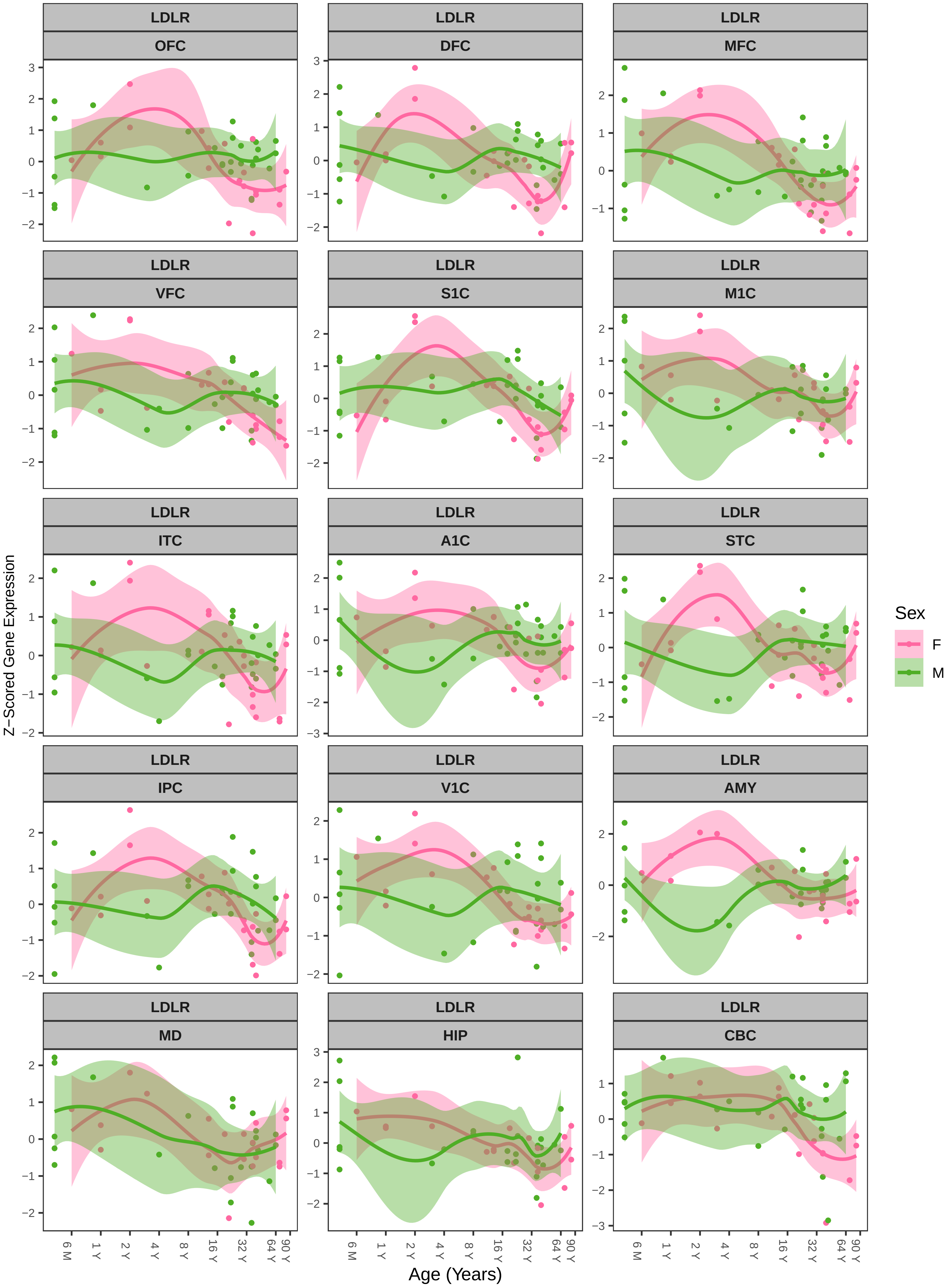
LDLR
OFC
DFC
MFC
VFC
S1C
M1C
ITC
A1C
STC
IPC
V1C
AMY
MD
HIP
CBC
Z−Scored Gene Expression
Age (Years)
6 M
1 Y
2 Y
4 Y
8 Y
16 Y
32 Y
64 Y
90 Y
Sex
F
M

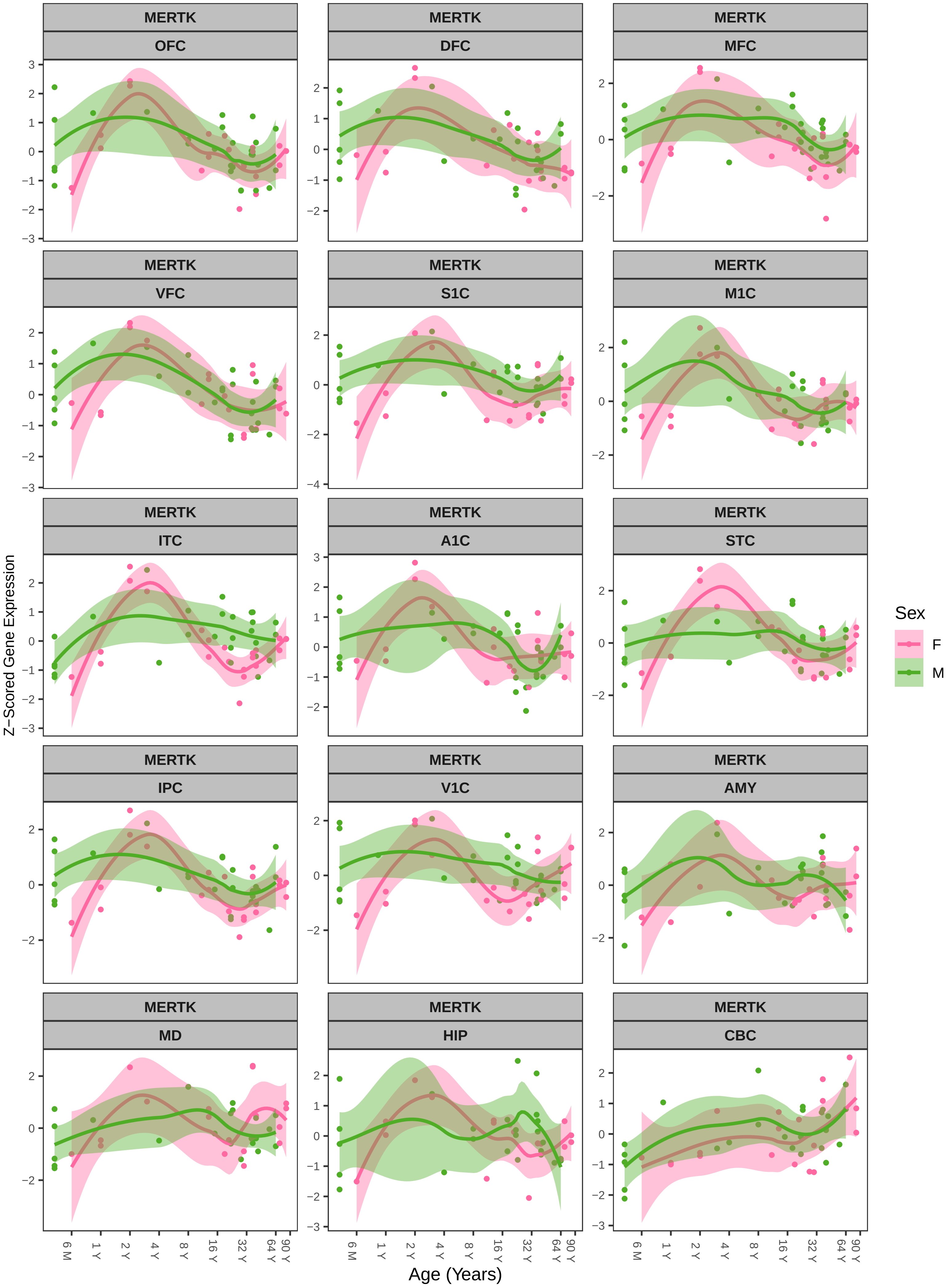
MERTK
OFC
DFC
MFC
VFC
S1C
M1C
ITC
A1C
STC
IPC
V1C
AMY
MD
HIP
CBC
Z−Scored Gene Expression
Age (Years)
6 M
1 Y
2 Y
4 Y
8 Y
16 Y
32 Y
64 Y
90 Y
Sex
F
M

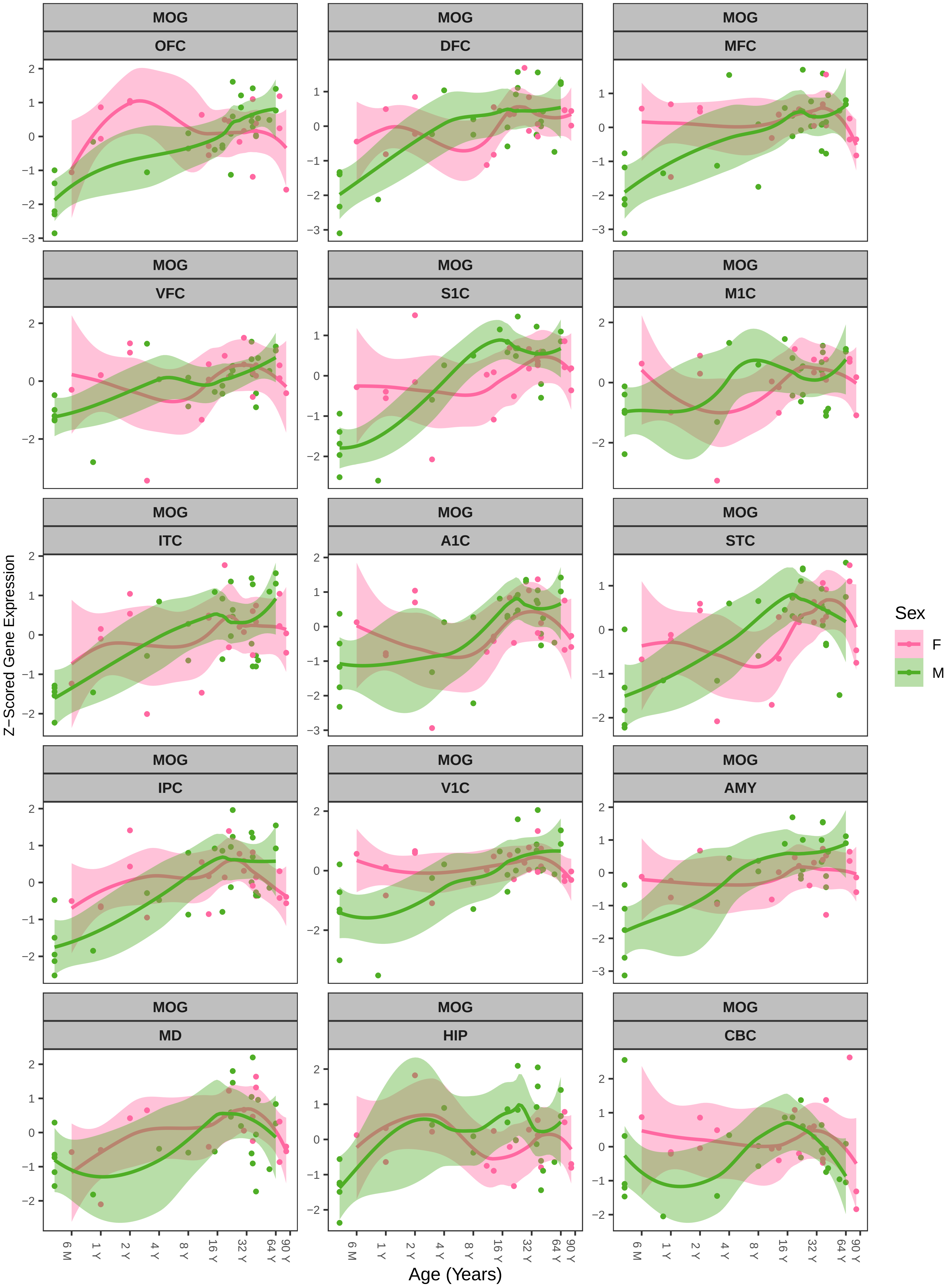
MOG
OFC
DFC
MFC
VFC
S1C
M1C
ITC
A1C
STC
IPC
V1C
AMY
MD
HIP
CBC
Z−Scored Gene Expression
Age (Years)
6 M
1 Y
2 Y
4 Y
8 Y
16 Y
32 Y
64 Y
90 Y
Sex
F
M

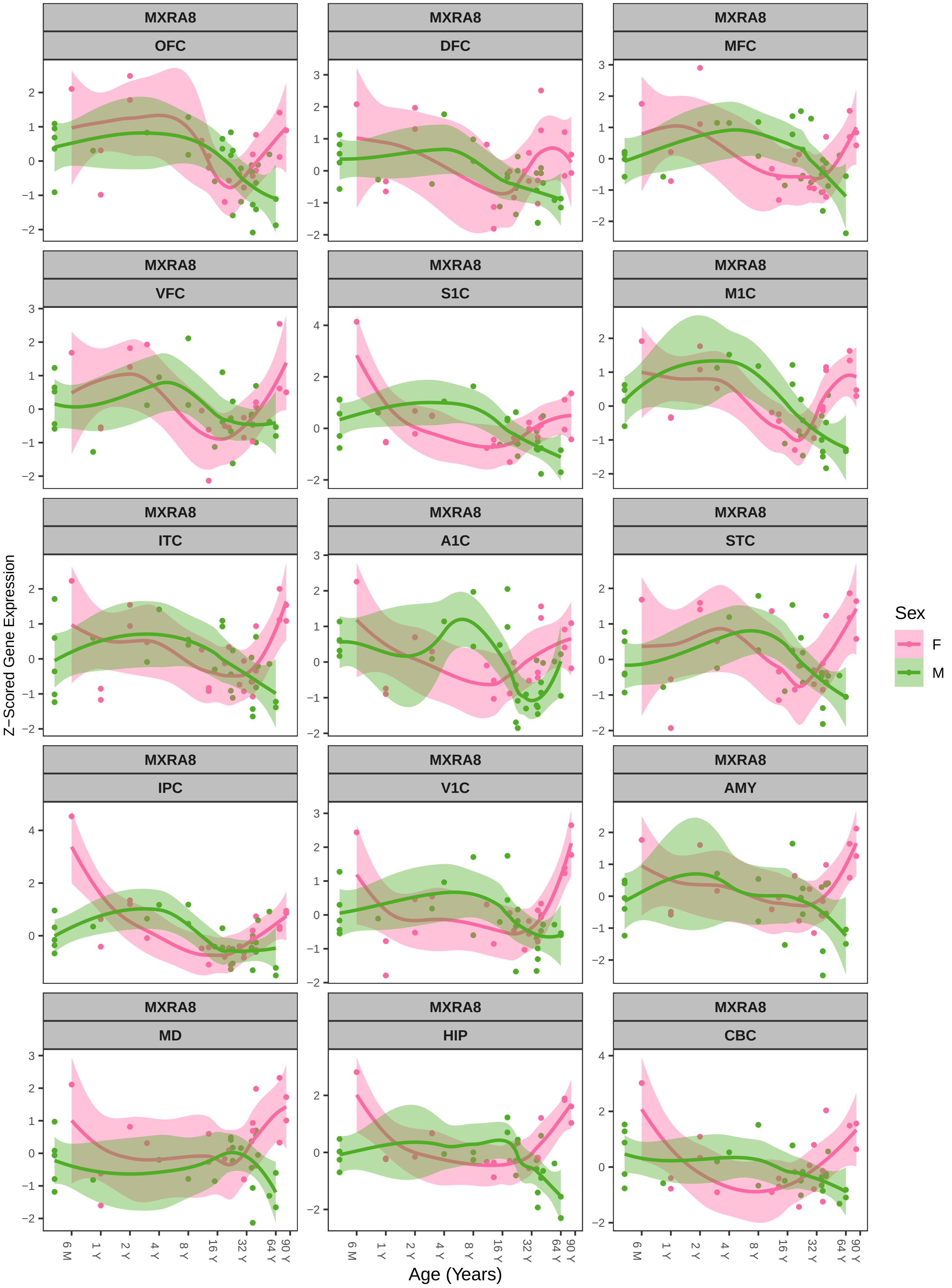
MXRA8
OFC
DFC
MFC
VFC
S1C
M1C
ITC
A1C
STC
IPC
V1C
AMY
MD
HIP
CBC
Z−Scored Gene Expression
Age (Years)
6 M
1 Y
2 Y
4 Y
8 Y
16 Y
32 Y
64 Y
90 Y
Sex
F
M

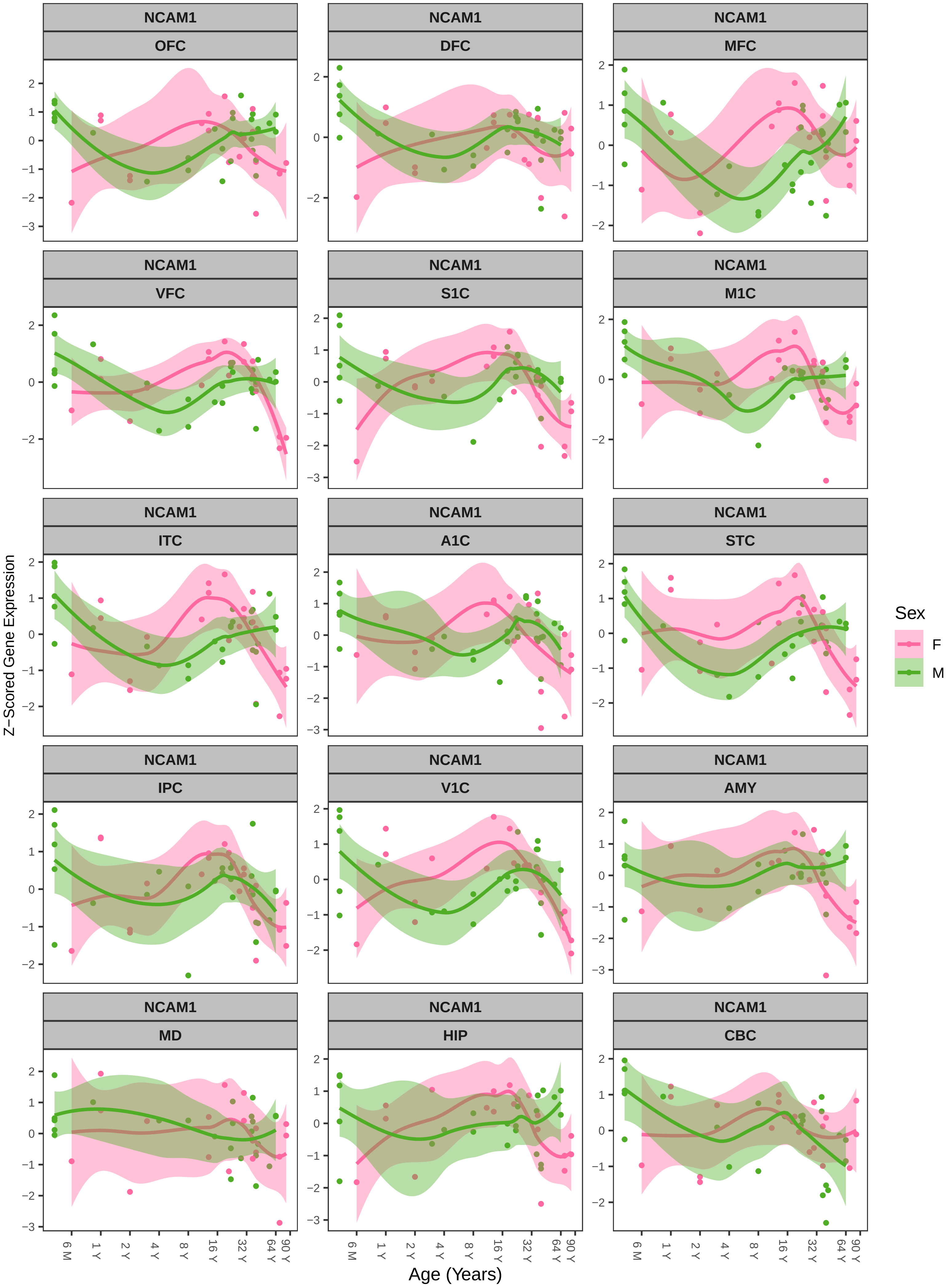
NCAM1
OFC
DFC
MFC
VFC
S1C
M1C
ITC
A1C
STC
IPC
V1C
AMY
MD
HIP
CBC
Z−Scored Gene Expression
Age (Years)
6 M
1 Y
2 Y
4 Y
8 Y
16 Y
32 Y
64 Y
90 Y
Sex
F
M

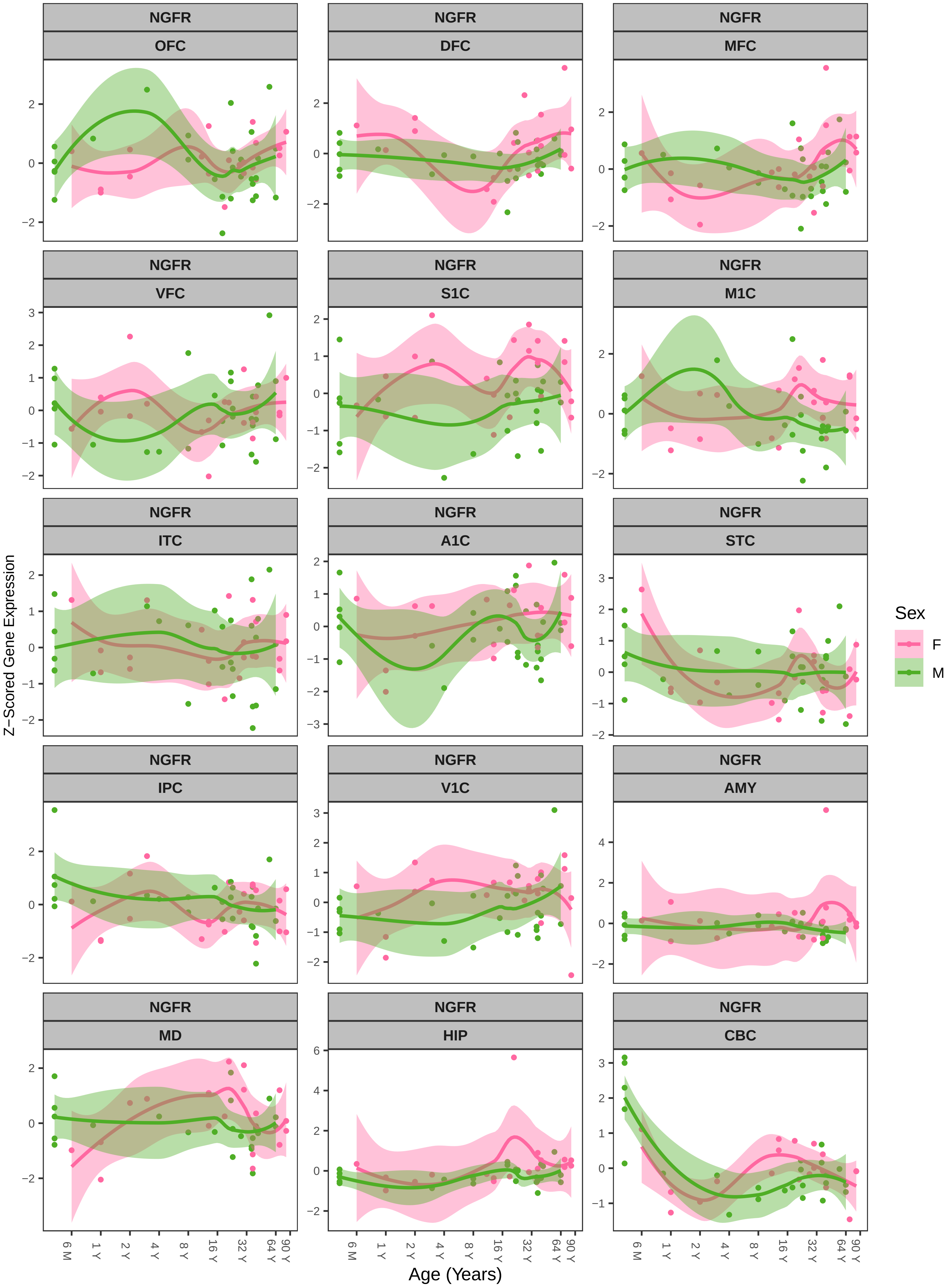

NGFR OFC
NGFR DFC
NGFR MFC
NGFR VFC
NGFR S1C
NGFR M1C
NGFR ITC
NGFR A1C
NGFR STC
NGFR IPC
NGFR V1C
NGFR AMY
NGFR MD
NGFR HIP
NGFR CBC
Z−Scored Gene Expression
Age (Years)
6 M 1 Y 2 Y 4 Y 8 Y 16 Y 32 Y 64 Y 90 Y
Sex
F
M

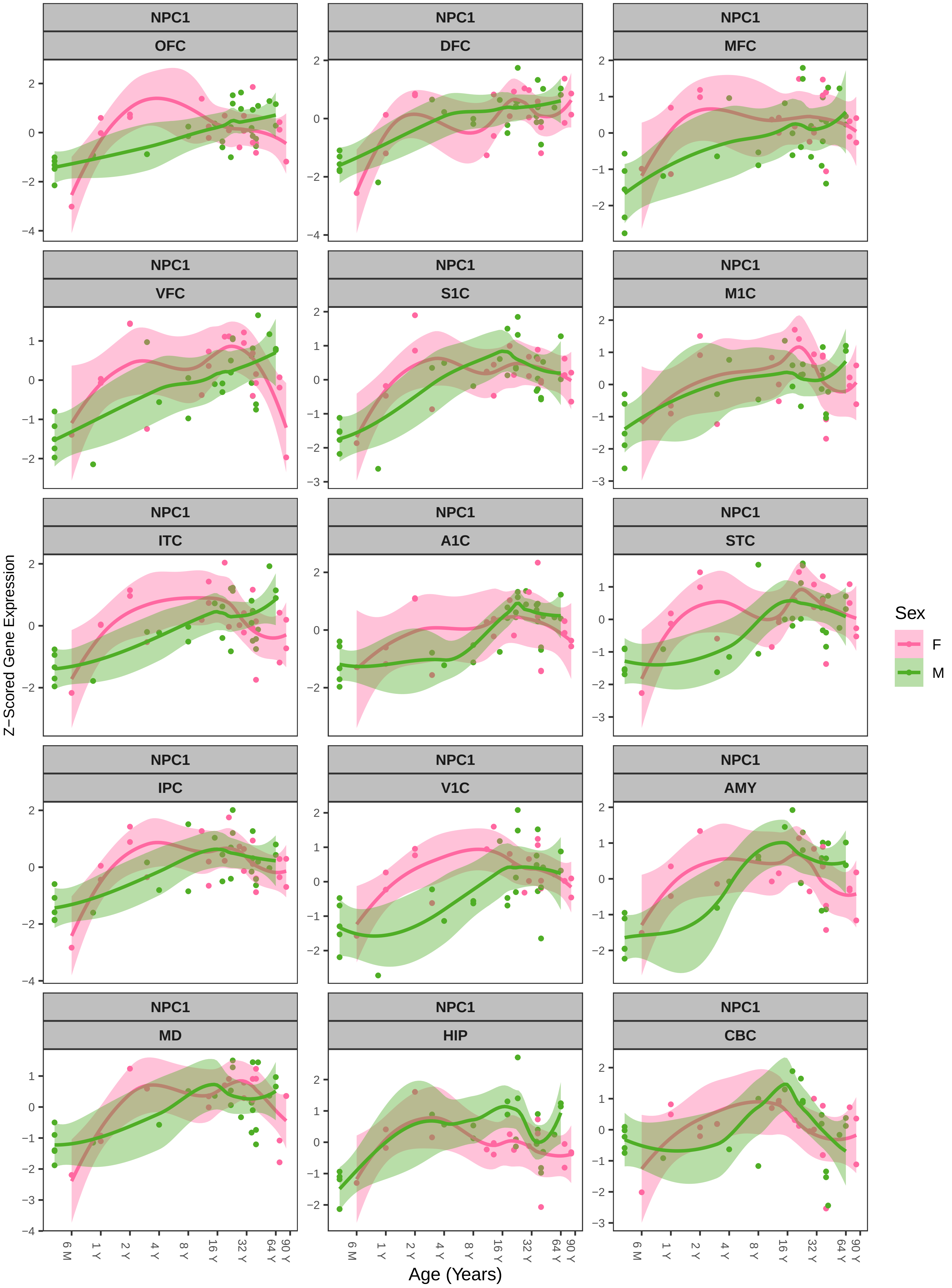

NPC1
OFC
DFC
MFC
VFC
S1C
M1C
ITC
A1C
STC
IPC
V1C
AMY
MD
HIP
CBC
Z−Scored Gene Expression
Age (Years)
6 M
1 Y
2 Y
4 Y
8 Y
16 Y
32 Y
64 Y
90 Y
Sex
F
M

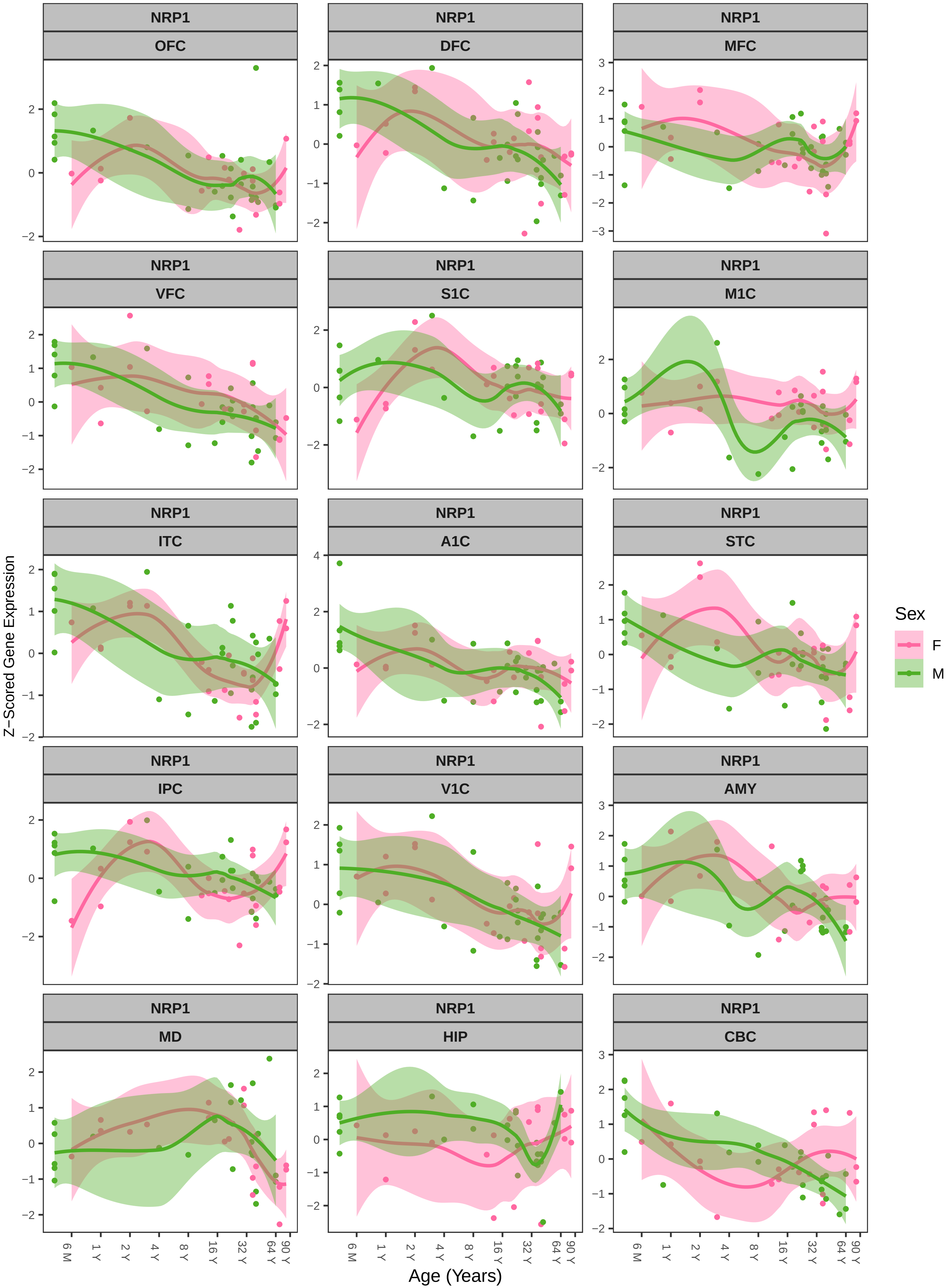

NRP1
OFC
DFC
MFC
VFC
S1C
M1C
ITC
A1C
STC
IPC
V1C
AMY
MD
HIP
CBC
Z−Scored Gene Expression
Age (Years)
6 M
1 Y
2 Y
4 Y
8 Y
16 Y
32 Y
64 Y
90 Y
Sex
F
M

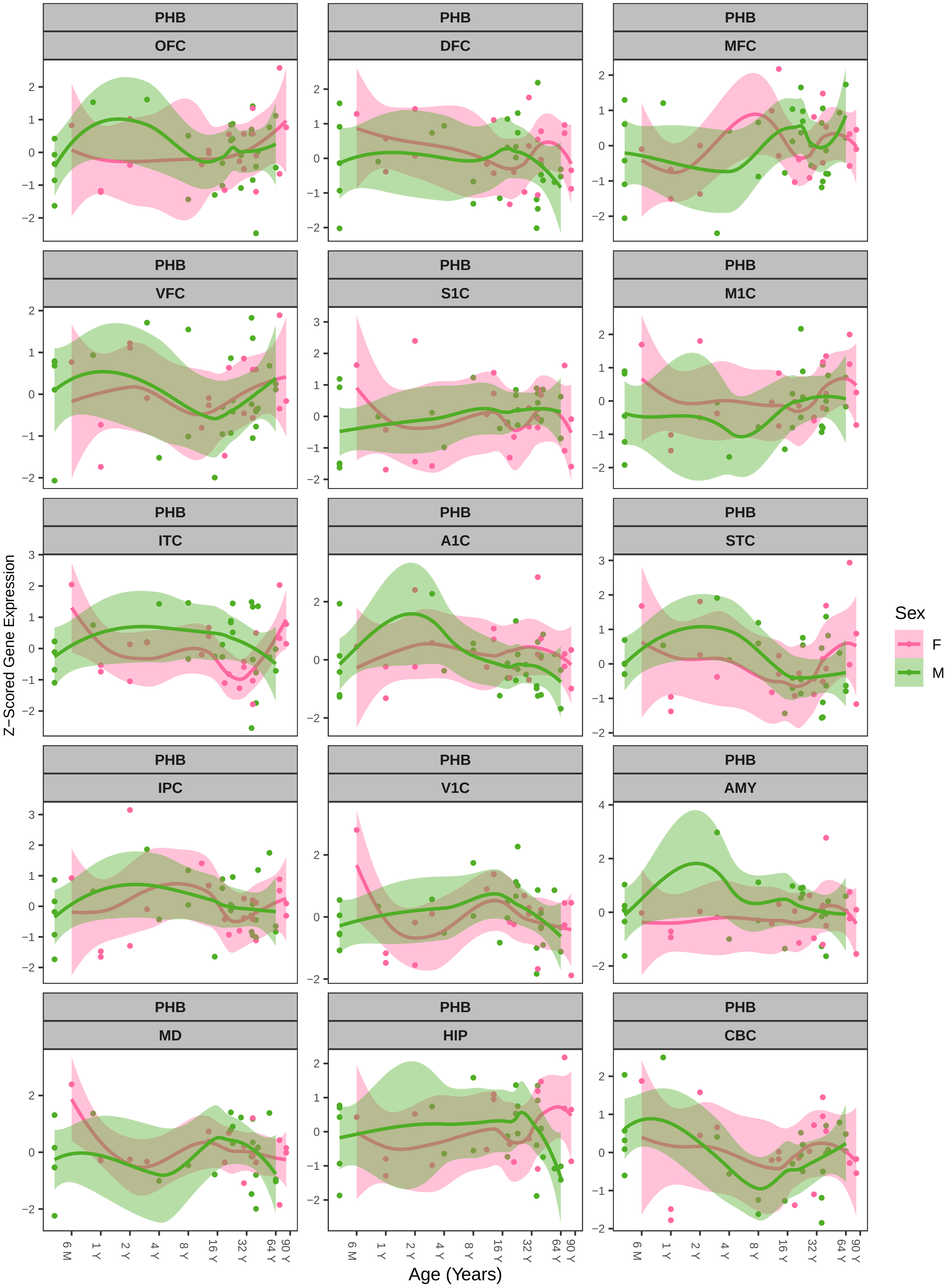

PHB
OFC
PHB
DFC
PHB
MFC
PHB
VFC
PHB
S1C
PHB
M1C
PHB
ITC
PHB
A1C
PHB
STC
PHB
IPC
PHB
V1C
PHB
AMY
PHB
MD
PHB
HIP
PHB
CBC
Z−Scored Gene Expression
Age (Years)
6 M
1 Y
2 Y
4 Y
8 Y
16 Y
32 Y
64 Y
90 Y
Sex
F
M

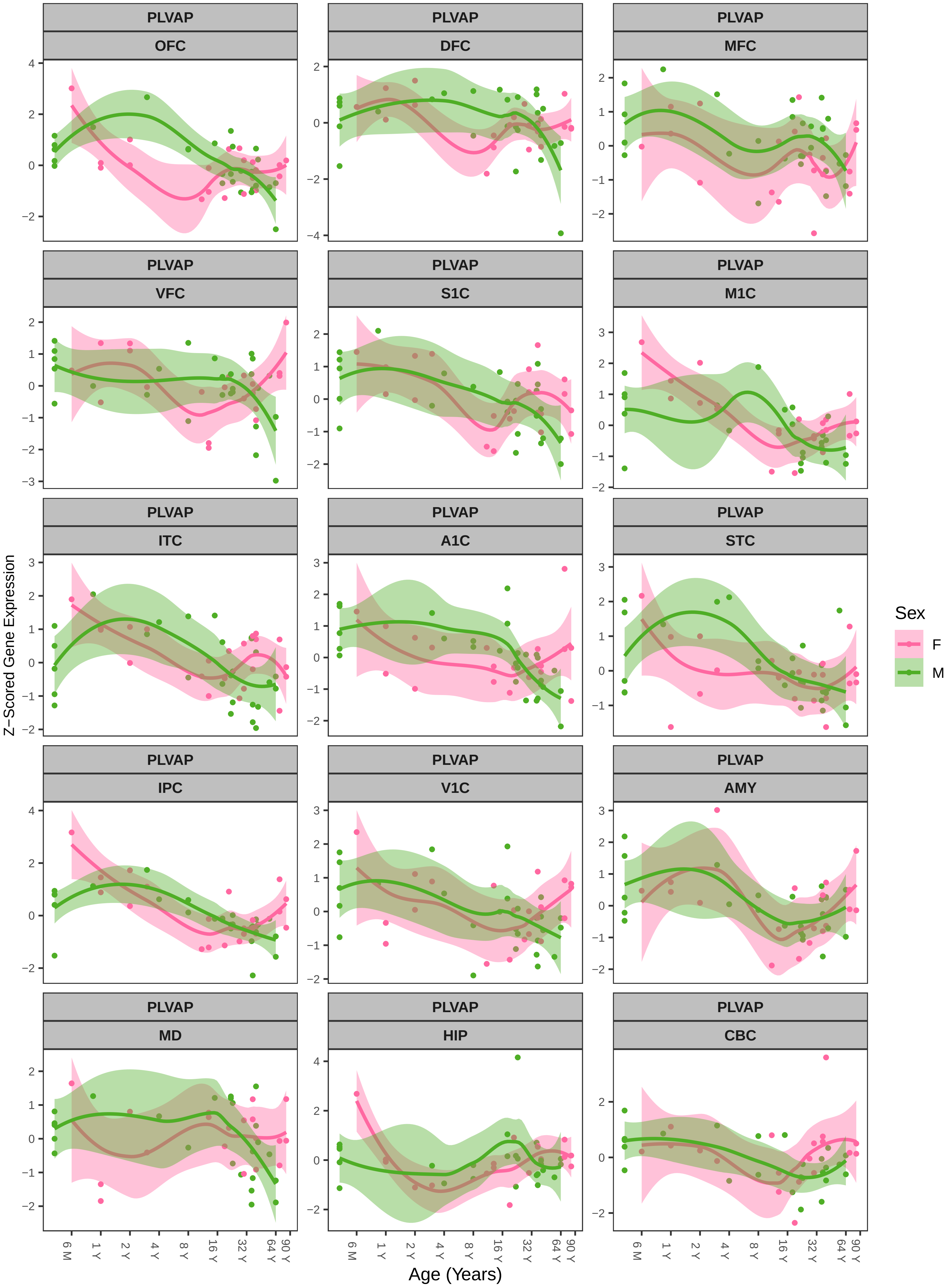

PLVAP
OFC
DFC
MFC
VFC
S1C
M1C
ITC
A1C
STC
IPC
V1C
AMY
MD
HIP
CBC
Z−Scored Gene Expression
Age (Years)
6 M
1 Y
2 Y
4 Y
8 Y
16 Y
32 Y
64 Y
90 Y
Sex
F
M

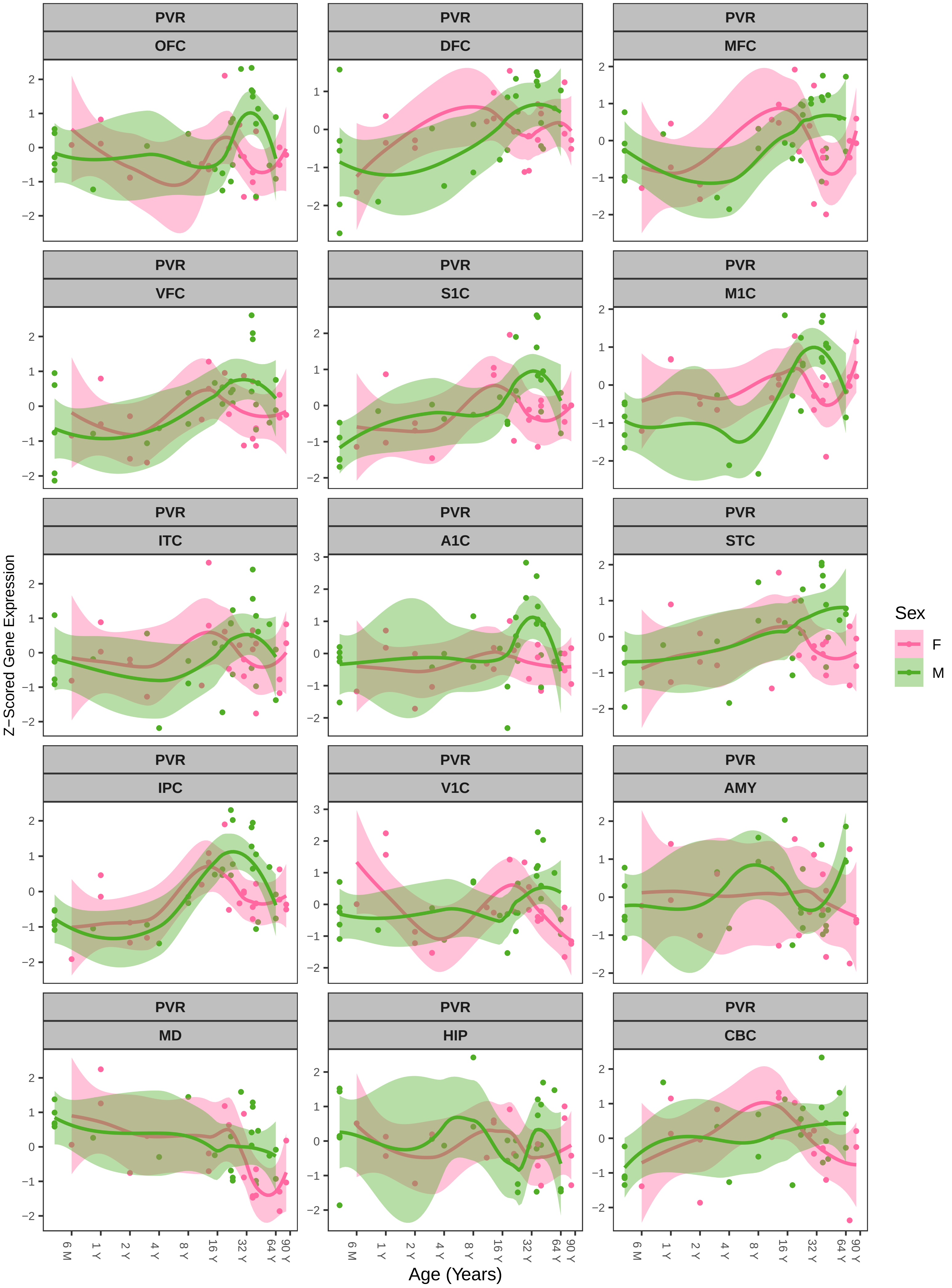
PVR
OFC
DFC
MFC
VFC
S1C
M1C
ITC
A1C
STC
IPC
V1C
AMY
MD
HIP
CBC
Z−Scored Gene Expression
Age (Years)
6 M
1 Y
2 Y
4 Y
8 Y
16 Y
32 Y
64 Y
90 Y
Sex
F
M

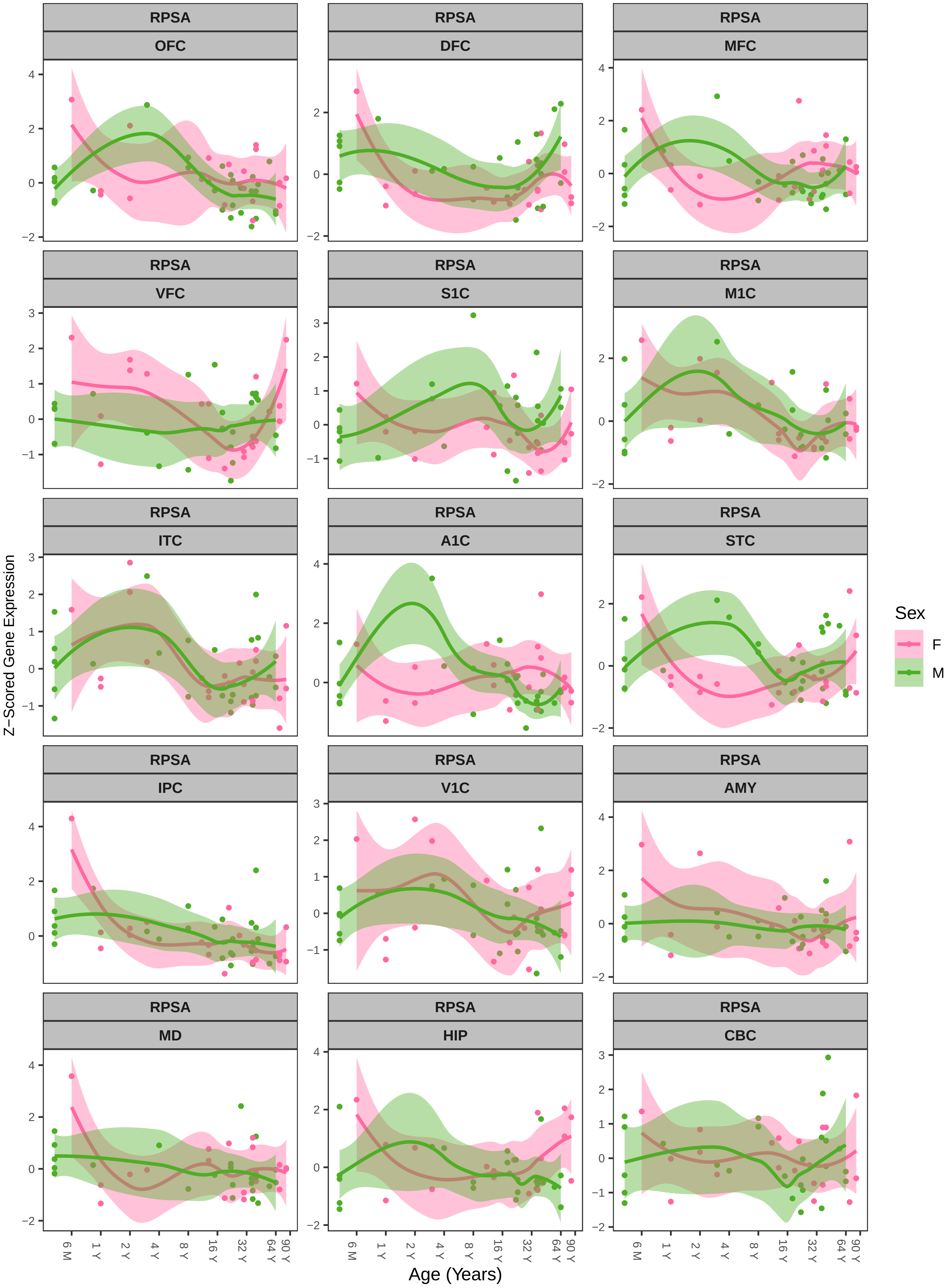

RPSA
OFC
DFC
MFC
VFC
S1C
M1C
ITC
A1C
STC
IPC
V1C
AMY
MD
HIP
CBC
Z−Scored Gene Expression
Age (Years)
6 M
1 Y
2 Y
4 Y
8 Y
16 Y
32 Y
64 Y
90 Y
Sex
F
M

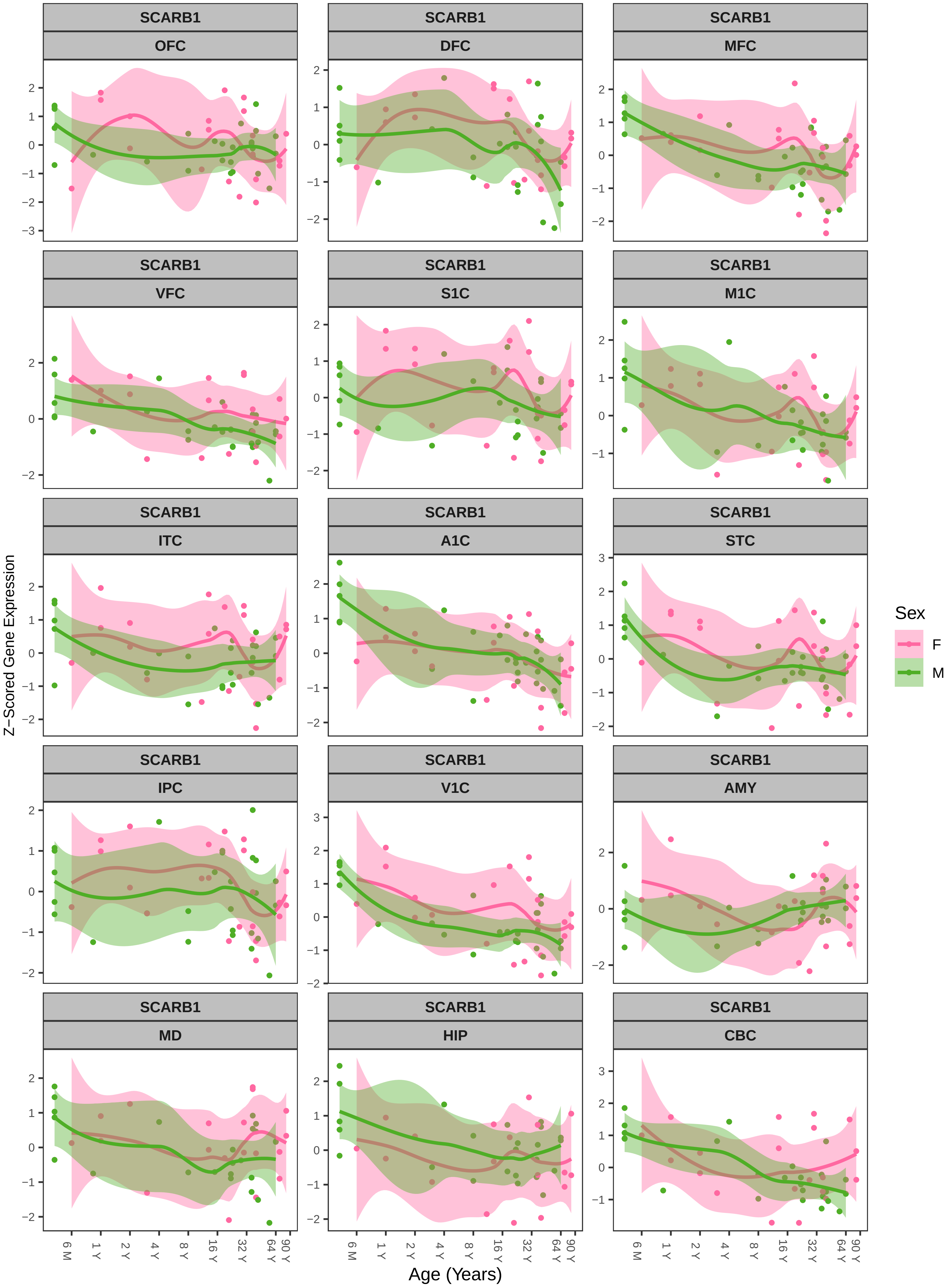
SCARB1
OFC
DFC
MFC
VFC
S1C
M1C
ITC
A1C
STC
IPC
V1C
AMY
MD
HIP
CBC
Z−Scored Gene Expression
Age (Years)
6 M
1 Y
2 Y
4 Y
8 Y
16 Y
32 Y
64 Y
90 Y
Sex
F
M

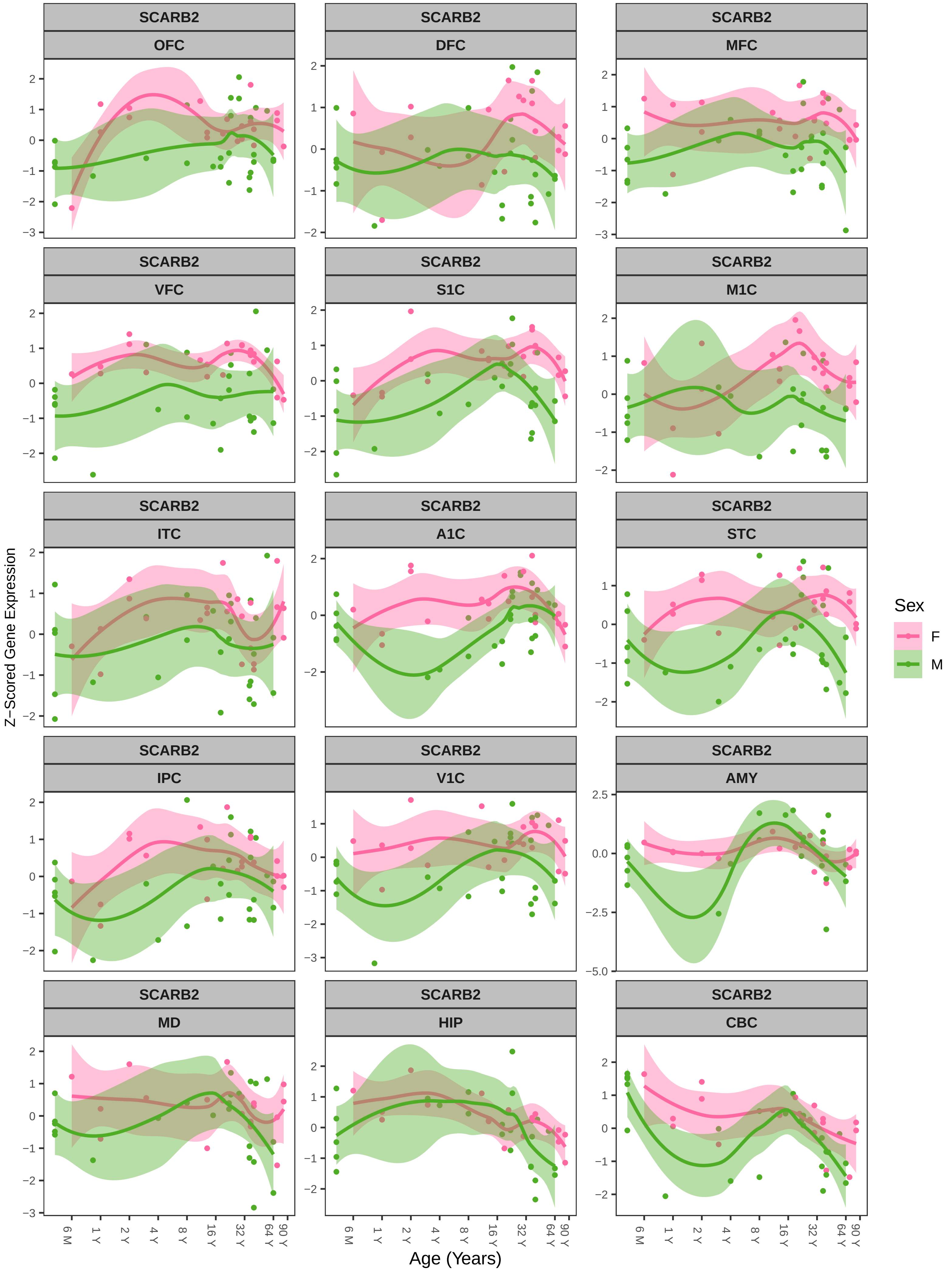

SCARB2
OFC
DFC
MFC
VFC
S1C
M1C
ITC
A1C
STC
IPC
V1C
AMY
MD
HIP
CBC
Z−Scored Gene Expression
Age (Years)
6 M
1 Y
2 Y
4 Y
8 Y
16 Y
32 Y
64 Y
90 Y
Sex
F
M

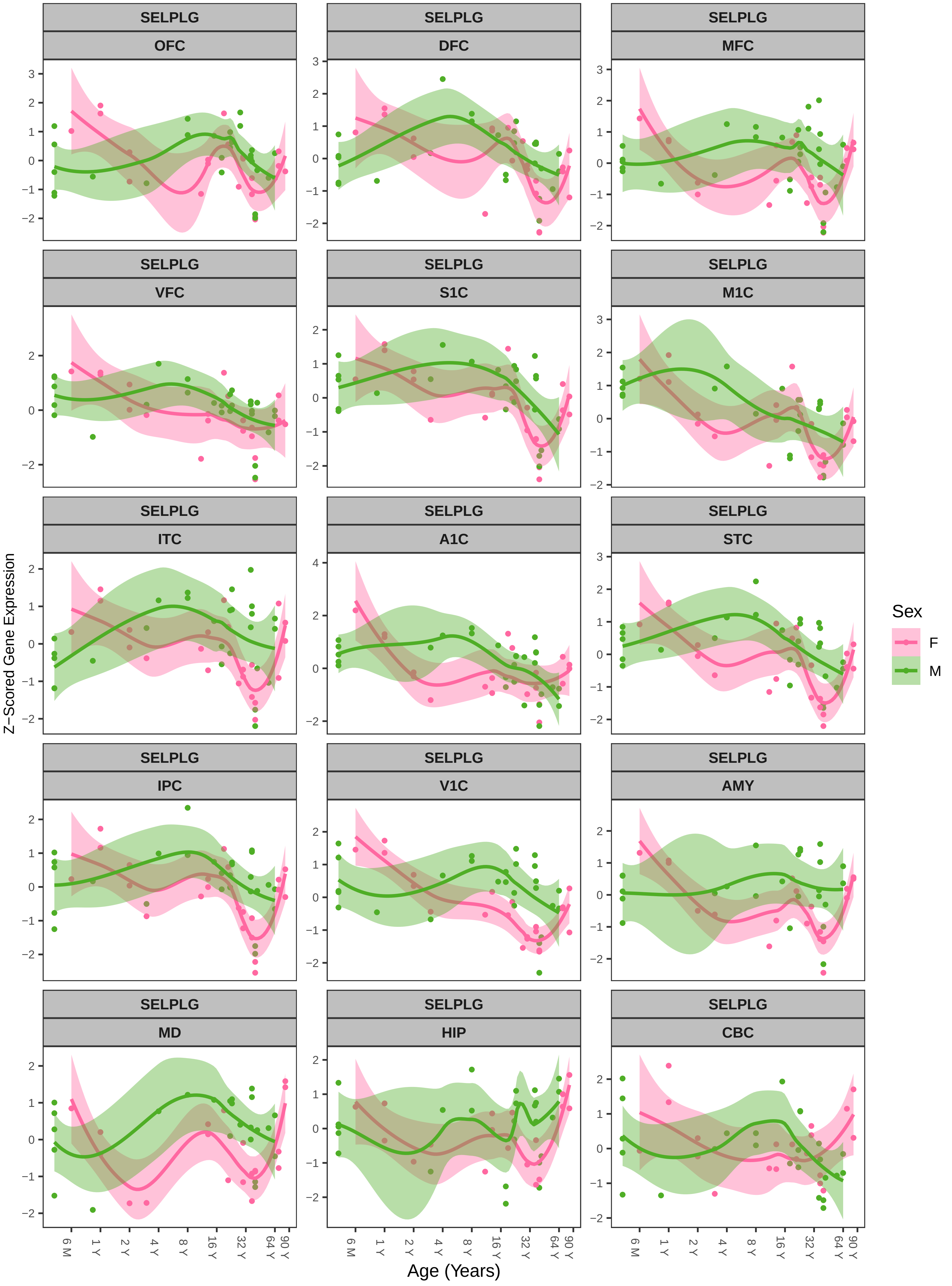
SELPLG
OFC
DFC
MFC
VFC
S1C
M1C
ITC
A1C
STC
IPC
V1C
AMY
MD
HIP
CBC
Z−Scored Gene Expression
Age (Years)
6 M
1 Y
2 Y
4 Y
8 Y
16 Y
32 Y
64 Y
90 Y
Sex
F
M

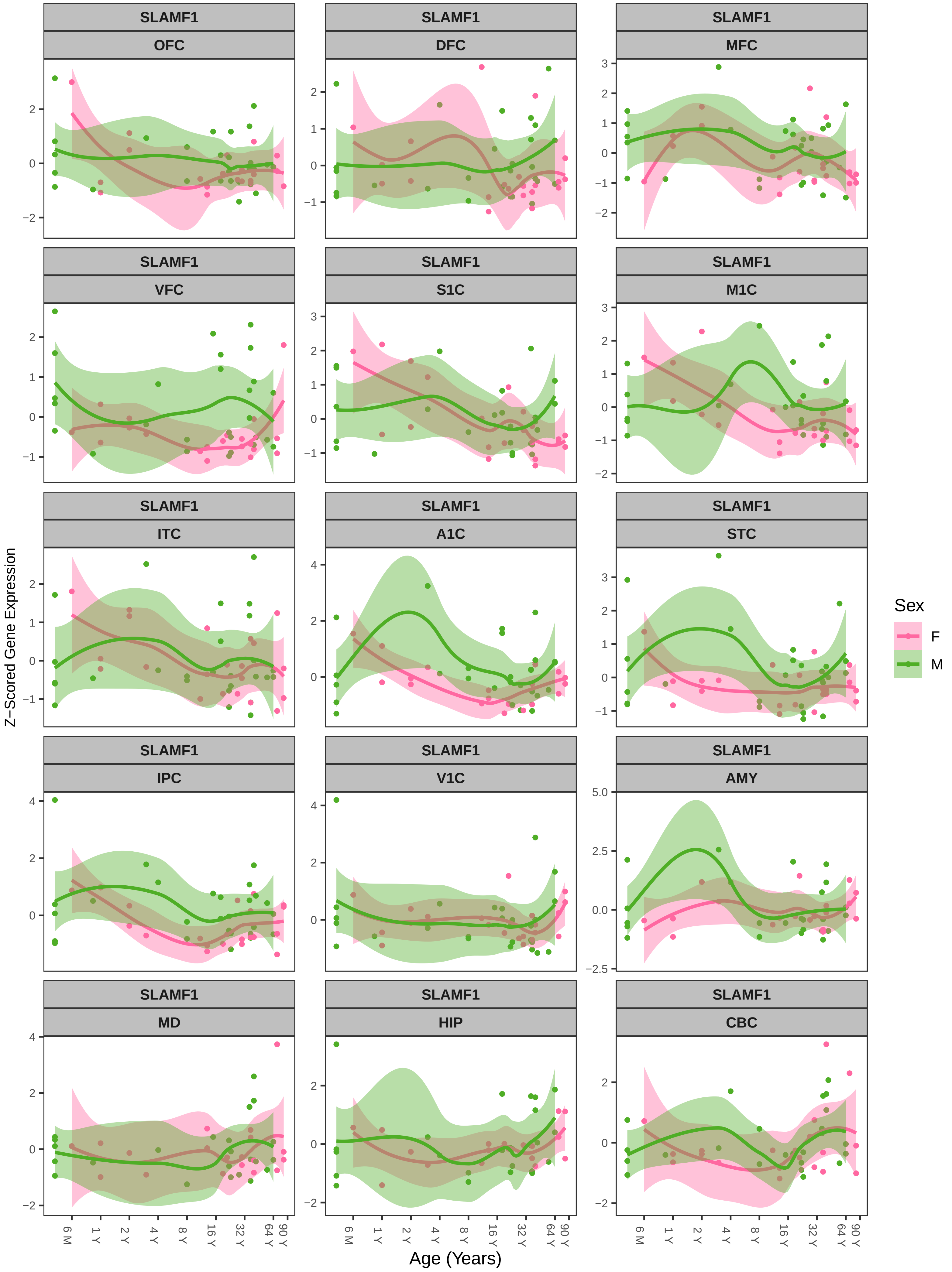
SLAMF1
OFC
DFC
MFC
VFC
S1C
M1C
ITC
A1C
STC
IPC
V1C
AMY
MD
HIP
CBC
Z−Scored Gene Expression
Age (Years)
6 M
1 Y
2 Y
4 Y
8 Y
16 Y
32 Y
64 Y
90 Y
Sex
F
M

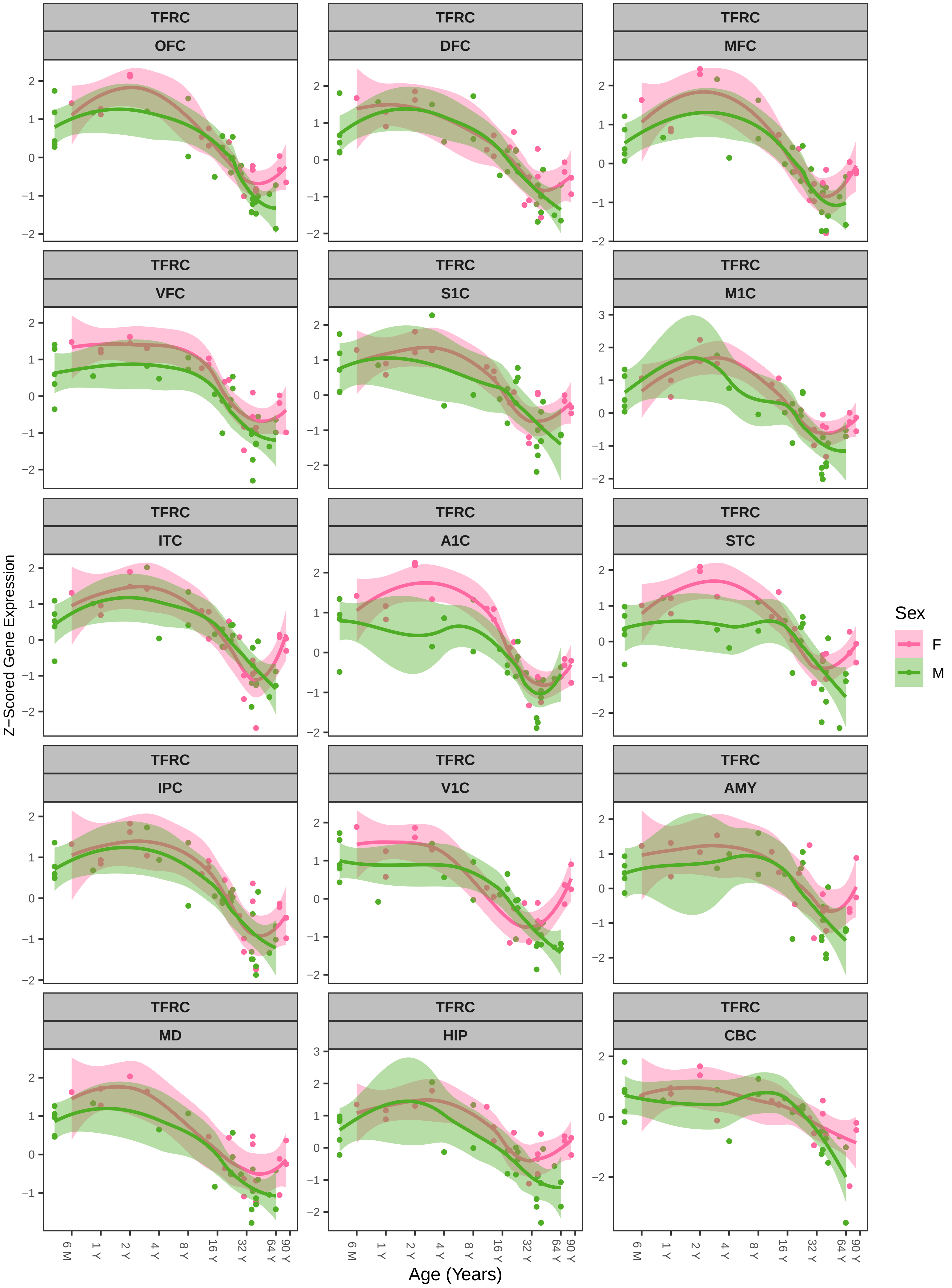
TFRC
OFC
DFC
MFC
VFC
S1C
M1C
ITC
A1C
STC
IPC
V1C
AMY
MD
HIP
CBC
Z−Scored Gene Expression
Age (Years)
6 M
1 Y
2 Y
4 Y
8 Y
16 Y
32 Y
64 Y
90 Y
Sex
F
M

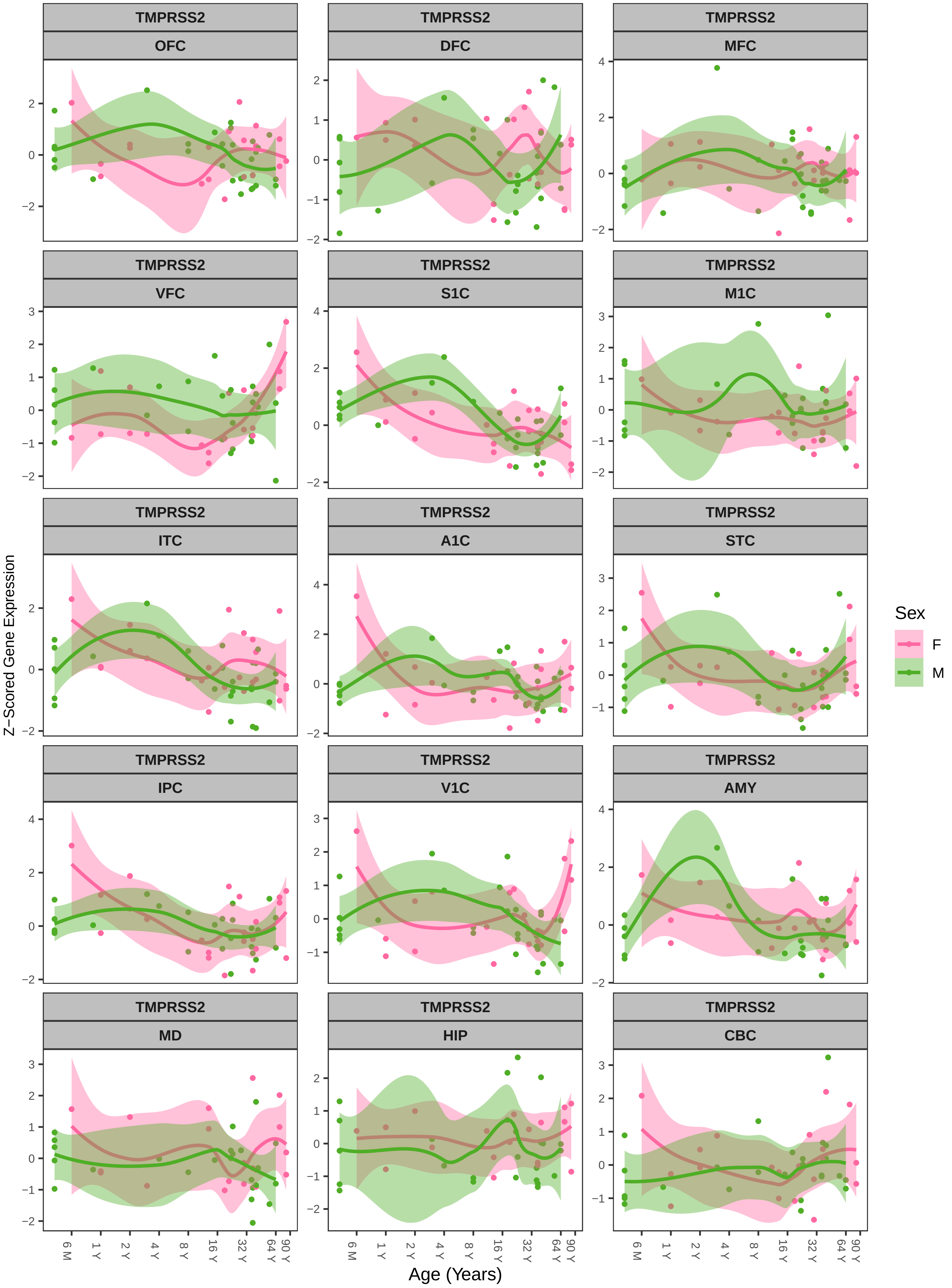

TMPRSS2
OFC
DFC
MFC
VFC
S1C
M1C
ITC
A1C
STC
IPC
V1C
AMY
MD
HIP
CBC
Z−Scored Gene Expression
Age (Years)
6 M
1 Y
2 Y
4 Y
8 Y
16 Y
32 Y
64 Y
90 Y
Sex
F
M

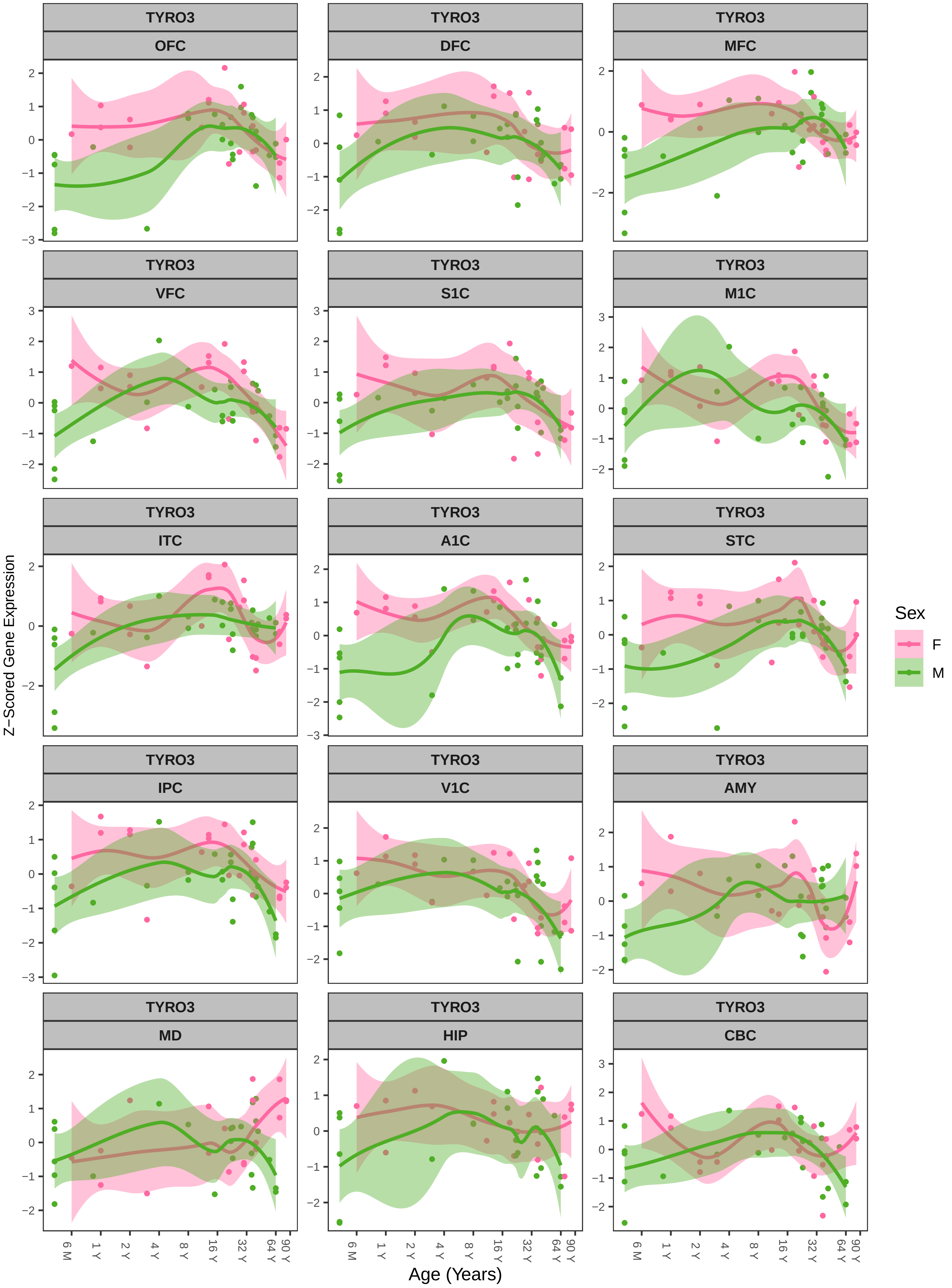

TYRO3
OFC
TYRO3
DFC
TYRO3
MFC
TYRO3
VFC
TYRO3
S1C
TYRO3
M1C
TYRO3
ITC
TYRO3
A1C
TYRO3
STC
TYRO3
IPC
TYRO3
V1C
TYRO3
AMY
TYRO3
MD
TYRO3
HIP
TYRO3
CBC
Z−Scored Gene Expression
Age (Years)
6 M
1 Y
2 Y
4 Y
8 Y
16 Y
32 Y
64 Y
90 Y
Sex
F
M

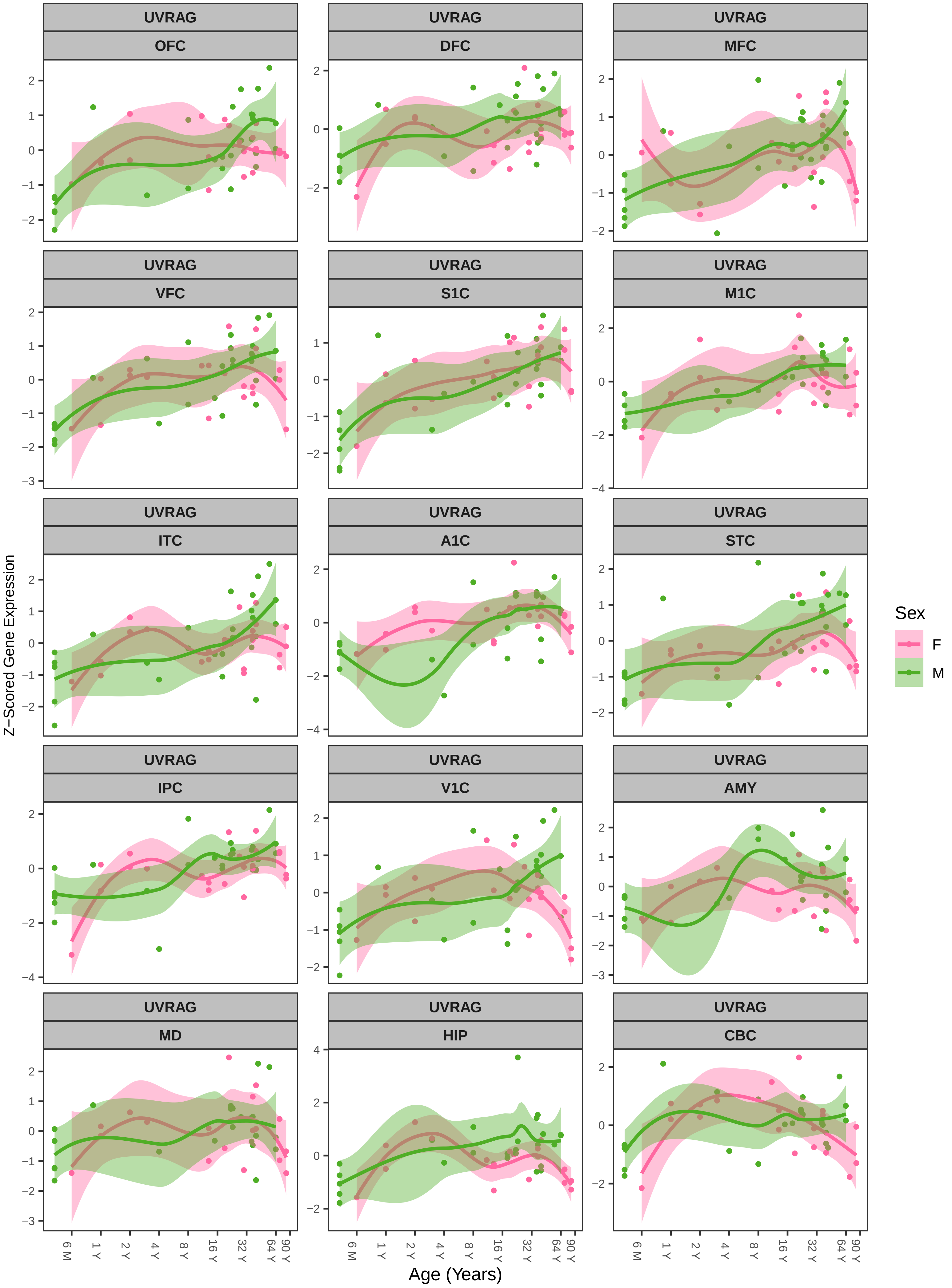
UVRAG OFC
UVRAG DFC
UVRAG MFC
UVRAG VFC
UVRAG S1C
UVRAG M1C
UVRAG ITC
UVRAG A1C
UVRAG STC
UVRAG IPC
UVRAG V1C
UVRAG AMY
UVRAG MD
UVRAG HIP
UVRAG CBC
Z−Scored Gene Expression
Age (Years)
6 M
1 Y
2 Y
4 Y
8 Y
16 Y
32 Y
64 Y
90 Y
Sex
F
M

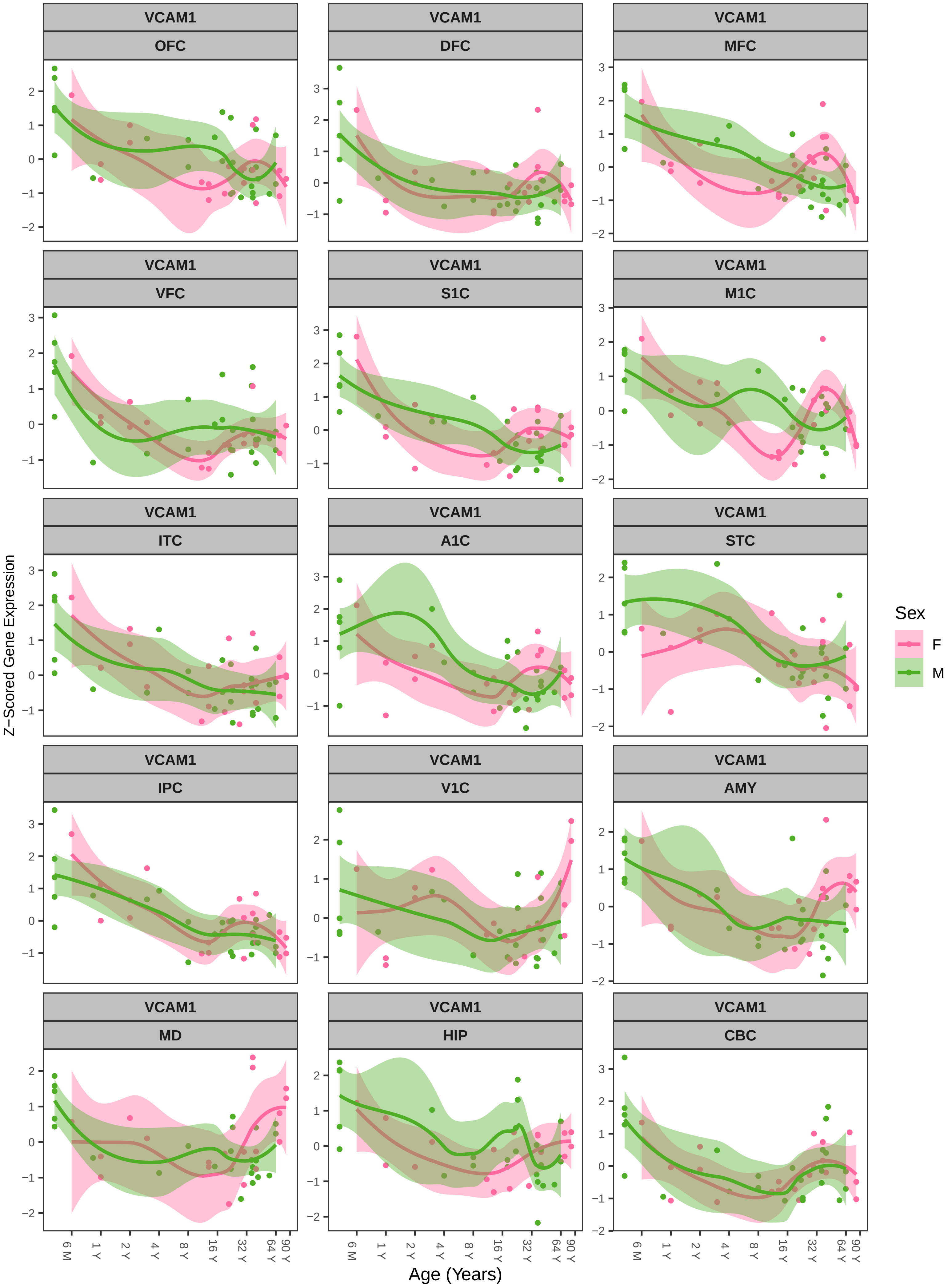

VCAM1
OFC
DFC
MFC
VFC
S1C
M1C
ITC
A1C
STC
IPC
V1C
AMY
MD
HIP
CBC
Z−Scored Gene Expression
Age (Years)
6 M
1 Y
2 Y
4 Y
8 Y
16 Y
32 Y
64 Y
90 Y
Sex
F
M
