## Supplementary Figures for "Sex- and age-specific development of ssRNA virus receptor expression in the human brain": Suppl Figs 2-9.pdf

**Supplementary Material**

**Supplementary Figures 2-9**

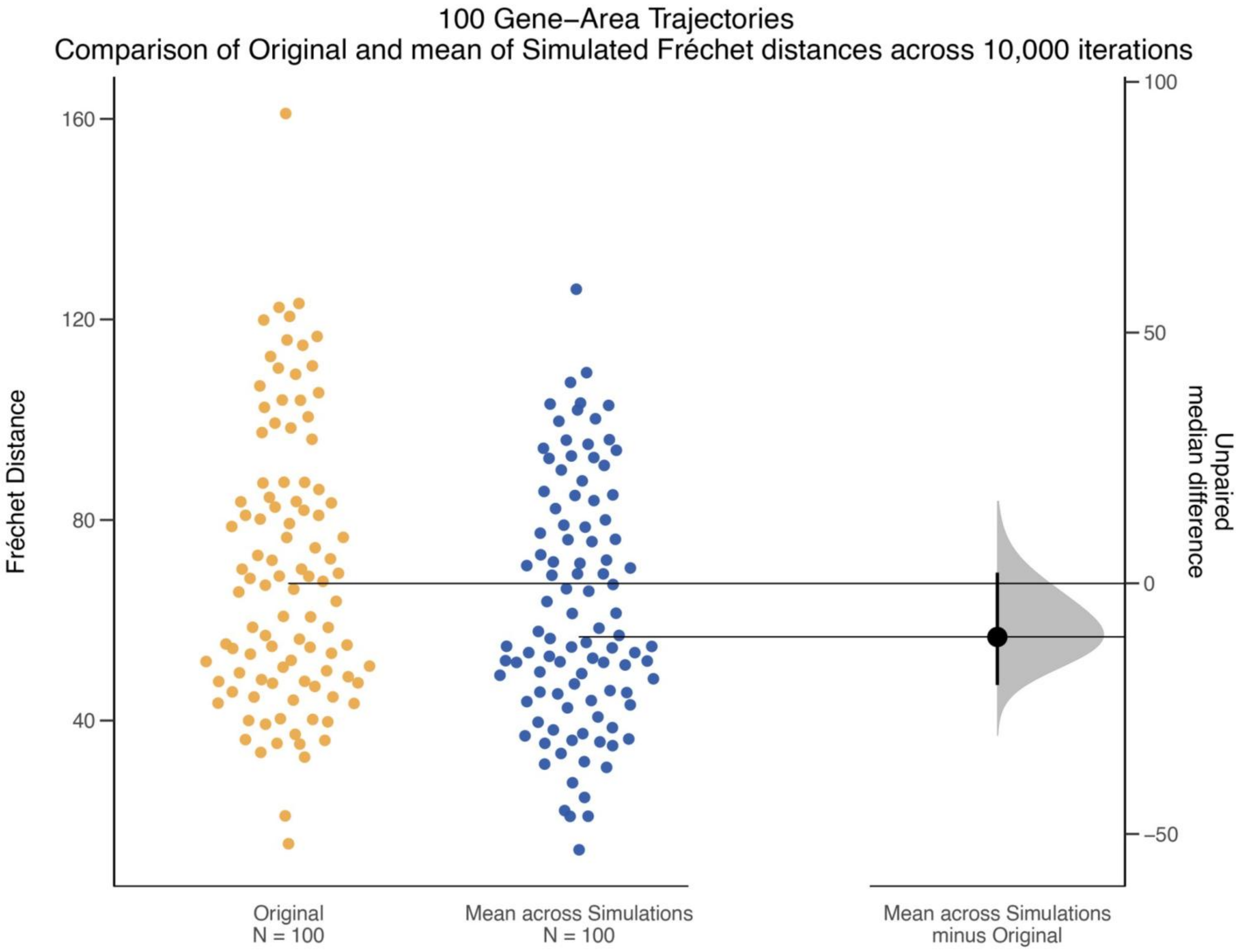


**Supplementary Figure 2** *Leave-one-case-out resampling to assess trajectory robustness and the stability of the sex-differences.* For a random subset of 100 gene-area combinations, female and male LOESS trajectories were repeatedly refit across 10,000 leave one case-out iterations, each time omitting one female and one male case. In each iteration, the Frchet distance between refit female and male trajectories was recalculated, generating a distribution of simulated distances for each combination. Estimation statistics were used to compare the original Frchet distance with the mean simulated value, quantified as the unpaired median difference shown as a black dot with its 95% confidence interval; grey shading represents the bootstrap sampling distributions.

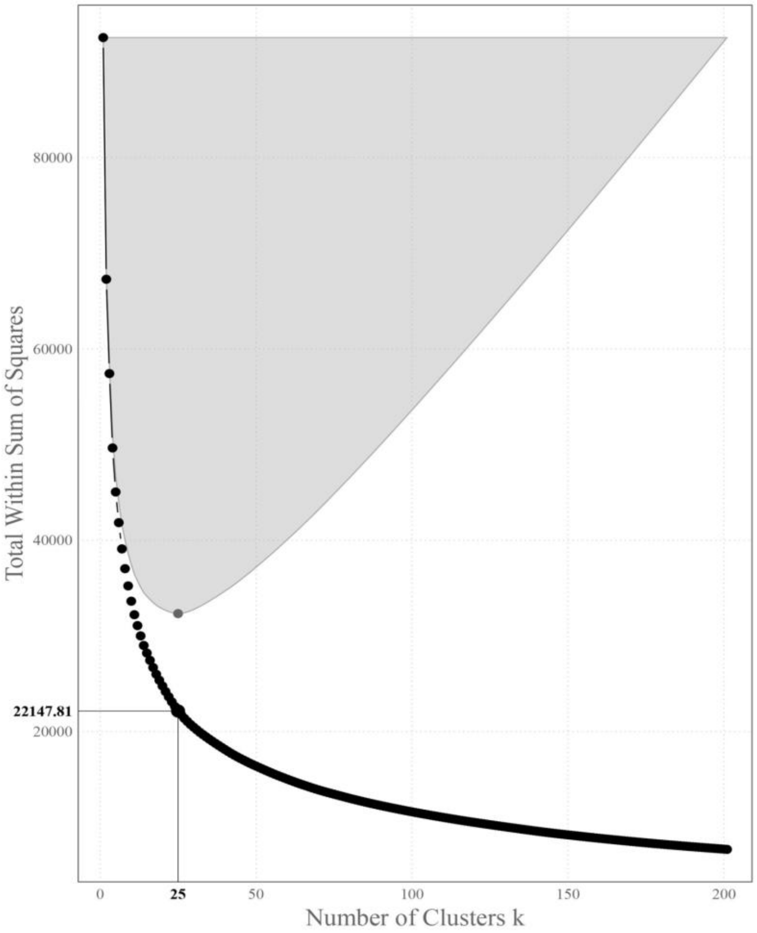


**Supplementary Figure 3** *Determining number of clusters.* Elbow plot showing the within cluster sum of squares plotted against number of clusters (*k*). The optimal k-value (k = 25) is indicated as the inflection point in the curve, corresponding to the point of diminishing returns.

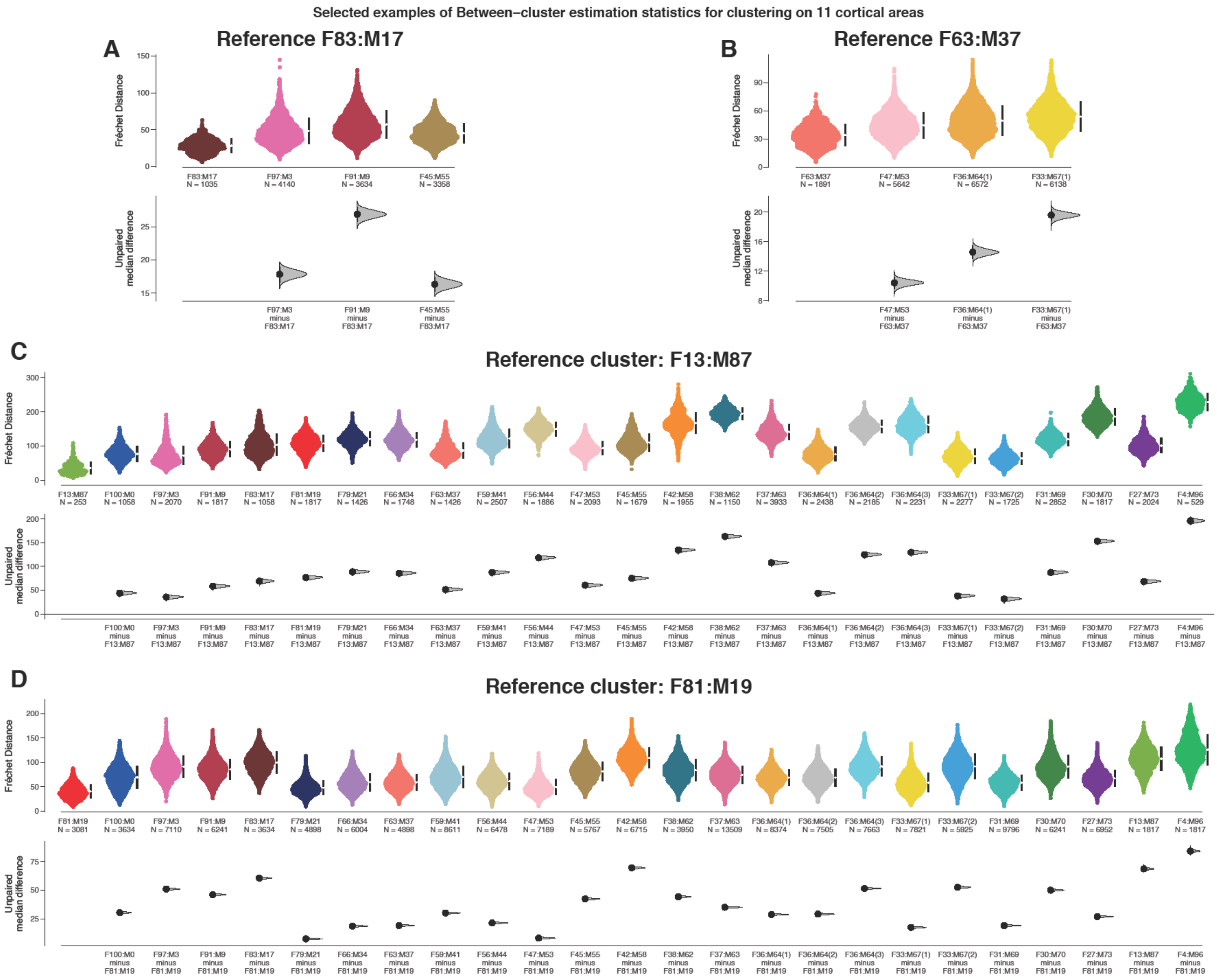


**Supplementary Figure 4** *Between cluster estimation statistics of developmental trajectories.* Fréchet distances were calculated between each of the 25 developmental trajectory clusters to quantify similarity in trajectory shape. (A–D) Swarm plots show the distributions of Fréchet distances and unpaired median differences for each cluster relative to a reference cluster (left-most in each plot). A) Comparison of four clusters to reference cluster F83:M17; B) four clusters to reference F63:M37; C) all 25 clusters to reference F13:M87; D) all 25 clusters to reference F81:M19. Dots indicate median differences, horizontal bars represent 95% confidence intervals and the grey shading represents the bootstrapped sampling distributions.

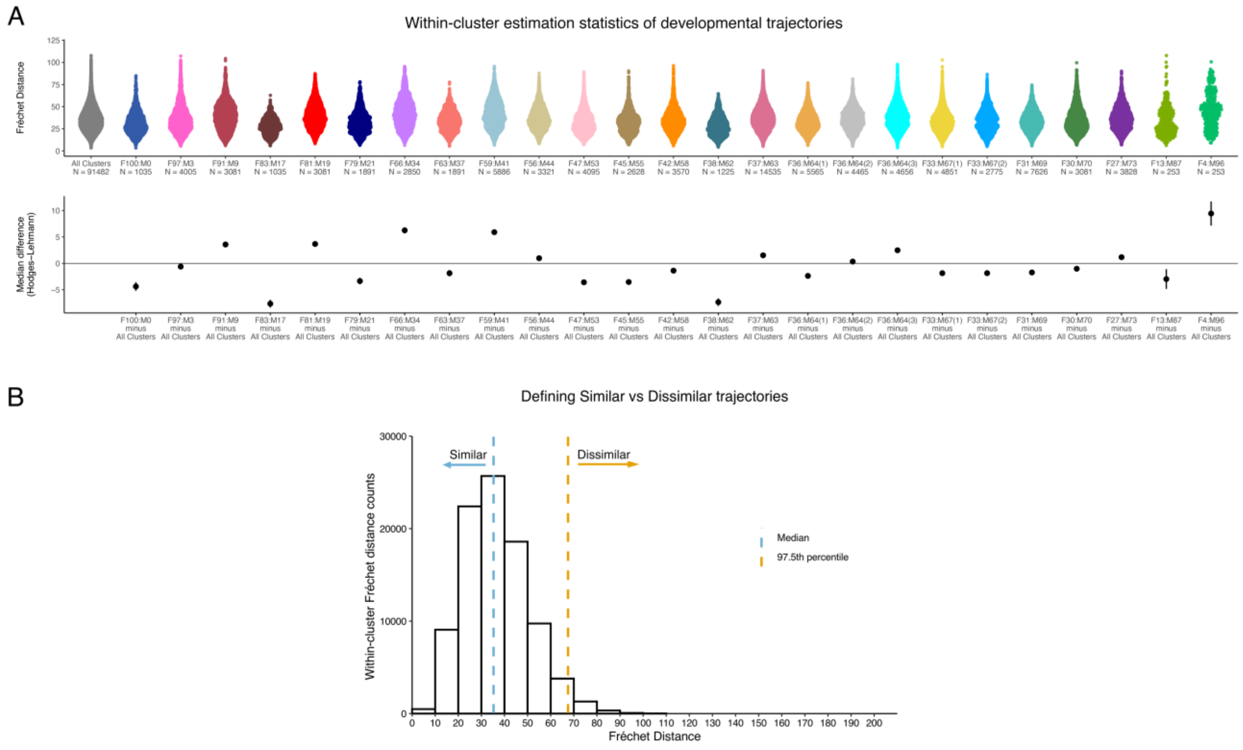


**Supplementary Figure 5** *Within-cluster estimation statistics of developmental trajectories.*

A) Swarm plots show the Fréchet distances within each cluster, and estimation plots display the median differences relative to the distribution of all within-cluster distances. Black dots indicate the Hodges–Lehmann median difference, and black bars represent 95% confidence intervals.

B) Distribution of within-cluster Fréchet distances, with the vertical lines indicating the median (blue) and the 97.5$^{th}$ percentile (orange). The median of this within-cluster Fréchet distance distribution was taken as the reference point for defining similarity. Values below the median are considered developmentally similar, values between the median and 97.5$^{th}$ percentile are within expected variation, and values greater than the 97.5$^{th}$ percentile are developmentally dissimilar.

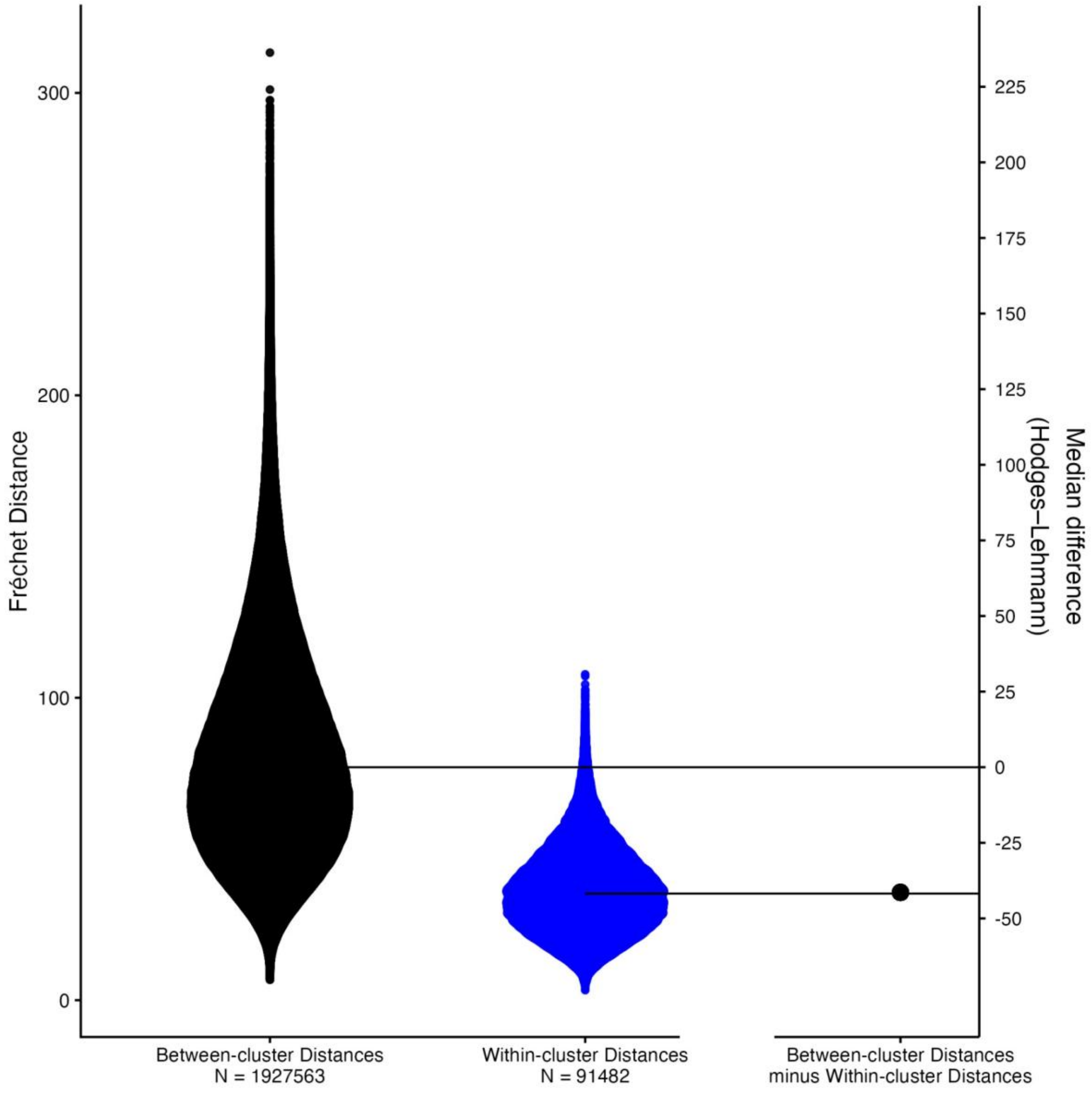


**Supplementary Figure 6** *Comparison of within- and between-cluster Fréchet distance distributions.* Distributions of Fréchet distances calculated within clusters and between clusters are shown, with the between-cluster distribution serving as the reference. The Hodges–Lehmann estimate of the median difference between distributions is indicated by a black dot, and horizontal lines denote the median of each distribution.

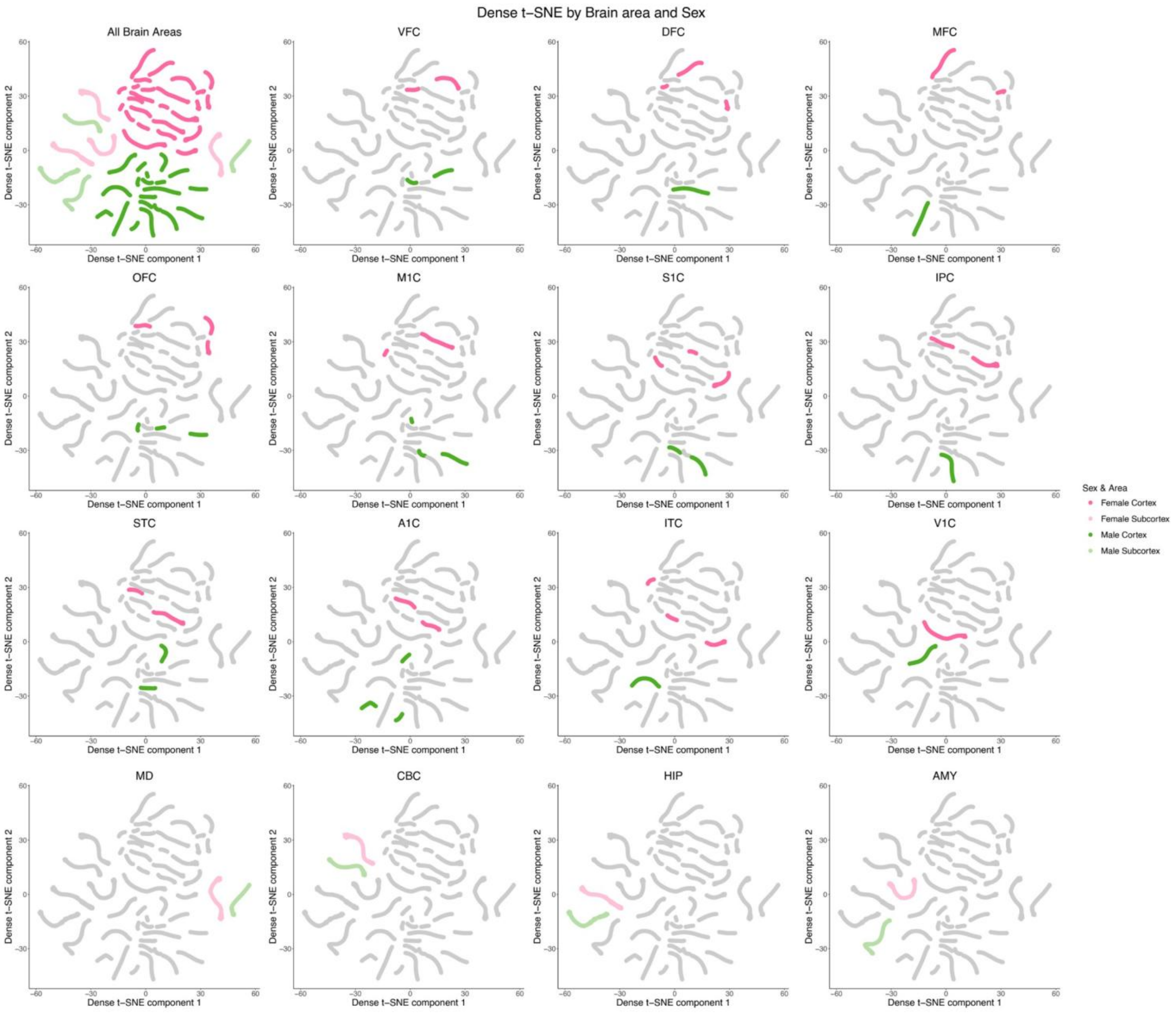


**Supplementary Figure 7** ***den-SNE by sex and brain area.*** Density-preserving t-SNE (den-SNE) projections of high-dimensional LOESS trajectory curves into two-dimensional space. Each curve represents a gene–brain area trajectory and is coloured by brain region and sex. Saturated pink and green denote female and male cortical areas, respectively, whereas pale pink and green denote female and male subcortical structures.

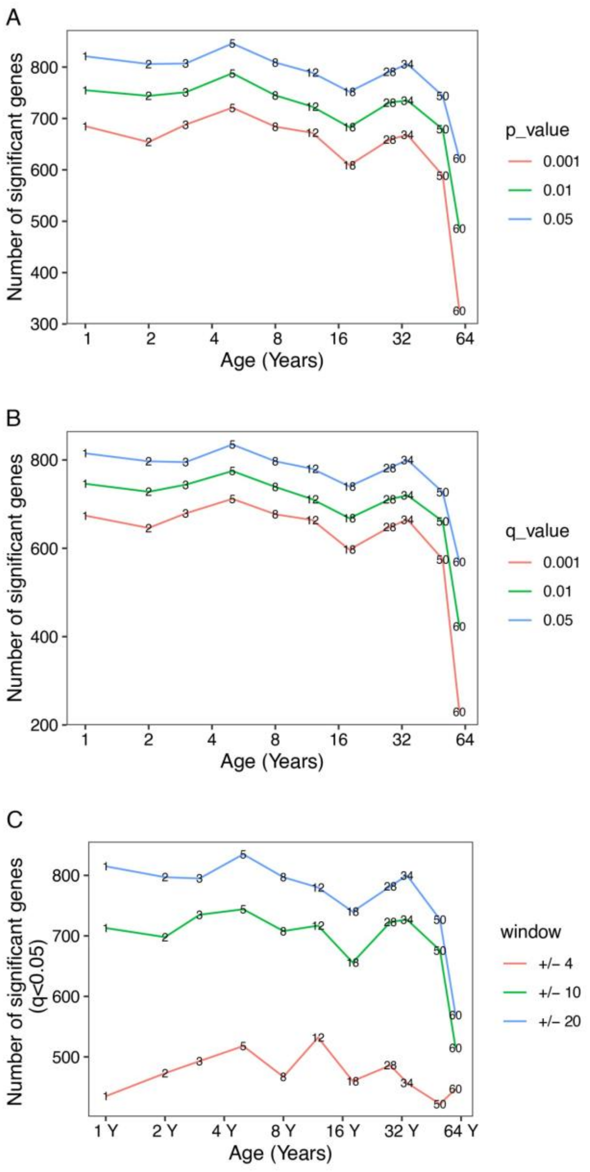


**Supplementary Figure 8** *Assessing robustness of DE-SWAN.* The number of significantly differentially expressed genes was quantified across age under different analysis parameters. A) Varying uncorrected p-value thresholds. B) Using q-values after Benjamini–Hochberg false discovery rate (FDR) correction. C) Varying sliding window sizes to assess robustness. Inflection points from these plots were used to define the window centers.

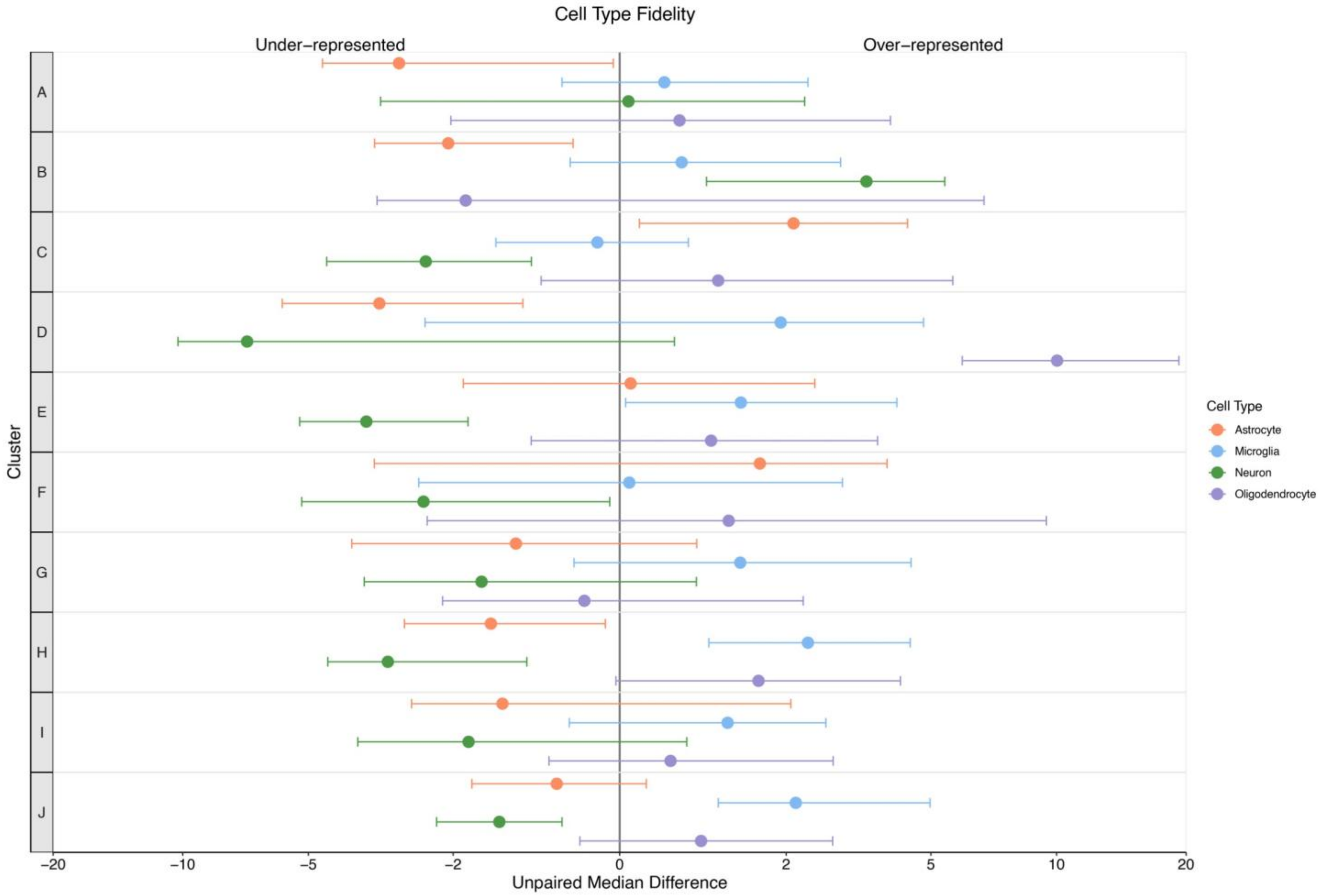


**Supplementary Figure 9** *Cell-type fidelity for DE-SWAN clusters*. Lollipop plot shows the unpaired median difference Fidelity score for clusters with under- or over-representation of a cell type. Dots represent median differences and horizontal lines indicate 95% confidence intervals.
